# Dynamic coupling between visual landmark processing in area 29e and parahippocampal path integration circuits of the rodent cortex

**DOI:** 10.1101/2025.11.11.687899

**Authors:** Gorkem Secer, Bharath Krishnan, Noah J. Cowan, James J. Knierim

## Abstract

External landmarks anchor internal spatial representations. The neural mechanisms underlying this process and the sources of landmark signals remain unknown. We recorded neurons across five parahippocampal regions while rats navigated in a virtual reality apparatus, inducing conflict between self-motion and landmark cues. Area 29e, a parahippocampal field putatively homologous to primate *area prostriata*, maintained firing coupled to landmarks even when other regions decoupled from the landmarks. This decoupling was preceded by a decline of gamma-band influence from 29e to MEC and a concomitant decline of theta-modulated feedback from MEC to 29e. Area 29e also displayed weak theta modulation, strong gamma rhythmicity, strong egocentric head-direction tuning, and enhanced landmark contrast sensitivity. These findings provide strong evidence that 29e serves as a specialized visuospatial hub that provides landmark signals for anchoring parahippocampal spatial representations to the external world.

## Main Text

Many higher-order neural systems display internal dynamics and represent explicit cognitive states in the absence of overt sensory stimulation. Sensory input does not drive a “blank slate” of quiescent neurons waiting for input about the external world. Rather, such input modulates a rich set of internally generated representations that perform important neural computations that are critical for the proper functioning of the system. Understanding the interplay between external sensory input and endogenous neural dynamics is key toward understanding the principles of neural computation. Spatial navigation provides an ideal model for dissecting this interplay. The hippocampal-parahippocampal system, coordinated by endogenous neural oscillations spanning multiple frequency bands (*1–5*), continuously updates a position signal on the animal’s cognitive map by integrating self-motion-generated cues over time—a process known as path integration (*6–9*). To prevent error from accumulating in this incremental estimate of the animal’s position, stable landmarks must be utilized to anchor the map to an external reference frame (*10–14*).

Although the neural correlates of path integration (e.g., grid cells and speed cells) are relatively well characterized in rodent hippocampal and parahippocampal areas (*9*, *15–25*), the neural pathways that convey visual landmark signals that calibrate and correct this internal computation remain unclear. Area 29e is a relatively understudied region interposed between the dorsal presubiculum/postsubiculum (PR) and the parasubiculum (PA). Recent work suggests that rodent area 29e is homologous with the primate area prostriata, which has been implicated in processing stimuli in the peripheral visual field (*26–28*).

Area 29e receives direct inputs from the dorsal lateral geniculate nucleus and V1 and projects to PR and medial entorhinal cortex (MEC; *29*–*32*), identifying it as a potential candidate relay of visual landmark signals to parahippocampal circuits in rodents. Using a virtual reality system, which allowed self-motion signals to be decoupled from visual landmarks (*12*, *33*, *34*), and a square open-field arena, we found that 29e contained a heterogeneous population of neurons displaying both spatial and visual firing correlates. One subpopulation had directionally tuned response curves and firing that remained tightly coupled to distal visual landmarks under conditions in which the firing fields of neurons in the other regions became decoupled from these landmarks. Area 29e showed a large proportion of egocentric head-direction neurons, consistent with retina-centered visual drive, and stronger sensitivity to landmark contrast compared to other regions. Furthermore, we found strong gamma-mediated feedforward communication from 29e to MEC and theta-mediated feedback communication from MEC to 29e.

Critically, this feedforward–feedback coupling became disrupted when the landmarks lost control of the MEC spatial firing. Our findings provide a physiological and functional characterization of area 29e in freely moving animals. They reveal the neural dynamics through which the parahippocampal system integrates visual landmark information with internal path-integration computations in a temporally coordinated loop between visual cortex and the brain’s cognitive mapping system.

## Results

### Area 29e showed spatially correlated activity that was more tightly bound to visual landmarks than neighboring parahippocampal areas

We performed simultaneous tetrode recordings in area 29e and four other parahippocampal areas (MEC, PA, PR, and postrhinal cortex [PO]; Fig. 1A and fig. S1-2) as rats (n = 5) navigated freely in our ‘Dome’ virtual-reality apparatus, which allowed for precise control of the locations and salience of visual landmarks during untethered physical locomotion (*12*, *33*; Fig. 1B). When the landmarks were stationary, a comparable proportion of 29e neurons displayed significant spatially correlated activity as in the other parahippocampal regions (29e: 70/78, MEC: 257/328, PA: 178/232, PO: 87/112, PR: 32/44; chi-squared test χ^2^ (4,796) = 7.64, p *=* 0.105). Many spatially correlated 29e neurons increased or decreased their firing rates selectively in the vicinity of these stationary landmarks (Fig. 1C). We defined two metrics to quantify the strength and between-landmark similarity of such peri-landmark firing-rate changes: (1) the *z-index* was computed as the mean absolute difference between z-scored firing rates at track locations closest to landmarks and those midway between landmarks and (2) the *local similarity index* was computed as the correlation of the firing rate profile of a neuron in the neighborhood of one landmark with its firing rate near a second landmark, averaged across all pairs of landmarks for a given neuron. Both metrics were significantly larger in spatially correlated neurons in 29e than those in other regions (Fig. 1D).

**Fig. 1.**
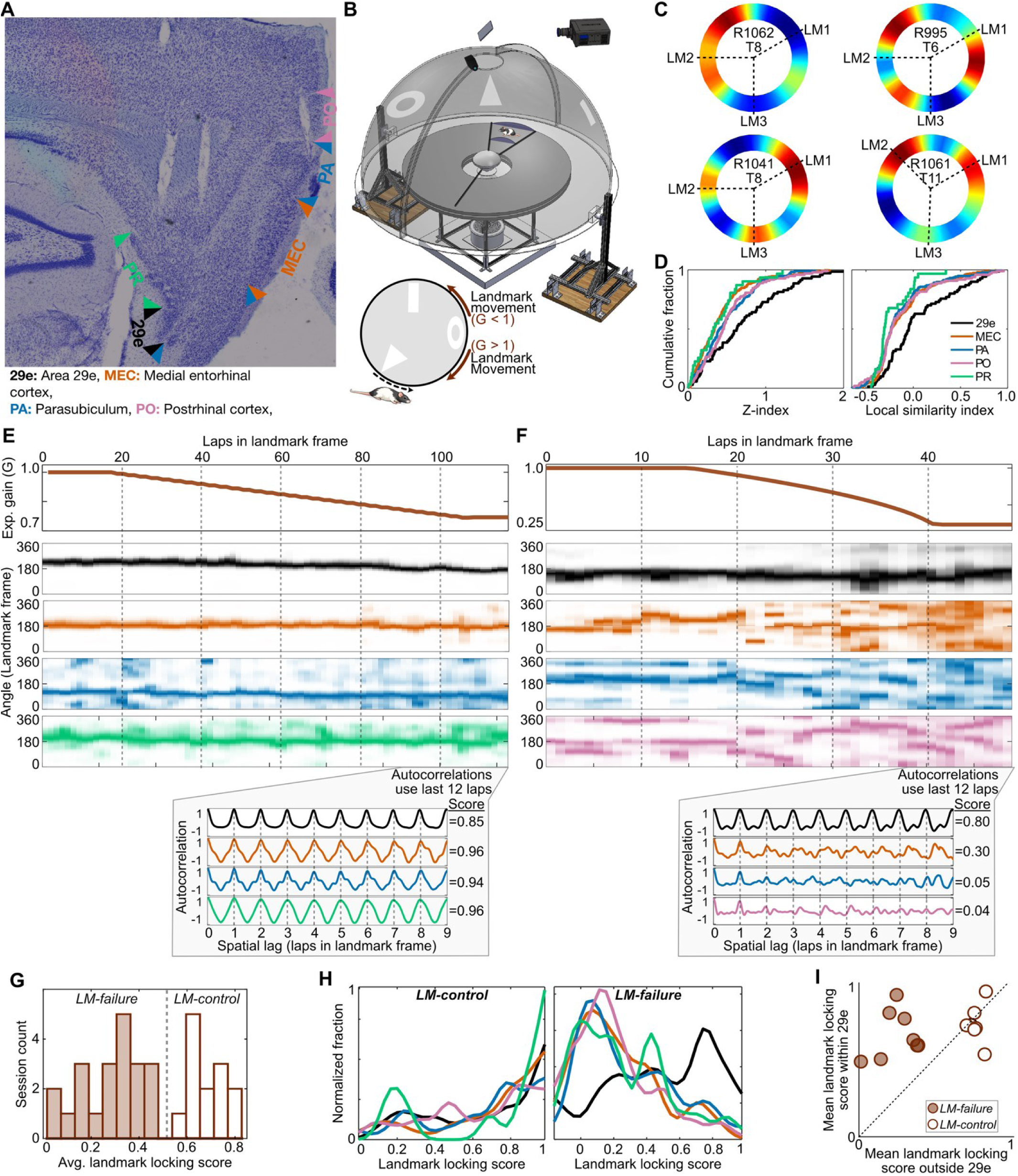
Landmark-anchored spatial tuning in 29e compared to spatial tuning in neighboring parahippocampal regions (MEC, PA, PO, and PR). **(A)** Representative histology section (Nissl-stained, sagittal) showing the recorded parahippocampal regions. Colored arrowheads demarcate the superficial-layer borders of each region. The same color scheme is used throughout all subsequent figures. See fig. S1-2 for sections showing actual recording locations within each region. **(B)** Schematic of the Dome virtual-reality apparatus and experimental gain manipulation. Top: A large opaque hemispherical shell (rendered semi-transparent to visualize the interior) displayed visual landmarks. Inside, the rat ran circular laps around the perimeter of a table. A rotating central pillar carried three radial boom arms: a front arm with a reward spout, a back arm that guided the recording tether, and a cleaning arm (hardware not shown). Short clear walls attached to the feeding and tether booms moved with the rat, maintaining a constant enclosure around it. Components that could otherwise provide angular cues either co-rotated with the animal or were circularly symmetric, making the projected landmarks the dominant polarizing cues for localization. Bottom left: Illustration of experimental gain manipulation, where visual landmarks were moved as a function of the rat’s movement, controlled by the experimental gain (G; see main text). **(C)** Circular rate maps of four example, spatially correlated 29e neurons, with red-to-blue color gradient indicating maximum-to-minimum firing rates. Within each rate map, rat ID and the tetrode number from which the neuron was recorded are shown. Dashed lines mark the angular positions of the centers of the three landmarks projected onto the Dome. The first row shows two neurons that decrease their firing near landmarks, while the second row shows two neurons that increase their firing near landmarks. **(D)** Cumulative distributions of z-index (left) and local-similarity index (right), which quantify the strength and uniformity of landmark selective firing, respectively. The z-index was significantly larger in 29e than other regions (29e median [IQR]: 0.60 [0.24, 1.02], MEC = 0.33 [0.16, 0.54], PA = 0.40 [0.22, 0.72], PO = 0.44 [0.21, 0.60], PR = 0.33 [0.14, 0.54]; Kruskal-Wallis test χ^2^(4) = 22.2, *p* < 0.001; post-hoc Wilcoxon rank-sum tests with Holm-Bonferroni correction 29e vs. MEC: *Z* = 4.18, *p <* 0.001; 29e vs. PA: *Z =* 2.70, *p =* 0.013; 29e vs. PO: *Z =* 2.38, *p =* 0.017; 29e vs. PR: *Z =* 2.87, *p =* 0.012). The local-similarity index was also larger in 29e than other regions (29e median [IQR]: 0.05 [–0.24, 0.28], MEC = –0.23 [–0.31, 0.00], PA = –0.23 [–0.32, –0.02], PO = –0.23 [–0.30, –0.01], PR = –0.30 [–0.33, –0.06]; Kruskal-Wallis test χ^2^(4) = 22.3, *p* < 0.001; post-hoc Wilcoxon rank-sum tests with Holm-Bonferroni correction 29e vs. MEC: *Z* = 3.68, *p <* 0.001; 29e vs. PA: *Z =* 4.06, *p <* 0.001; 29e vs. PO: *Z =* 3.14, *p <* 0.001; 29e vs. PR: *Z =* 3.75, *p <* 0.001). **(E)** Representative closed-loop gain manipulation session showing landmark control across all regions. The top row shows the experimental gain as a function of laps in the landmark frame (x-axis). The session began with a gain of 1 (i.e., stationary landmarks). The gain was then gradually ramped to its target value, during which the landmark array rotated at increasing speeds opposite to the animal’s movement direction. The final gain was held constant for the duration of the session. The bottom row shows the lap-to-lap (with respect to landmarks) firing rate as a function of the animal’s angle in the landmark frame (y-axis) from example neurons, color-coded by region (as in panel A). The intensities were normalized to the maximum firing rate of the given neuron. In this session, example neurons from 29e (black), MEC (orange), PA (blue) and PR (green) illustrate that the firing locations of all spatially correlated neurons remained anchored to the rotating landmarks throughout the session (i.e., they fired at a consistent angle on the track [y-axis] across laps). Inset shows the spatial autocorrelograms of each representative neuron’s activity in the rotating landmark frame, calculated using the last 12 laps for lags from 0 to 9 laps (ensuring a minimum of three lap-samples per lag for reliable correlation estimates). Note the lap-to-lap periodicity in the autocorrelation traces for these neurons. **(F)** Representative session where most spatially correlated neurons—except in 29e—lost their anchoring during the gain ramp (same format as panel E). After about lap 20, the firing fields of the MEC (orange), PA (blue), and PO (pink) neurons became increasingly diffuse. The 29e neuron (black), by contrast, maintained stable firing in the landmark frame throughout the session. Inset shows the spatial autocorrelograms (calculated as in panel E) of each neuron’s activity in the rotating landmark frame. Note that the lap-to-lap periodicity for the neurons that lost landmark anchoring showed disrupted autocorrelation structures: either no clear periodicity (e.g., the orange neuron) or a non-unity periodicity (e.g., the blue and pink neurons), reflecting the continued firing of the neuron at repeated intervals different than a single, landmark-based lap. **(G)** Session-level bimodality of neuronal populations’ average landmark-locking scores. Sessions were classified as *LM-failure* (shaded bars) *or LM-control* (unshaded bars). **(H)** Distributions of landmark-locking scores for individual spatially correlated neurons. *LM-control sessions* (left) show statistically comparable landmark-locking scores across all regions (median [IQR]: 29e = 0.73 [0.39, 0.90], MEC = 0.81 [0.52, 0.92], PA = 0.76 [0.46, 0.88], PO = 0.66 [0.39, 0.85], PR = 0.89 [0.23, 0.97]; Kruskal-Wallis test χ^2^(4) = 4.2, *p* = 0.376). By contrast, *LM-failure sessions* (right) show significantly larger scores in 29e than other regions (median [IQR]: 29e = 0.50 [0.25, 0.77], MEC = 0.19 [0.06, 0.41], PA = 0.22 [0.07, 0.50], PO = 0.15 [0.03, 0.34], PR = 0.20 [0.04, 0.44]; Kruskal-Wallis test χ^2^(4) = 31.4, *p* < 0.001; post-hoc Wilcoxon rank-sum tests with Holm-Bonferroni correction 29e vs. MEC: *Z* = 5.13, *p <* 0.001; 29e vs. PA: *Z =* 4.11, *p <* 0.001; 29e vs. PO: *Z =* 4.74, *p <* 0.001; 29e vs. PR: *Z =* 3.08, *p <* 0.001). **(I)** Mean landmark-locking scores of spatially correlated 29e neurons within a session (y-axis) versus those of their simultaneously recorded counterparts in other regions (x-axis) within the same session. In *LM-control* sessions (n = 7), mean landmark-locking scores did not significantly differ between 29e (mean ± s.e.m. = 0.62 ± 0.04) and other regions (0.66 ± 0.02), whereas in *LM-failure sessions* (n = 8), 29e (0.56 ± 0.05) showed significantly higher mean landmark-locking scores than other regions (0.21 ± 0.03). In *LM-control sessions* (n = 7; unfilled circles), the normalized difference between scores of neurons inside 29e vs. outside 29e (i.e., (inside – outside)/(inside + outside)) was not significantly different than zero (one-sample t-test *t*(*6*) = 0.964, *p* = 0.372), whereas in *LM-failure* sessions (filled circles), the normalized difference was significantly positive (one-sample t-test *t*(*7*) = 6.93, *p* < 0.001), indicating greater anchoring to landmarks in 29e than in other regions. *LM-failure* sessions showed significantly larger normalized differences than *LM-control* sessions (t-test *LM-failure vs. LM-control*: *t*(13) = 5.7, *p* < 0.001).

Considering this modulation of 29e neurons’ spatially correlated firing near landmarks, we next tested the extent to which their firing fields were dynamically controlled by landmarks when the entire landmark constellation was rotated around the track in a behaviorally closed-loop fashion. The rotation of landmarks was controlled by an experiment gain (G; Fig. 1B) that set the ratio between the rat’s travel distance in the landmark frame of reference to the travel distance in the laboratory frame of reference (*12*). When G < 1, the landmark array was rotated around the center of the track in the same direction as the rat, providing the rat with the illusion that it was moving slower than it really was; when G > 1, it was rotated in the opposite direction, providing the rat with the illusion that it was moving faster than it really was. In each session, the value of G was initialized at 1 (stationary landmarks), gradually ramped to its final value, and then held constant at that final gain value for the remainder of the session.

Under these manipulations, spatially correlated neurons across all regions often maintained stable firing fields in the rotating landmark frame of reference across laps—a manifestation of landmark control over the internal representations of space in the Dome (Fig. 1E and fig. S3A), as previously demonstrated for CA1 place cells (*12*) and anterodorsal thalamic nuclei head direction cells (*35*). This lap-to-lap stability, however, was disrupted in some sessions, typically during the gain-ramp epoch and persisting until the end of the experiment (See Fig. 1F and fig. S3B for example sessions). To quantify the lap-to-lap stability of each neuron’s firing fields in the rotating landmark frame of reference, we defined a landmark-locking score based on the spatial autocorrelogram (up to a 9-lap lag) of the neuron’s activity during the last 12 laps of each session (see Methods). A landmark-locking score close to 1 indicates highly stable lap-to-lap firing of a neuron in the landmark frame (e.g., Fig. 1E bottom inset, all regions), whereas a score near 0 indicates a lack of this stability (e.g., Fig. 1F bottom inset, non-29e regions).

Averaging landmark-locking scores across all spatially correlated neurons across all regions in each session (number of neurons recorded per session: mean ± s.d. = 33.0 ± 17.2) revealed a bimodal distribution (Fig. 1G). An unsupervised clustering algorithm identified two clusters within this distribution. We refer to the cluster of 13 sessions with higher landmark-locking scores (mean ± s.d. = 0.68 ± 0.08) as *LM-control* sessions, indicating reliable control of the landmarks over firing fields of spatially correlated neurons from all regions (as in Fig. 1E). By contrast, the remaining cluster of 21 sessions with lower landmark-locking scores (0.27 ± 0.13) are termed *LM-failure* sessions, indicating the failure of this landmark control over most spatially correlated neurons, particularly those outside 29e (as in Fig. 1F).

Area 29e neurons were more robustly controlled by the landmarks than the other parahippocampal regions. In *LM-control sessions*, landmark-locking scores of spatially correlated neurons were strongly skewed towards 1 in all regions (Fig. 1H, left). In *LM-failure sessions*, however, landmark-locking scores of spatially correlated neurons in MEC, PA, PO, and PR were skewed towards 0, while those in 29e remained skewed toward 1, indicating greater anchoring of 29e neurons to landmarks (Fig. 1H, right). Comparing mean landmark-locking scores between simultaneously recorded neurons from 29e and other regions ensured that these regional differences were not due to session-specific effects or non-uniform sampling (Fig. 1I). This greater landmark-anchoring by 29e neurons persisted even when landmark rotation was independent of the rat’s movement during open-loop gain manipulations (fig. S4; see Methods). Collectively, these data suggest that, compared to neighboring parahippocampal regions specialized for spatial processing with greater reliance on path integration, area 29e may be a visuospatial region that codes for spatial representations anchored to visual landmarks.

### Area 29e was sensitive to sensory aspects of visual landmarks

We next investigated the sensitivity of area 29e to sensory aspects of the landmarks in comparison to neighboring parahippocampal areas. We first tested the immediate effect of turning off the landmarks following the gain manipulations (i.e., immediately after Epoch 3). Because the effects of extinguishing the landmarks would be confounded at the single-unit level by where the rat was located when the landmarks were turned off (i.e., whether the rat was in a spatial firing field of a particular cell), we quantified the responses to landmark removal using multi-unit firing rates in five animals. For each tetrode, we computed a landmark-removal response score as the absolute normalized change in its firing rate from the 600 ms period immediately preceding landmark removal to the 400 ms immediately following it (see Methods). Larger positive scores indicate stronger firing-rate changes, regardless of the direction, whereas scores near zero indicate little or no change. Only tetrodes in 29e and PO showed a large majority of positive landmark-removal response scores, with 29e exhibiting the largest scores on average (Fig. 2A). A linear mixed-effects model accounting for dependencies among tetrodes from the same animal and brain region revealed that 29e showed significantly higher scores than MEC and PA, but it was not significantly different from PR or PO.

**Fig. 2.**
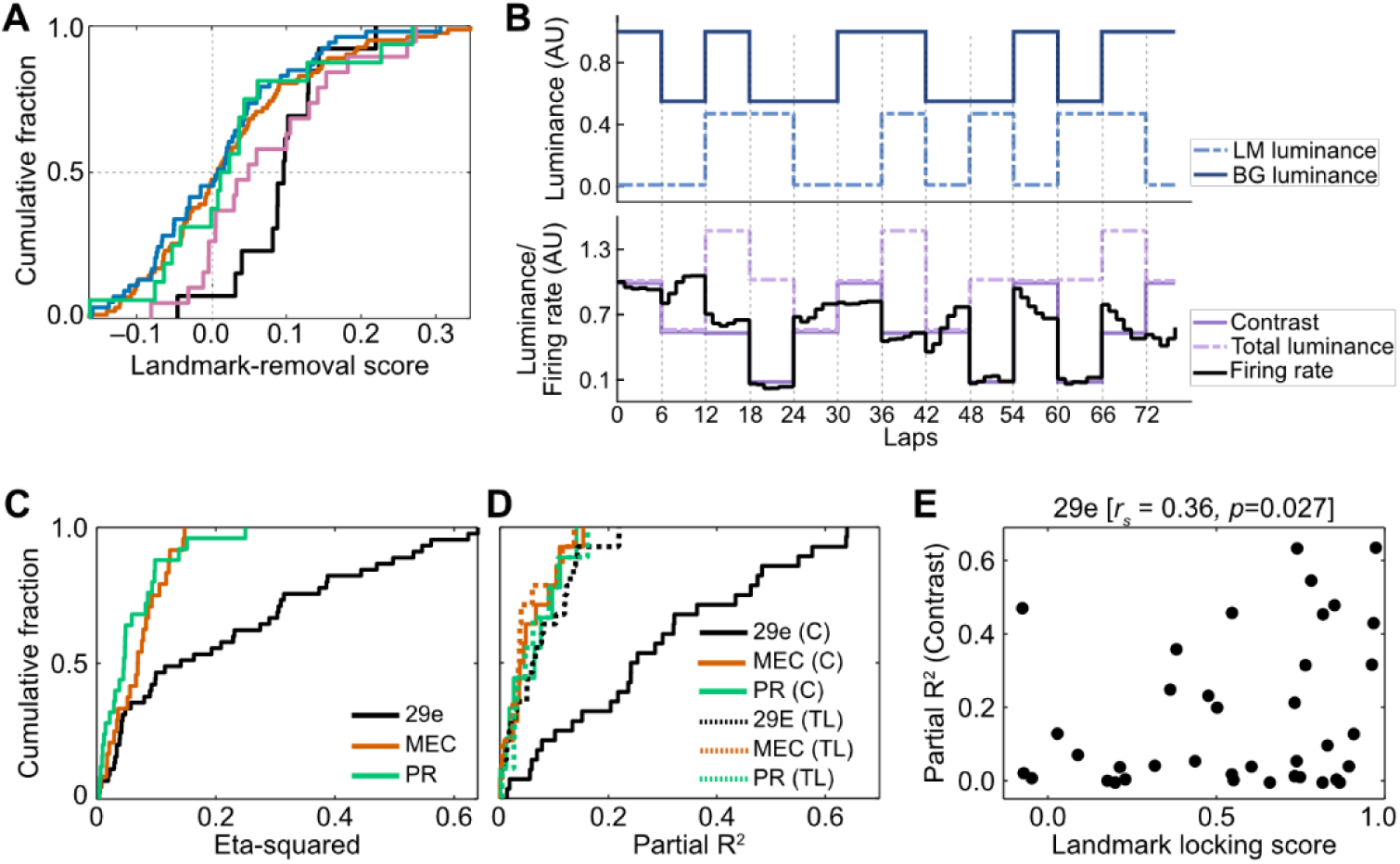
Greater sensitivity of area 29e to sensory aspects of visual landmarks compared to neighboring parahippocampal areas. **(A)** Cumulative distributions of landmark-removal response scores from unique tetrode locations (29e: n = 13, MEC: n = 87, PA: n = 54, PO: n = 19, PR: n = 16; colors as in Panel A). Response scores of tetrodes were significantly positive in 29e (median [IQR] = 0.096 [0.071, 0.129]; Wilcoxon signed-rank test *Z* = 2.97, *p =* 0.003) and PO (0.049 [–0.001, 0.139]; *Z* = 2.77, *p =* 0.005) but not in MEC (0.008 [–0.054, 0.082]; *Z* = 1.35, *p =* 0.176), PA (0.009 [–0.068, 0.061]; *Z* = 0.20, *p =* 0.842), or PR (0.018 [–0.047, 0.053]; *Z* = 0.72, *p =* 0.469). Examining distribution of scores with a linear mixed-effects model showed significant differences across regions (*F*(165.2) = 3.24, *p* = 0.014; see Methods for random-effects structure of the model and further details). The model also showed that 29e exhibited significantly higher scores than MEC (*t*(151.3) = 2.4, *p* = 0.018) and PA (*t*(138.0) = 3.2, *p* = 0.002), but not PR (*t* (153.3) = 1.22, *p* = 0.222) or PO (*t*(175.6) = 1.56, *p* = 0.119). **(B)** Protocol for luminance manipulation experiments and an example of a contrast-responsive neuron. Top: Experiment design. Bottom: Example neuron from area 29e showing peak lapwise firing rate changes (black) that more closely followed changes in landmark contrast (C = LM - BG, solid purple) rather than total luminance (TL = LM + BG, dashed purple). Firing rates are normalized to the neuron’s peak firing rate in the first lap of the session. **(C)** Region-wise cumulative distributions of total η^2^ values from two-way ANOVA with background and landmark luminance as factors. Total η^2^ was significantly larger in 29e neurons than in MEC and PR neurons (median [IQR] 29e = 0.14 [0.04, 0.32], MEC = 0.06 [0.03, 0.09], PR = 0.04 [0.01, 0.08]; Kruskal-Wallis test χ^2^(2) = 13.95, *p* < 0.001; post-hoc Wilcoxon rank-sum tests with Holm-Bonferroni correction 29e vs. MEC: *Z* = 2.61, *p =* 0.008; 29e vs. PR: *Z =* 3.30, *p* = 0.001). **(D)** Cumulative distributions of partial R^2^ values from linear models predicting neurons’ lapwise peak firing rates using landmark contrast (C, solid lines) and total brightness (TL, dashed lines) as predictors. Two-way ANOVA on sqrt-transformed partial R^2^ values revealed significant main effects of region (*F*(2,96) = 16.3, *p* < 0.001) and predictor type (*F*(1,96) = 7.5, *p* = 0.007), as well as a significant interaction (*F*(2,96) = 9.3, *p* < 0.001). Post-hoc tests (with Holm-Bonferroni corrections) showed that partial R^2^ for landmark contrast in 29e (mean ± s.e.m. = 0.28 ± 0.03) was significantly greater than landmark-contrast partial R^2^ in MEC (0.05 ± 0.01) and PR (0.06 ± 0.01), and total-brightness partial R^2^ in all recorded regions (29e: 0.07 ± 0.01; MEC: 0.04 ± 0.01; PR: 0.06 ± 0.01); all pairwise comparisons with landmark contrast in 29e: *p* < 0.001. **(E)** Significantly positive correlation between the partial R^2^ (y-axis) for landmark-contrast tuning of 29e neurons with spatially and their landmark-locking scores (x-axis; Pearson’s r = 0.36, *p* = 0.027; landmark-locking scores were normalized with the Fisher-z transform). Each dot corresponds to one neuron.

To determine whether individual 29e neurons were tuned to the luminance or contrast of the landmarks, we performed additional experiments in a subset of animals (n = 2), recording from 29e, MEC, and PR (no tetrodes were in PA or PO in these animals). As rats ran laps, we pseudo-randomly varied landmark and background luminance between two levels (i.e., high or low) across six-lap blocks, with the experimental gain fixed at 1 to keep landmark locations stationary (Fig. 2B). Similar proportions of neurons with significant luminance sensitivity were identified across all recorded regions (29e: 28/45; MEC: 14/24; PR: 9/25; chi-squared test χ^2^(2,94) = 4.66, p *=* 0.096). However, the magnitude of luminance sensitivity was significantly larger in neurons from 29e than those from MEC and PR (Fig. 2C). This heightened effect of landmark and background luminance on 29e single-neuron activity may reflect their tuning to the total luminance in the environment or to the contrast between the landmark and background luminance. For each neuron with significant landmark-or background-luminance sensitivity, we fit a linear model to the peak firing rates on each lap, using landmark contrast and total luminance as predictors. Two-way ANOVA (region × predictor) on the partial R² values—which quantify the firing-rate variance uniquely explained by each predictor—revealed particularly pronounced tuning of 29e neurons to landmark contrast, the strength of which significantly exceeded not only contrast tuning in other regions but also total luminance tuning in any region (Fig. 2D).

Following the luminance-change epochs, we performed closed-loop gain manipulations with large gains to induce landmark-control failure. Spatially correlated neurons of 29e showed a significant positive relationship between the strengths of their landmark-contrast tuning and their landmark anchoring (Fig. 2E; fig. S5).

### Area 29e demonstrated tuning to egocentric and allocentric head direction, with egocentric tuning more closely associated with sensory-like responses to landmarks

To further characterize spatially correlated activity in 29e in comparison to neighboring parahippocampal areas, we analyzed single-unit activity obtained as the rats (n = 5) foraged in an open arena after they completed each Dome session (fig. S6A). The arena was in a separate room from the Dome. We examined allocentric tuning (Fig. 3A; fig. S6B) by fitting a generalized linear model (GLM) that jointly captures both place and head-direction (HD) tuning of a neuron, thereby minimizing biases due to behavioral confounds such as uneven sampling of HDs in different locations (*36*, *37*). From the GLM-derived 2D positional rate map and allocentric HD tuning curve of each neuron, we then calculated the Skaggs’ spatial information (SI) score (*38*) and the Rayleigh score (*39*, *40*), respectively. SI scores were significantly lower in area 29e than other regions except PR (Fig. 3B top row). By contrast, Rayleigh scores of allocentric HD tuning curves were significantly larger in area 29e than other regions (Fig. 3B middle row).

**Fig. 3.**
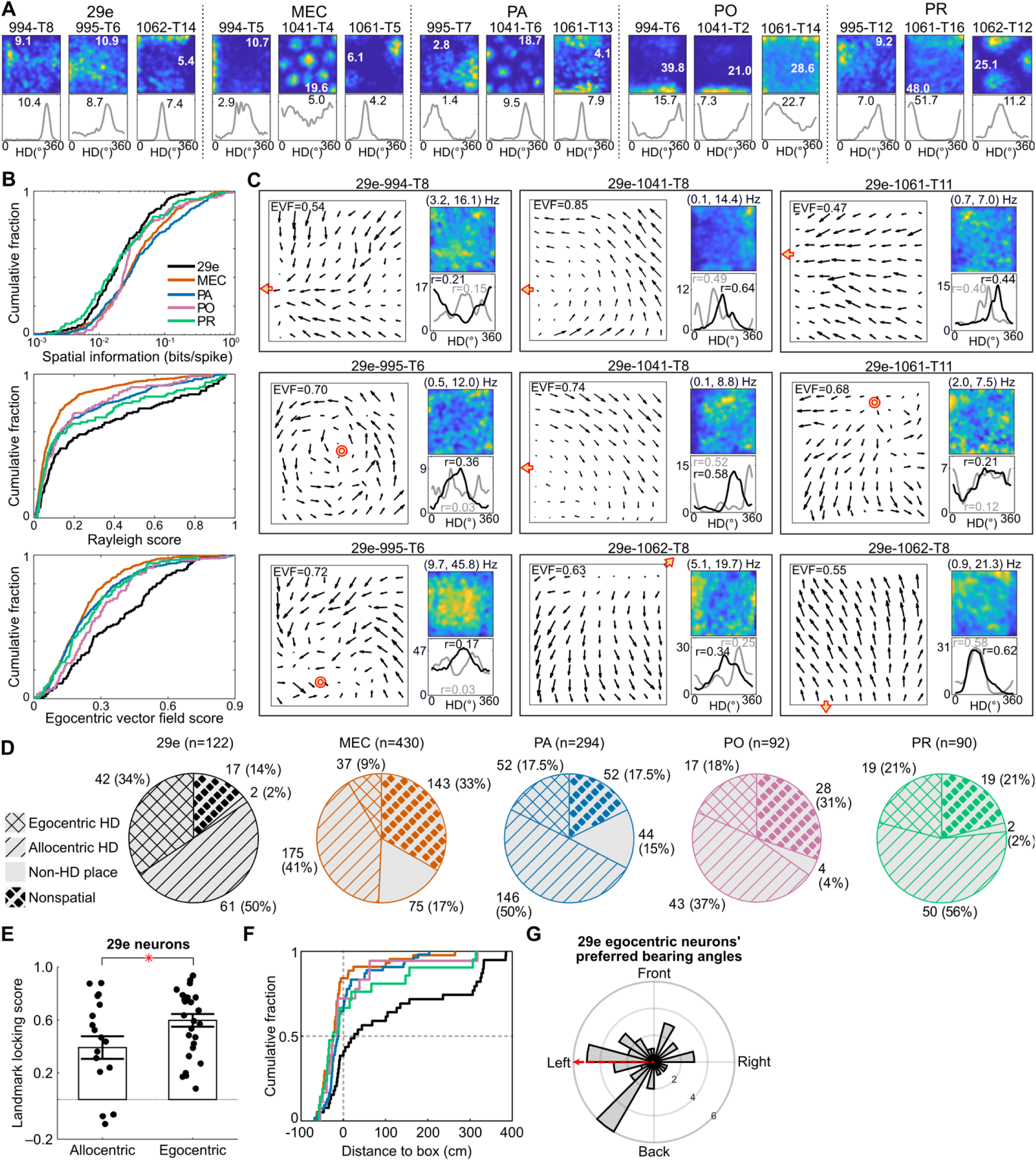
Two-dimensional spatial representations of Area 29e neurons compared to neurons in neighboring parahippocampal regions (MEC, PA, PO, and PR). **(A)** Allocentric tuning of three example neurons from each region. For each neuron, a 2D positional rate map (top) and a 1D allocentric head-direction (HD) tuning curve (bottom) are shown. Rat ID and tetrode number for each neuron are shown above its rate map. The number inset in each plot denotes the peak-firing rate. The y-axis of each HD plot starts at zero. **(B)** Region-wise cumulative distributions of SI score (top), Rayleigh score (middle), and egocentric vector field (EVF) score (bottom). All scores differed significantly across regions: SI scores were significantly lower in 29e (median [IQR] = 0.019 [0.010, 0.045]) than MEC (0.034 [0.016, 0.083]), PA (0.036 [0.016, 0.120]), and PO (0.027 [0.018, 0.054]), but not PR (0.019 [0.009, 0.051]); Kruskal-Wallis omnibus test: χ^2^(4) = 38.25, *p* < 0.001; post-hoc Wilcoxon rank-sum test with Holm-Bonferroni correction 29e vs. MEC*: Z* = –4.6, *p <* 0.001; 29e vs. PA: *Z* = –4.8, *p* < 0.001; 29e vs. PO: *Z* = −2.8, *p* = 0.009; 29e vs. PR: *Z* = 0.0, *p* = 0.961.Rayleigh scores were significantly higher in 29e (median [IQR] = 0.165 [0.064, 0.521]) than other regions (MEC = 0.066 [0.033, 0.134]), PA = 0.096 [0.045, 0.295], PO = 0.099 [0.045, 0.264], PR = 0.085 [0.043, 0.396]); Kruskal-Wallis omnibus test χ^2^(4) = 54.37, *p* < 0.001; post-hoc Wilcoxon rank-sum test with Holm-Bonferroni correction 29e vs. MEC*: Z* = 6.6, *p <* 0.001; 29e vs. PA: *Z* = 3.3, *p* < 0.001; 29e vs. PO: *Z* = 2.5, *p* = 0.022; 29e vs. PR: *Z* = 2.1, *p* = 0.035.Similar to Rayleigh scores, EVF scores were also significantly higher in 29e (median [IQR] = 0.336 [0.195, 0.516]) than other regions (MEC = 0.193 [0.116, 0.282], PA = 0.197 [0.120, 0.338], PO = 0.254 [0.163, 0.395], PR = 0.225 [0.123, 0.353]); Kruskal-Wallis omnibus test χ^2^(4) = 55.73, *p* < 0.001; post-hoc Wilcoxon rank-sum test with Holm-Bonferroni correction 29e vs. MEC*: Z* = 7.1, *p <* 0.001; 29e vs. PA: *Z* = 5.5, *p* < 0.001; 29e vs. PO: *Z* = 2.7, *p* = 0.006; 29e vs. PR: *Z* = 3.9, *p* < 0.001. **(C)** Egocentric tuning of example 29e neurons. For each neuron, the left panel shows the HD vector field, with the EVF score indicated. Arrow direction depicts the neuron’s preferred HD within a 15 × 15 cm positional bin centered at the arrow base; arrow size represents the product of the neuron’s total firing rate (summed across head-direction bins) and its mean resultant length inside the same positional bin (see Methods). Within each HD vector field, red/yellow circles mark anchor locations inside the arena, while red/yellow arrowheads point to anchors outside the arena boundaries. (Note that the EVF scores are independent of the anchor locations.) On the right, the top panel shows the allocentric 2D positional rate map and the bottom panel shows the HD tuning curves in allocentric (gray) and egocentric (black) reference frames. The color scale (blue to yellow) represents firing rate from minimum to maximum, which are displayed above each rate map as (min, max) Hz. Numbers in the HD plots denote the mean resultant length of HD tuning in each frame. **(D)** Pie charts showing the proportions of egocentric HD-tuned, allocentric HD-tuned, non-directional place-tuned neurons, and non-spatial neurons. Region IDs and the total neuron counts are given above each chart. There were significant regional differences in the prevalence of spatial tuning types (Chi-squared test χ^2^(12,1028) = 107.55, p *<* 0.001). Post-hoc Fisher’s exact test with Holm-Bonferroni correction showed that egocentric HD-tuned neurons were significantly more prevalent in 29e than in MEC (*p* < 0.001), PA (*p* < 0.001), and PO (*p* = 0.039) but not PR (*p* = 0.182). Nondirectional place-tuned neurons were less prevalent in 29e than in MEC (*p* < 0.001) and PA (*p* < 0.001), but not PO (*p* = 0.812) or PR (*p* = 1.000). The prevalence of allocentric HD-tuned neurons in 29e was similar across regions (all post-hoc pairwise *p >* 0.075). **(E)** Significant difference in the landmark-locking scores between 29e allocentric and egocentric neurons (represented as dots) during *LM-failure* sessions in the Dome. Bars show population means (allocentric: 0.39; egocentric: 0.60), and error bars indicate s.e.m. (allocentric: 0.09; egocentric: 0.05); t-test allocentric vs. egocentric: *t(44) = –*2.5, *p =* 0.030. **(F)** Cumulative distributions of the signed distances from the estimated anchor points of egocentric neurons to the open-arena border. We first tested, for each region, whether its signed distances were significantly different from zero. Area 29e showed significantly positive median distance (median [IQR] = 26 [–14, 289] cm; Wilcoxon’s signed-rank test with Holm-Bonferroni correction: *p* = 0.016). By contrast, the medians were negative in other regions (MEC = –24 [–45, –11], *p* = 0.005; PA = –13 [–28, 8], *p* = 0.186; PO = –20 [–39, 28], p = 0.892; PR = –24 [–40, 31], p = 0.892), Direct comparison between 29e and other regions showed that the signed distance was significantly larger in 29e than other regions (Kruskal-Wallis test χ^2^(4) = 22.55, *p* < 0.001; post-hoc Wilcoxon rank-sum test with Holm-Bonferroni correction 29e vs. MEC*: Z* = 4.4, *p <* 0.001; 29e vs. PA: *Z* = 3.2, *p* = 0.004; 29e vs. PO: *Z* = 2.8, *p* = 0.011; 29e vs. PR: *Z* = 2.5, *p* = 0.012). **(G)** Distribution of preferred bearing angles for egocentric neurons in area 29e. Radial axis indicates the number of neurons. Significantly more neurons had anchor points on the animal’s left (n = 28) than right side (n = 11; binomial test with a null probability of 0.5, p = 0.009). Furthermore, the overall circular distribution was significantly non-uniform (Rayleigh test: z = 4.02 *p* = 0.017), skewed toward the left-rear quadrant as indicated by the circular mean direction (red arrow).

Classic, allocentric HD cells fire maximally in a consistent direction regardless of the animal’s location, yielding parallel preferred HD vectors across the arena (*37*, *41*; fig. S7A). In contrast, egocentric HD cells demonstrate progressive, location-dependent shifts in their preferred HD, forming vector fields with rotational, radial (outward/inward-pointing), or hybrid patterns (*42–45*), consistent with tuning for the egocentric bearing of the head relative to an external anchor point. For example, a cell that fires whenever a specific location is to the right of the animal’s head, regardless of the animal’s allocentric position, would show a vector field with a clockwise rotational flow about that location, and a cell that fires whenever that location is directly ahead of the rat would show a centripetal, radial flow. Many 29e neurons displayed preferred HD-vector fields consistent with egocentric HD tuning (Fig. 3C; see also fig. S7B-C for more examples). To quantify egocentric HD tuning, we developed an egocentric vector-field (EVF) score that measures the consistency of rotational and radial egocentric patterns within a neuron’s HD vector field across the arena, without relying on an exhaustive search to identify an optimal anchor point in the environment, as was done in previous studies (*43*, *46–48*) and which introduces biases in comparing egocentric vs. allocentric tuning curves (*43*; fig. S8; see Methods). The EVF score ranges from 0 (indicating no egocentric tuning, such as when preferred HD vectors are randomly oriented or parallel everywhere) to 1 (indicating maximal egocentric tuning). Across all neurons, the EVF scores had significantly higher values in 29e than other regions (Fig. 3B bottom row).

We used a combination of EVF, Rayleigh, and SI scores, together with a within-session tuning stability requirement, to classify each cell as an allocentric HD cell; an egocentric HD cell; a nondirectional, spatially correlated cell; or a nonspatial cell (see Methods). Egocentric HD-tuned neurons were significantly more prevalent in 29e than other regions (Fig. 3D). Nondirectional place-tuned neurons were rare in 29e, at a proportion significantly lower than in MEC and PA, but not PO or PR. By contrast, the prevalence of allocentric HD-tuned neurons was similar across regions.

Because visual sensory cues are initially encoded in egocentric (i.e., retinocentric) frames before being converted to an allocentric frame in higher-order areas, we hypothesized that egocentric HD-tuned neurons would be more closely associated with encoding visual landmarks than allocentric ones. Indeed, compared to 29e neurons classified as allocentric in the open arena, egocentric neurons exhibited, in the Dome, significantly higher landmark-locking scores (Fig. 3E; fig S9A) and a trend toward greater sensitivity to landmark contrast (fig. S9B).

Given that egocentric 29e neurons exhibited properties consistent with visual-landmark coding, we hypothesized that these neurons would be anchored to locations outside the open arena, where salient landmarks were located. We computed for each egocentric neuron the shortest signed distance from the arena boundary to its optimal anchor point estimated by maximizing the negative log-likelihood of a GLM. A negative distance indicates an anchor point inside the arena, while a positive distance indicates one outside. Median signed distance was significantly positive in 29e, indicating anchoring to locations beyond the arena boundary. By contrast, the medians were negative in other regions, with only MEC reaching statistical significance. Cross-regional comparison of signed distance distributions confirmed that anchoring to locations outside the arena is a distinguishing feature of egocentric neurons in 29e compared to egocentric neurons of other regions (Fig. 3F). We further investigated the anchoring properties of egocentric 29e neurons by analyzing their preferred bearing angles estimated as described in fig. S8. The distribution of bearing angles showed a significant bias that was contralateral to the recording hemisphere (right hemisphere), with most anchor points concentrated in the animal’s left-to-back-left sector, corresponding to the peripheral visual field (Fig. 3G). Supplementary analyses of other regions also showed a contralateral preference in MEC and PA, but not in PO and PR, perhaps due to low statistical power (fig. S9C).

### Area 29e differed from neighboring parahippocampal regions in theta and gamma modulation

Having demonstrated differences between 29e and the other regions in the strength of landmark anchoring and responses to changes in visual inputs, we next explored physiological characteristics that might underlie the network mechanisms governing the interactions among these regions. Area 29e differed from the other regions in both its local field potential (LFP; Fig. 4A) properties and its spike– LFP coupling. All regions showed significant LFP oscillatory power in the theta (5-10 Hz), beta (20– 35Hz), and gamma (50–120 Hz) bands (Fig. 4B). Given the prominent roles of theta and gamma rhythms in hippocampal-parahippocampal spatial processing, we focused subsequent analyses on these two frequency bands. Area 29e had the lowest theta power (mean ± s.e.m. = 0.687 ± 0.072), which was determined to be significantly different from MEC (1.148 ± 0.028) and PO (1.365 ± 0.058), but not PA (1.036 ± 0.043) or PR (0.735 ± 0.056) by a linear mixed-effects model accounting for non-independence among LFP signals from the same rat (fig. S10A). Gamma-band power in 29e was statistically comparable to other regions (fig. S10B).

**Fig. 4.**
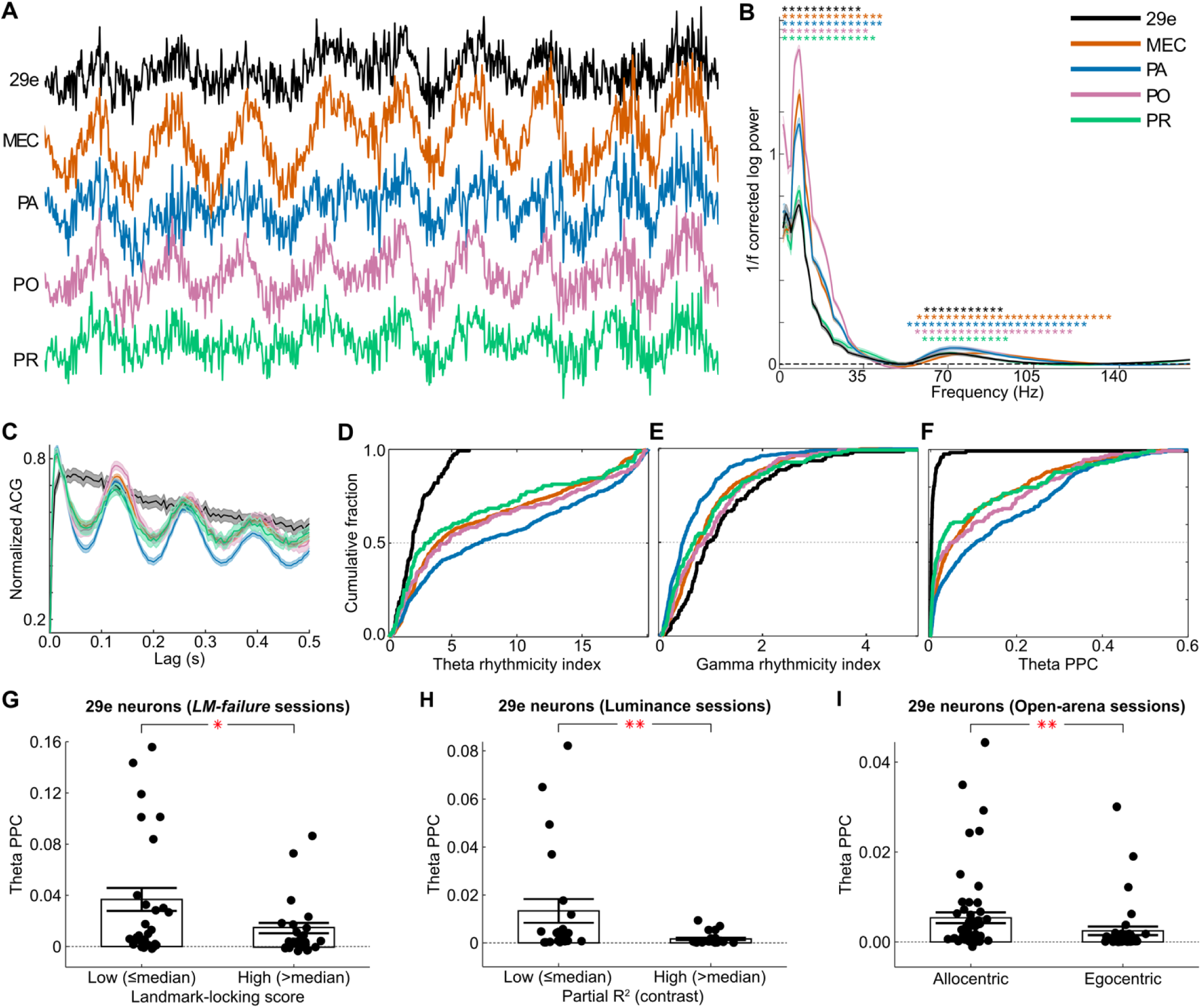
LFP-related differences between area 29e and neighboring parahippocampal areas (MEC, PA, PO, and PR). **(A)** Representative LFP traces, simultaneously recorded from 29e, MEC, PA, PO, and PR. **(B)** Mean ± 2 s.e.m. log-power spectral density (PSD) of LFPs in all recorded regions (unique recording locations for 29e: n = 20, MEC: n = 104, PA: n = 64, PO: n =22, PR: n = 24). PSDs are computed after correcting the aperiodic 1/f background in the LFP recordings. Asterisks mark the statistical significance of the power at a particular frequency, as measured by a cluster permutation test to control familywise error rate at α = 0.05 (*53*). **(C)** Population mean ± s.e.m. of the normalized spike autocorrelograms (ACGs) across single neurons in each region (colors as in panel B). Area 29e shows a monotonic, non-oscillatory decay, whereas other regions exhibit prominent oscillations at the theta frequency. **(D-F)** Cumulative distributions of single-neuron physiological measures across recorded regions (area 29e: n = 123, MEC: n = 566, PA: n = 414, PO: n = 133, PR: n = 124). *Panel D* shows significantly lower theta rhythmicity of spike ACGs in 29e (median [IQR] = 2.01 [1.29, 2.89]) than other regions (MEC = 3.91 [2.06, 12.4], PA = 7.08 [2.35, 15.53], PO = 4.67 [1.79, 13.19], and PR = 3.08 [1.44, 10.29]; Kruskal-Wallis test χ^2^(4) = 100.4, *p* < 0.001; post-hoc Wilcoxon rank-sum tests with Holm-Bonferroni correction 29e vs. MEC: *Z* = –8.1, *p <* 0.001; 29e vs. PA: *Z =* –9.4, *p <* 0.001; 29e vs. PO: *Z =* –6.2, *p <* 0.001; 29e vs. PR: *Z =* –4.1, *p <* 0.001). *Panel E* shows significantly larger gamma rhythmicity in 29e (median [IQR] = 0.95 [0.63, 1.69]) than other regions (MEC = 0.76 [0.44, 1.25], PA = 0.46 [0.22, 0.91], PO = 0.87 [0.34, 1.43], PR = 0.69 [0.30, 1.38]; Kruskal-Wallis test χ^2^(4) = 91.1, *p* < 0.001; post-hoc Wilcoxon rank-sum tests with Holm-Bonferroni correction 29e vs. MEC: *Z* = 3.40, *p <* 0.001; 29e vs. PA: *Z =* 7.7, *p <* 0.001; 29e vs. PO: *Z =* 2.2, *p =* 0.025; 29e vs. PR: *Z =* 3.2, *p =* 0.002). *Panel F* shows significantly reduced theta PPC of neurons in 29e (median [IQR] = 0.0016 [0.0004, 0.0052]) than in other regions (MEC = 0.0496 [0.0119, 0.1629], PA = 0.1040 [0.0192, 0.2773], PO = 0.0376 [0.0046, 0.2066], and PR = 0.0218 [0.0044, 0.1652]; Kruskal-Wallis test χ^2^(4) = 235.7, *p* < 0.001; post-hoc Wilcoxon rank-sum tests with Holm-Bonferroni correction 29e vs. MEC: *Z* = –13.7, *p <* 0.001; 29e vs. PA: *Z =* –14.1, *p <* 0.001; 29e vs. PO: *Z =* –8.7, *p <* 0.001; 29e vs. PR: *Z =* –8.4, *p <* 0.001). **(G-I)** Reduced spike–LFP theta coupling in 29e neurons with higher landmark locking (G), stronger landmark-contrast tuning (H), and egocentric (vs. allocentric) HD tuning (I). Dots indicate single neurons, bars show mean ± s.e.m., *Panel G:* Spatially correlated 29e neurons recorded in *LM-failure* sessions in the Dome were split at the median landmark-locking score. The low landmark-locking score group exhibited significantly higher theta PPC than the high landmark-locking score group (t-test on arcsinh-transformed PPC values; *t*(55) = 2.51, *p* = 0.015). *Panel H*: 29e neurons recorded in luminance-manipulation sessions were split at the median of their partial R^2^ values. The low partial R^2^ group showed significantly higher theta PPC than the high partial R^2^ group (t-test on arcsinh-transformed PPC values; *t*(43) = 2.94, *p* = 0.005). *Panel I*: 29e neurons classified as allocentric-HD tuned in open arena sessions had significantly higher theta PPC than their egocentric counterparts (Wilcoxon’s rank-sum test on asinh-transformed PPC values: *Z* = 2.63, *p* = 0.009).

These regional differences in LFP were matched at the level of single-neuron firing properties. Mean firing rates in the Dome were lowest in 29e compared to the other regions (fig. S10C), although the difference was statistically significant only relative to MEC, PA and PR. Theta rhythmicity (*22*, *49–51*) in the neurons’ spike autocorrelograms was significantly weaker in 29e than the other regions (Fig. 4C-D). In contrast, gamma-band rhythmicity was significantly higher in 29e (Fig. 4E). Spike coupling to LFP theta oscillations, quantified by the pairwise phase consistency (PPC) of the spike-phase distribution (*52*), was significantly lower in 29e than other recorded regions (Fig. 4F).

We hypothesized that within area 29e, neurons with stronger sensory-driven, landmark-related properties would be less modulated by the theta rhythm, whereas other 29e neurons would more closely resemble the strongly theta-modulated populations in MEC, PA, PO, and PR. To test these hypotheses, we first focused on *LM-failure* sessions, i.e., the sessions for which we could dissociate landmark-anchored neurons of 29e from non-anchored ones. Splitting the 29e population into two halves based on their landmark-locking scores showed that neurons in the top half had significantly lower theta PPC than those in the bottom half (Fig. 4G; fig. S11A-C). Next, focusing on luminance-manipulation sessions, we similarly observed that 29e neurons in the top half of the landmark-contrast tuning strength had significantly lower theta PPC than those in the bottom half (Fig. 4H; fig. S11D-F). Finally, examining open-arena sessions, we found that 29e neurons classified as egocentric exhibited significantly lower theta PPC than those classified as allocentric (Fig. 4I; fig. S11G). These results demonstrate that neurons with more sensory-driven properties prevalent in 29e (landmark locking, landmark-contrast tuning, and egocentric tuning) were less strongly coupled to LFP theta, the dominant rhythm of the rodent hippocampal-parahippocampal system.

### Breakdown of theta- and gamma-mediated interactions between area 29e and MEC LFPs preceded the loss of landmark control across the parahippocampal system

To investigate the interactions and coordination between 29e and MEC networks under conditions of *LM-control* and *LM-failure*, we measured coherence and Granger causality between 29e and MEC at theta and gamma frequencies. During periods of stationary landmarks (i.e., the initial epoch with experimental gain = 1), average LFP–LFP coherence across 29e–MEC tetrode pairs (n = 32; 4 rats) exceeded chance levels throughout the theta band and at multiple gamma bands (Fig. 5A), paralleling the spectral profile of significant oscillatory power in both regions (Fig. 4B). To assess the directionality of influence between 29e and MEC LFPs, we computed the frequency-resolved Granger causality (GC) spectra (*54–57*) and calculated a *directed-influence asymmetry index* (DAI) as (GC_29e →MEC_ – GC_MEC →29e_)/(GC_29e →MEC_ + GC_MEC →29e_) (*58*). A significantly negative DAI indicates a stronger drive from MEC onto 29e, whereas a significantly positive DAI indicates a stronger drive from 29e to MEC. This analysis revealed that MEC significantly drove 29e in the theta band, whereas 29e significantly drove MEC in the gamma band (Fig. 5B).

**Fig. 5.**
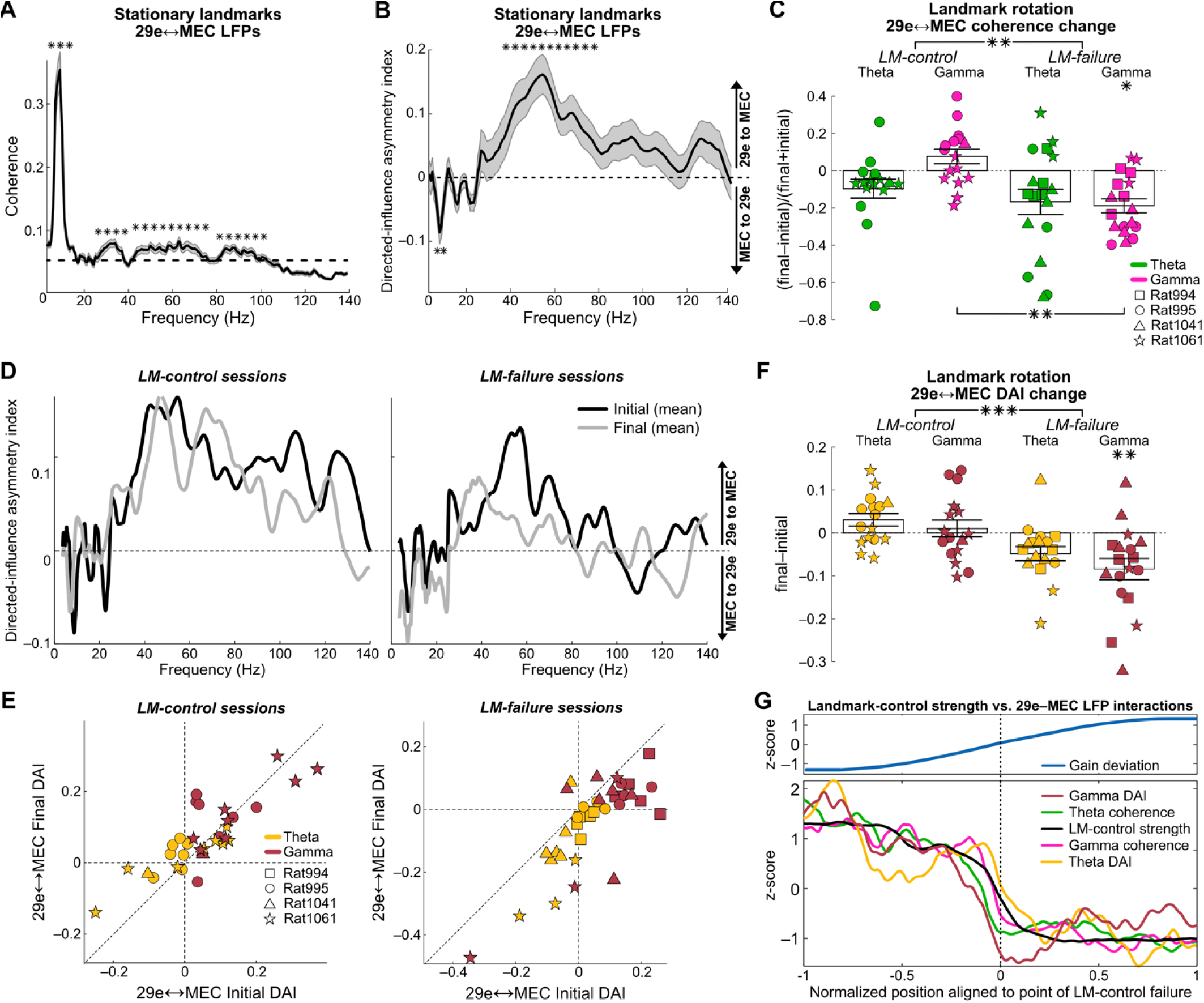
**Theta-gamma coordination between area 29e and MEC.** (**A**) LFP–LFP coherence spectrum between 29e and MEC LFPs when landmarks were stationary. Gray shaded area surrounding the solid black line indicates the mean ± s.e.m. of coherence spectra across 32 tetrode pairs. Asterisks mark the statistical significance of the coherence at a particular frequency, as measured by a hierarchical cluster-based permutation test to control familywise error rate at α = 0.05. (**B**) Directed-influence asymmetry index (DAI) between 29e and MEC LFPs when landmarks were stationary. The convention is the same as panel A. (**C**) Normalized changes in theta**-** and gamma-band coherence between 29e and MEC LFPs from the initial epoch to the final epoch across *LM-control* and *LM-failure* sessions. Each data point represents a 29e-MEC tetrode pair; bars represent mean ± s.e.m. The model included fixed effects of session type (*LM-control* vs *LM-failure*), frequency band (theta vs. gamma), and their interaction, and random effects accounting for repeated measurements within rats and tetrode pairs (see Methods for details). Coherence changes in *LM-control* sessions did not differ significantly from zero (theta band: *t*(8.3) = –1.6, *p* = 0.300; gamma band: *t*(8.3) = 1.4, *p* = 0.300), whereas coherence changes in *LM-failure* sessions were negative, reaching statistical significance in the gamma band (*t*(9.7) = –3.3, *p* = 0.035) but not in the theta band (*t*(9.7) = –2.8, *p* = 0.062). Holm-Bonferroni correction was applied to all four comparisons. The model also revealed a significant main effect of session type (LM-*control* vs. *LM-failure: F*(1, 31.1) = 10.6, *p* = 0.003) and a significant session type × frequency-band interaction effect (*F*(1,33.8) = 5.1, *p =* 0.030) on within-session coherence changes, but no significant main effect of the frequency band (*F*(1,3.3) = 1.4, *p =* 0.313). Post-hoc pairwise comparisons indicated that the interaction was driven by a difference in gamma-band coherence changes between *LM-control* and *LM-failure* sessions (*t*(44.4) = 3.9, *p =* 0.001; Holm–Bonferroni corrected across four post-hoc comparisons). (**D**) Mean DAI spectra across tetrode pairs in the initial (black) and final (grey) epochs of *LM-control* (left) and *LM-failure* (right) sessions. The pattern of frequency-dependent DAI between epochs is similar across initial and final epochs in *LM-control* sessions. By contrast, the DAI in *LM-failure* sessions shifts downward in both theta- and gamma-bands, indicating a shift toward stronger theta drive from MEC onto 29e (in comparison to theta drive in the reciprocal direction) and toward weaker gamma drive from 29e to MEC in the gamma band. (**E**) Theta- and gamma-band directed-influence asymmetry index (DAI) values in the initial epoch (x-axis) with stationary landmarks vs. the final epoch (y-axis) with rotating landmarks across *LM-control* (left) and *LM-failure* (right) sessions. Each data point represents a tetrode pair. The diagonal dashed line represents the identity y = x (i.e., initial DAI = final DAI). In *LM-control* sessions (left), points cluster around the identity line, indicating minimal change. In *LM-failure* sessions (right), however, most points fall below the identity line, indicating a systematic decrease in DAI. The within-session DAI decrease has opposite implications for the two frequency bands: in the theta-band, where initial DAI values were predominantly negative, the further decrease points to a shift toward enhanced MEC→29e drive during the loss of landmark control; by contrast, in the gamma-band, where initial DAI values were predominantly positive, the decrease indicates a shift toward diminished 29e→MEC drive during the loss of landmark control. (**F**) Initial to final epoch change in theta- and gamma-band DAI (quantified as DAI_final_ – DAI_initial_) between 29e-MEC tetrode pairs across *LM-control* and *LM-failure* sessions. We analyzed data using a linear mixed-effects model that has the same random- and fixed-effects structure as in panel C. Like panel C, the fitted model was first used to test whether DAI changes differed from zero in each of the four conditions (Control–Theta, Control–Gamma, Failure–Theta, Failure–Gamma; Holm– Bonferroni corrected across four tests): DAI changes in *LM-control* sessions were slightly positive but not significant (theta band: *t*(7.8) = 1.9, *p* = 0.199; gamma band: *t*(8.3) = 0.5, *p* = 0.660). By contrast, DAI changes in *LM-failure* sessions were negative, reaching statistical significance in the gamma band (*t*(8.7) = –4.4, *p* = 0.008) and a statistical trend in the theta band (*t*(8.7) = –2.9, *p* = 0.058). The model also revealed a significant main effect of session type (*LM-control* vs. *LM-failure*: *F*(1, 43.3) = 20.7, *p* < 0.001), consistent with an overall decrease in DAI from initial to final epoch in *LM-failure* sessions in contrast to an overall DAI increase in *LM-control* sessions. There was no significant main effect of the frequency band or its interaction with the session type. (**G**) Average lapwise trajectories of normalized gain deviation (i.e., the unsigned difference between the current value of the experimental gain and the initial baseline gain of 1) and of 29e–MEC LFP coupling measures (bottom) across 14 tetrode pairs from *LM-failure* sessions, aligned to the point of landmark-control failure (x = 0). The normalized x-axis interval [–1, 1] corresponds to approximately 75 laps. One rat was excluded from this visualization due to instability of firing fields of its non-29e neurons with respect to landmarks during stationary periods, but data from this rat were included in quantitative analyses. For the visualization, all traces were z-scored, smoothed with a moving average filter (half-width = 2 laps), aligned to the point of landmark-control failure (vertical dashed line at 0), time normalized (i.e., temporally scaled to also align the beginning and end points of all experiments), and averaged across tetrode pairs. For each session, the failure point was determined by visual inspection of lapwise firing rate plots of non-29e neurons relative to landmarks (as in Fig. 2D–E). Overall, the plots show that as gain deviation increased (top), landmark-control strength (bottom; blue) progressively declined, with a marked drop occurring at the point of landmark failure. This trend was closely tracked by the trajectories of 29e–MEC LFP coupling measures (theta-band coherence: median [IQR] Pearson’s rho = 0.64 [0.28, 0.78]; gamma-band coherence = 0.66 [0.39, 0.79]; theta-band DAI = 0.41 [0.32, 0.53]; gamma-band DAI = 0.41 [0.30, 0.56]), but with apparent temporal (lapwise, x-axis) offsets. Specifically, reduction in gamma DAI—primarily reflecting loss of 29e→MEC gamma drive (see panel D-E)—preceded both the decline in landmark-control strength and changes in other LFP coupling measures. The decline in landmark-locking strength was largely synchronous with the decline in gamma coherence (median [IQR] lag (lap) = –0.2 [–1.4, 1.7]; Wilcoxon sign-rank test *Z* = –0.22, *p* = 0.823) and theta-band DAI (–1.6 [–3.6, 1.5]; *Z* = –0.67, *p* = 0.498), but was significantly preceded by changes in theta-band coherence (–0.6 [–1.6, 0]; *Z* = –2.20, *p* = 0.027) and gamma-band DAI (–3.0 [–4.0, –2.3]; *Z* = −2.55, *p* = 0.010). Further testing with linear mixed-effects models to account for non-independence of observations within animals and within tetrode pairs (see Methods for random-effects structure) confirmed the significance of the these associations between changes in landmark-control strength and changes in both coherence (theta: *β* = 0.24, *t*(2.9) = 4.6, *p* = 0.027); gamma: *β* = 0.34, *t*(3.1) = 7.3, *p* = 0.009) and DAI (theta: *β* = 0.37, *t*(3.0) = 7.2, *p* = 0.011; gamma: *β* = 0.31, *t*(2.9) = 3.7, *p* = 0.035).

We quantified, for each 29e-MEC tetrode pair, the changes in LFP–LFP coherence from the initial epoch with stationary landmarks to the final epoch with rotating landmarks. In *LM-control* sessions, neither theta nor gamma coherence changed significantly. By contrast, in *LM-failure* sessions, coherence decreased in both frequency bands, with gamma reaching significance and theta only approaching it (Fig. 5C). Overall, coherence decreased significantly more in *LM-failure* sessions than in *LM-control* sessions. This difference between session types was primarily driven by changes in gamma coherence.

These results demonstrate that LM-failure was associated with gamma-band desynchronization between 29e and MEC LFPs, with weaker evidence for a similar trend in the theta band.

LM-failure was also associated with a shift in directionality of 29e-MEC LFP interactions. In *LM-control* sessions, the DAI curve remained stable from the initial to the final epoch across the whole frequency range (Fig. 5D-E, left). Thus, even though gain manipulation in the Dome creates a conflict between path integration and landmarks (*12*, *59*), the strength of gamma-mediated, 29e→MEC drive and the strength of theta-mediated, MEC→29e feedback did not change so long as the landmarks maintained control over the parahippocampal representations of space. However, in *LM-failure* sessions where landmark control was lost, the DAI curve shifted downwards (Fig. 5D-E, right), indicating a *decrease* in the gamma-mediated, 29e→MEC drive and an *increase* in the theta-mediated, MEC→29e drive (note that a more negative DAI reflects a greater MEC→29e drive). The DAI curve’s downward shift was significant for gamma but only approached significance for theta—a pattern similar to the coherence results (Fig. 5F; see fig. S12A for further breakdown of the components of the DAI). Overall, DAI changes in *LM-failure* sessions were significantly different from those in *LM-control* sessions. Thus, the loss of visual landmark control over the parahippocampal representations during a session was associated with a breakdown of the normal gamma-band influence from 29e to MEC.

We next asked whether the changes in 29e-MEC LFP interactions emerged before or after the loss of landmark control in *LM-failure* sessions. To address this question, we quantified the ongoing strength of landmark control by measuring how consistently the non-29e neuronal population activity vector repeated across laps in the reference frame of the rotating landmarks (see Methods). The trajectory of this relative landmark-control strength closely matched the ongoing changes in 29e-MEC LFP coupling measures (coherence and DAI) in both theta and gamma bands, albeit with some temporal differences (Fig. 5G). Linear mixed-effects models that account for non-independence of observations within animals and within tetrode pairs confirmed the statistical significance of these positive associations for both LFP coupling measures and both frequency bands. Cross-correlation analysis, comparing observed correlations to a null distribution of correlations from circularly shuffled landmark-control strength trajectories, revealed significant positive correlations (α= 0.05) in 11 of the 18 tetrode pairs for theta-band coherence, 13 of the 18 for gamma-band coherence, and 10 of the 18 for both theta- and gamma-band DAI. (all *p* < 0.001 as per binomial tests against a null probability of 0.05).

Analysis of cross-correlation lags using significant tetrode pairs showed that ongoing changes in landmark-control strength were synchronous with changes in gamma-band coherence and theta-band DAI across laps but were significantly preceded by changes in theta-band coherence and gamma-band DAI. A direct comparison between these leading indicators showed that changes in gamma-band DAI occurred significantly earlier than changes in theta-band coherence (*Z* = –2.34, *p* = 0.019). Taken together with the existence of landmark-anchored neurons within 29e and the dominant directionality of gamma drive from 29e to MEC during stable landmark control, this finding suggests reduced 29e-to-MEC gamma drive as an early indicator of impending landmark-control failure.

### Gamma-band feedforward coupling and local circuit connectivity support coordination of landmark signals in 29e with parahippocampal circuits

To determine whether the spiking activity of 29e and MEC neurons were consistent with the LFP interactions reported in Fig. 5, we examined the coupling of spatially correlated 29e and MEC neurons to the other region’s LFP in the Dome. Consistent with the LFP–LFP coupling result (Fig. 5A), the spectral profile of cross-regional spike–LFP pairwise phase consistency (PPC) showed clear peaks in the theta and gamma bands during periods of stationary landmarks in the Dome (Fig. 6A). In the gamma band, the coupling of 29e neurons’ firing to MEC was significantly stronger than that of MEC neurons to 29e, whereas in the theta band, the coupling of MEC neurons’ firing to 29e was significantly stronger than that of 29e neurons’ to MEC. This pattern matched the reversal in the dominant directionality of LFP–LFP Granger Causality influences across theta and gamma bands shown in Fig. 5B. While cross-regional PPC in both bands did not change significantly during gain manipulations in *LM-control* sessions, significant reductions in PPC were present during *LM-failure* sessions, specifically in the coupling of MEC neurons to 29e theta and 29e neurons to MEC gamma (Fig. 6B).

**Fig. 6.**
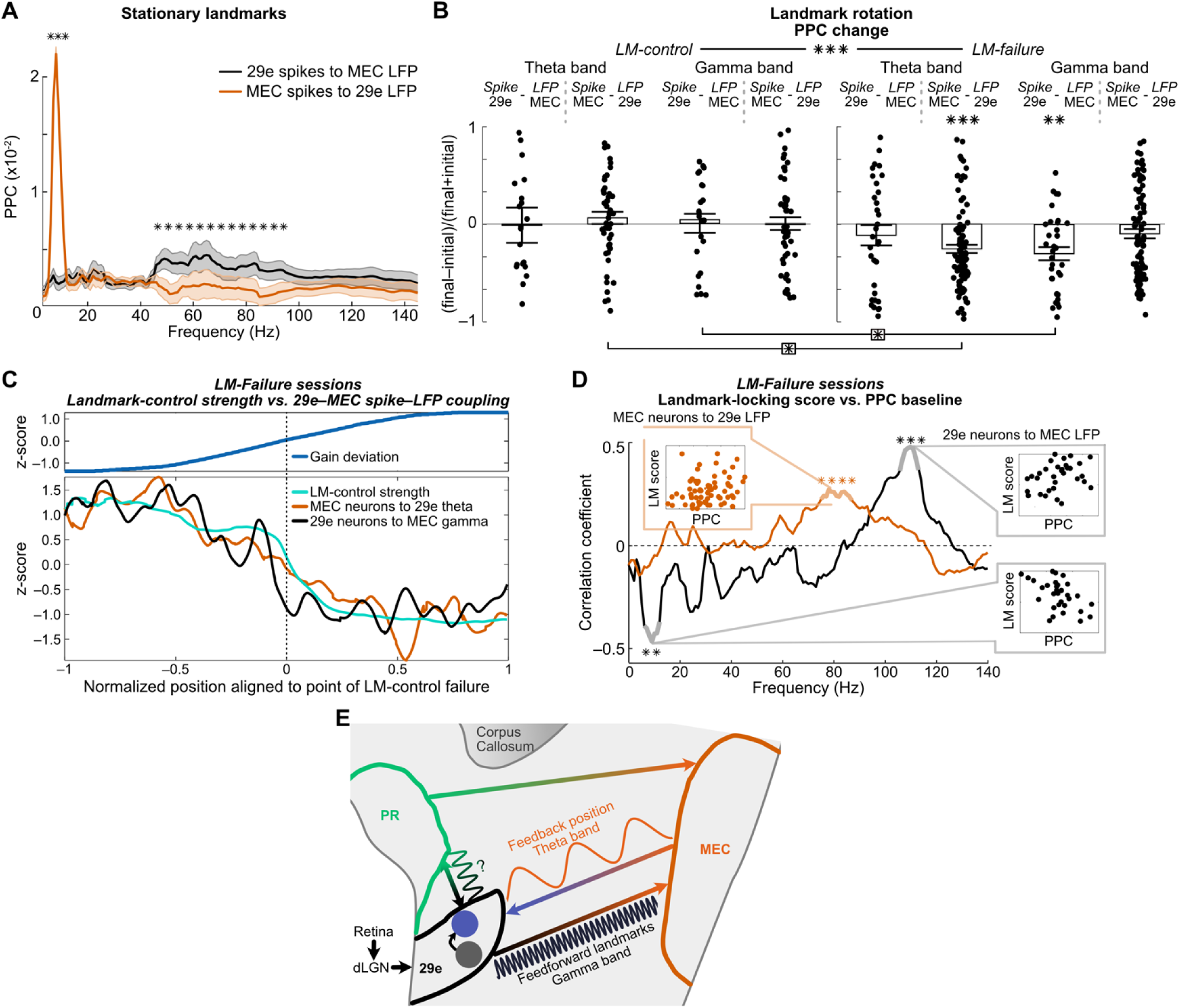
MEC-gamma coupling, local connectivity of landmark-anchored 29e neurons, and a conceptual model of inter-areal communication between area 29e and other parahippocampal areas. **(A)** Cross-regional spike–LFP coupling spectrum between 29e and MEC when landmarks were stationary. Shaded area surrounding the solid black (29e neurons’ coupling to MEC LFP) and orange (MEC neurons’ coupling to 29e LFP) lines indicate the mean ± s.e.m. of the respective spike–LFP spectra. Asterisks mark the statistically significant differences determined by a cluster-based permutation test to control familywise error rate at α =0.05. **(B)** Normalized changes in cross-regional spike–LFP coupling ([final – initial]/[final + initial]) for each neuron between 29e and MEC across *LM-control* and *LM-failure* sessions. Each dot represents a single neuron; bars indicate mean ± s.e.m. Asterisks above bars denote significance levels compared to zero; asterisks below brackets denote significance between respective conditions. In *LM-control* sessions, coupling changes did not differ significantly from zero in either theta band (29 neurons-to-MEC theta: median [IQR] = –0.02 [–0.42, 0.44]; Holm-Bonferroni corrected Wilcoxon sign-rank test *Z =* 0.2, *p* = 1.000; MEC neuron–29e theta: 0.07 [–0.21, –0.32]; *Z* = 1.1, *p* = 1.000) or gamma band (29e neurons-to-MEC gamma: 0.04 [–0.49, 0.36]; *Z* = –0.3, *p* = 1.000; MEC neurons-to-29e gamma: –0.06 [–0.32, 0.26]; *Z* = 0.09, *p* = 1.000). By contrast, *LM-failure* sessions showed significant reductions in couplings of 29e neurons-to-MEC gamma (–0.31 [–0.55, –0.02]; *Z* = –3.56, *p* = 0.001) and of MEC neurons-to-29e theta (–0.32 [–0.52, –0.08]; *Z* = –5.21, *p* < 0.001) but no significant change in couplings of 29e neurons-to-MEC theta (–0.15 [–0.77, 0.41]; *Z* = –1.12, *p* = 0.262) or of MEC neurons-to-29e gamma (–0.16 [–0.51, 0.27]; *Z* = -–2.00, *p = 0.092*). Three-way repeated measures ANOVA (factors: session type, frequency-band, and neuron region-LFP region pair identity) revealed significant main effect of the session type (*F(*1,202) = 9.7, *p* = 0.002) and a significant three-way interaction (*F*(1,202) = 4.0, *p* = 0.048) but no significant main effects of the frequency-band (*F*(1,202) = 0.3, *p =* 0.583) or neuron-LFP region pair (*F*(1,202) = 0.54, *p =* 0.463), or any two way interaction. Holm-Bonferroni corrected post-hoc pairwise Wilcoxon tests within each frequency band identified two significant differences pertaining to the interaction effect. Within the theta-band, the changes in MEC neuron-29e LFP theta coupling was significantly lower in *LM-failure* sessions than *LM-control* sessions (*Z = –*4.4, *p* < 0.001). Within the gamma-band, the changes in 29e neuron-MEC LFP coupling across *LM-failure* sessions was significantly lower than the changes in MEC neuron-29e LFP coupling across *LM-control* sessions (*Z* = –3.2, *p* = 0.012). **(C)** Average lapwise trajectories of gain deviation (top; defined as in Fig. 5G) and of cross-regional spike–LFP coupling measures across *LM-failure* sessions (bottom). Same format as Fig. 5G. Only neurons that were active in at least 75% of laps and exhibited split-session stability were included (n = 32 from area 29e, n = 95 from MEC) to ensure reliable lap-by-lap estimates of coupling. The same pre-processing steps were applied as in Fig. 5G, except that the smoothing filter used a half-width of 2.5 laps to compensate for the sparser activity of individual neurons, which have spatially localized firing fields, compared to continuously varying LFP signals. Overall, the plots show that both spike–LFP coupling measures declined as experimental gain deviated from 1 (top), closely tracking the progressive decline in landmark-control strength (blue; 29e neurons-to-MEC gamma: median [IQR] Pearson’s rho = 0.60 [0.52, 0.70]; MEC neurons-to-29e theta: 0.49 [0.26, 0.69]). Notably, the coupling of 29e neurons to MEC gamma (black) showed a rapid decline before the drop in landmark-control strength (blue; median [IQR] cross-correlation lag (lap) = –3.0 [–4.2, 0]; Wilcoxon signed-rank test *Z* = –2.61, *p* = 0.008). By contrast, MEC neurons’ coupling to 29e theta (pink) also declined but did so more gradually without showing any clear temporal lead or lag relative to landmark-control strength (0.1 [–3.2, 3.7]; *Z =* 0.06, *p* = 0.956). Applying linear mixed-effects modeling that accounted for non-independence of observations within animals and within neurons (see Methods for details), we quantitatively confirmed significant positive associations between ongoing changes in landmark-control strength and changes in both spike–LFP coupling measures (Holm-Bonferroni correction; 29e neurons to MEC gamma: *β* = 0.37, *t*(2.9) = 4.4, *p* = 0.042; MEC neurons to 29e theta: *β* = 0.22, *t*(2.7) *=* 3.55, *p* = 0.044). **(D)** Correlation between baseline cross-regional spike–LFP PPC and landmark-locking score measured during the final 12 laps with rotating landmarks during failure sessions. Spearman’s correlation coefficient is shown as a function of frequency (x-axis) for MEC neurons to 29e LFP (orange) and for 29e neurons to MEC LFP (black). Insets show example scatterplots of landmark-locking scores (LM score) versus PPC values (arcsinh-transformed for visualization) at selected frequencies. Asterisks and lighter colored segments indicate frequency ranges with significant nonzero correlations (cluster-based permutation test to control familywise error rate at α = 0.05). **(E)** Conceptual model of area-29e–mediated landmark control of path integration. Based on established anatomy and our physiological findings, we propose that gamma-band drive from 29e conveys feedforward landmark information to MEC, whereas theta-band drive from MEC conveys feedback, path-integration–based position estimates to 29e. Area 29e receives low-latency landmark signals via a direct retina→dLGN→29e pathway and is reciprocally connected to downstream areas MEC and PR. We propose that visual landmark signals differentially innervate 29e neurons, yielding the gradient of landmark anchoring observed in our data. For simplicity, the schematic renders this continuum as two subpopulations, landmark-locked (dark gray) and landmark-unlocked (purple). Feedforward output from 29e could arise from the strongly landmark-anchored subpopulation, consistent with their stronger spike–LFP coupling to MEC gamma during landmark control and the large reduction of gamma coupling when landmark control failed. We further hypothesize that MEC feedback preferentially innervates the less landmark-anchored subset, accounting for their stronger theta coupling and functional similarities to neurons in other parahippocampal areas, including MEC. MEC, the putative path-integration hub, is often modeled as a continuous attractor network; during landmark failure, the MEC neuron population remained largely coherent— either landmark-controlled or not—consistent with attractor dynamics. Within 29e, putative monosynaptic connections are biased from landmark-locked to landmark-unlocked neurons (fig. S13), consistent with the skewness in the distribution of (presynaptic – postsynaptic) landmark-locking scores. Consequently, the 29e landmark-unlocked population integrates inputs from locked 29e cells as well as from other parahippocampal areas. An indirect route from 29e to MEC may be relayed via PR, although effective communication between 29e and PR remains to be shown. Together, these interactions form a functional loop in which 29e provides sensory-based, landmark-anchored positional signals to MEC path-integration circuits, while MEC, combining the landmarks with self-motion cues, calculates an internal position signal that feeds back to 29e. Arrows denote established anatomical connectivity; colored waveforms indicate relevant frequency bands.

To address whether the disruption of spike–LFP coupling (particularly MEC neurons-to-29e theta and 29e neurons-to-MEC gamma) emerged before or after the loss of landmark control, we analyzed cross-regional PPC on a lap-by-lap basis in *LM-failure* sessions, analogous to our LFP–LFP analysis (Fig. 5G). Ongoing changes in landmark-control strength closely tracked lap-by-lap changes for both coupling measures, albeit with some temporal differences (Fig. 6C). We confirmed the consistency of these positive relationships across animals by linear mixed-effects modeling that account for non-dependencies among observations. Cross-correlation analysis, which accounts for temporal offsets between landmark-control strength and spike-LFP coupling, revealed significant correlations (α = 0.05, compared to circularly shuffled distributions) in 17 of 32 neurons from area 29e and 38 out of 95 from MEC (both *p* < 0.001 as per a binomial test against a null probability of 0.05). Analysis of lags at the peak correlation showed that changes in 29e→MEC gamma coupling significantly preceded changes in landmark-control strength, whereas changes in MEC→29e theta coupling were approximately synchronous with landmark-control strength. Direct comparison confirmed that changes in 29e neurons’ coupling to MEC gamma occurred significantly earlier than changes in MEC neurons’ coupling to 29e theta. This pattern mirrors the LFP–LFP results, where gamma-band DAI—reflecting the strength of 29e-to-MEC drive—was the earliest predictor of landmark-control failure (Fig. 5G). Thus, across both LFP–LFP and spike–LFP measures, the breakdown of gamma-mediated feedforward signaling from 29e to MEC precedes the failure of landmark control.

If 29e neurons provided landmark information to MEC via gamma oscillations, we would expect landmark-anchored 29e neurons to be more strongly coupled to the MEC LFP gamma than the non-anchored neurons. To test this hypothesis, we calculated, separately for 29e and MEC populations recorded in *LM-failure* sessions, the correlation between neurons’ landmark-locking scores (measured in the final epoch) and their baseline cross-regional PPC values (measured in the initial epoch with stationary landmarks) at different frequencies (Fig. 6D). 29e neurons displayed a striking frequency-dependent pattern: neurons with high landmark-locking scores showed strong coupling (i.e., high positive PPC-LM score correlation) to MEC gamma (particularly 100–120 Hz) but weak coupling (i.e., high negative PPC-LM score correlation) to MEC theta. This frequency-dependent pattern in 29e was significantly stronger than expected by chance (*p* = 0.006; fig. S12B). A similar analysis of MEC neurons’ coupling to 29e LFP showed that MEC neurons with high landmark-locking scores showed moderately strong coupling to 29e gamma (80-95 Hz), while those with weaker anchoring tended to couple more strongly to 29e theta. However, this frequency-dependent difference did not reach statistical significance (*p* = 0.078; fig. S12C). Furthermore, 29e neurons with high landmark-locking scores were more likely than those with low landmark-locking scores to lose their gamma-band coupling to MEC LFP (fig. S12D-E) as MEC spatial maps broke free from landmarks. These results suggest that gamma-band synchrony between 29e’s landmark-anchored subpopulation and MEC may support transfer of landmark information to parahippocampal path-integration circuits. Collectively, these findings suggest that landmark-anchored 29e neurons serve as a key source of landmark information for anchoring representations of parahippocampal path-integration circuits to the external world.

## Discussion

Brains build internal models to predict the world, and because the world is dynamic and predictions can be inaccurate, these models must be continually updated by sensory feedback (*60*, *61*). In navigation, the cognitive map embodies an internal model of space, and external landmarks provide allothetic spatial cues to recognize environments and locations within them and to correct accumulated errors in the path-integration-based estimates of position on this map. The neural origins of the landmark-based correction signal remain unknown in mammals. To address this gap, we performed simultaneous recordings in rats across five parahippocampal regions in an immersive VR apparatus that preserves naturalistic locomotion while affording accurate control over the position and salience of visual landmarks (*33*, *62*), thereby allowing us to create conflicts between internal path integration computations and feedback from external visual cues. When these conflicts disrupted landmark anchoring in most spatial neurons across MEC, PA, PO, and PR, many neurons maintained landmark-anchored firing in area 29e. Area 29e showed weaker theta in the LFP and weaker theta modulation of its neurons’ spiking compared to MEC, PA, PO, and PR. Area 29e also showed stronger egocentric and landmark-contrast tuning than other regions, and these sensory properties were markedly enhanced in its landmark-anchored subpopulation. These results reveal physiological and functional dissociations between area 29e and its neighboring regions, providing the first such characterization of area 29e in rodents. Taken together with the dynamic functional coupling between 29e and MEC that correlates with the degree of landmark control, our results identify a visually driven, egocentric-HD population in 29e that relays landmark signals to parahippocampal circuits supporting path integration.

Historically, anatomists treated area 29e variably: some incorporated it into retrosplenial cortex (*63*), PR (*64–66*) or PA (*67–69*), while others suggested that it may be a separate region (*70*, *71*). In this work, we adopted Blackstad’s definition of area 29e in rats as a cytoarchitecturally distinct triangular region interposed between PR and PA (*70*, *72*). Recent work has proposed that area 29e is the rodent homolog of the primate visual area prostriata (*26*, *29*, *73*), which is located next to the region of striate cortex (V1) that represents the most extreme periphery of the visual field and is characterized by broad orientation tuning, large receptive fields, and selectivity for fast-moving stimuli in the visual periphery (*28*, *27*, *74*, *75*).

Area 29e neurons in the present study displayed hallmarks of distal visual landmark representation— strong landmark anchoring, contrast sensitivity, and egocentric HD tuning anchored to locations beyond the open arena where distal cues were situated, with bearing angles preferentially falling in the peripheral visual field. Distal cues are thought to be a major signal that orients the cognitive map relative to the external world (*76–78*). Our data from rodent area 29e suggest that the orienting influence of distal visual cues may be mediated through peripheral vision—consistent with human behavioral studies reporting that individuals with restricted peripheral vision make greater errors in orientation and distance when navigating to remembered landmark locations (*79–81*). This peripheral specialization is anatomically supported by widespread input to 29e from medial V1 (which processes peripheral vision) and dorsal lateral geniculate nucleus (dLGN; *36*, *37*, *39*). The direct dLGN connection places 29e only two synapses from the retinal ganglion cells, potentially providing the fastest visual pathway to limbic circuits and explaining the short-latency responses observed in primates (*28*, *74*). Recent work has shown that dLGN drives gamma events promoting visual encoding in V1 (*82*) and that dLGN lesions severely disrupt the anchoring of HD tuning in PR to visual landmarks (*83*), together raising the possibility that dLGN efferents are the source of gamma-modulated activity characterizing visual landmark coding within 29e.

Anatomical and functional evidence suggests that 29e transmits these landmark signals to MEC, the putative hub of path integration (*9*, *84*, *8*, *15–18*, *20*, *85*). Efferents of 29e in rodents target PR and MEC (albeit more sparsely than PR), in addition to contralateral prostriata and visuomotor areas (*31*, *32*).

MEC is also innervated by PR (*86*, *87*), considered a candidate site for integration of an internal HD signal with external landmarks (*40*, *88*, *89*, *42*, *44*, *45*, *90*). This anatomical organization suggests two pathways via which 29e might deliver gamma-modulated visual landmark signals to MEC: a direct 29e→MEC pathway and an indirect 29e→PR→MEC pathway. Our LFP–LFP and spike–LFP coupling analyses provide functional evidence consistent with such pathways: 29e and MEC showed theta- and gamma-band synchronization, with a net gamma-band influence of 29e’s landmark-anchored population on MEC and theta-band influence of MEC on 29e. Notably, feedforward gamma-band influence declined before the failure of landmark control over MEC (as well as other parahippocampal populations) during large cue conflicts, supporting the idea that 29e transmits the landmark signals that anchor parahippocampal representations of space to the external world.

The *predictive coding framework* (*91*, *92*) is a general cortical organizing principle in human and nonhuman animals, proposing that rapidly changing sensory information and prediction errors are conveyed via feedforward connections, while slower contextual predictions are transmitted via feedback connections (*93–95*). Consistent with these timescales, feedforward and feedback anatomical pathways across sensory and frontoparietal networks operate at distinct frequency bands: bottom-up, feedforward signaling in high-frequency (gamma) and top-down, feedback signaling in low-frequency (alpha/beta) brain rhythms (*96*, *58*, *97–100*). We found an analogous organization within parahippocampal networks: area 29e influenced MEC in the gamma band, whereas MEC influenced 29e in the theta band, forming a gamma–theta-mediated closed loop that signals feedforward landmark information and feedback, path-integration-based positional predictions between 29e and MEC (Fig. 6E). When path integration and landmark-based positional cues became increasingly in conflict in our experiments, in many cases these frequency-specific interactions were disrupted and 29e-MEC rhythms became desynchronized, followed by a disruption of landmark control across the parahippocampal system. At the level of individual neurons, 29e neurons that were strongly coupled to MEC theta exhibited comparatively lower landmark anchoring and contrast sensitivity—similar to MEC neurons and thus presumably reflecting greater feedback influence from MEC—whereas those strongly coupled to MEC gamma exhibited stronger landmark anchoring, consistent with a more feedforward role in 29e signaling landmark information to MEC.

These results demonstrate how a visual area (29e) interacts dynamically with a set of high-order limbic regions (parahippocampal areas) to bind an internally generated, cognitive representation of a rat’s location to the visual landmarks present in the external world. The cognitive representations in the parahippocampal regions (MEC, PR, PA, and PO) maintain internal coherence (i.e., the maps remain stable within their own internal frame of reference), even when the maps become decoupled from the external sensory world (*85*, *101–105*). The binding of the map to the external world is apparently accomplished (at least in part) via the gamma-mediated transmission of visual input from 29e to the parahippocampal regions, with a theta-mediated feedback signal from these regions to 29e that maintains coherence between the cognitive and sensory processing regions during normal operation.

Under conditions of continual error between visual landmark-derived position information and path-integration-based updating of the rat’s position on its cognitive map, this theta- and gamma-mediated interareal coupling begins to break down, eventually resulting in the loss of landmark control over the cognitive map. Consistent with the predictive coding framework, these results suggest a general organizing framework for interareal coordination of diverse brain regions, necessary to produce adaptive, coordinated cognition and behavior in a distributed processing system composed of multiple, anatomically and functionally segregated brain areas.

## Acknowledgments

We thank Rahul Jakati, Ravikrishnan Jayakumar, Shahin Lashkari, Francesco Savelli, Manu Madhav, Katelan Balkissoon, Bethel Gebreselassie and Makenna Dixon for useful discussions and assistance with the experiments. We thank Daeyeol Lee and Sung Soo Kim for providing valuable comments on the manuscript.

## Author contributions

Conceptualization: GS, BK, NJC, JJK Experiments: GS, BK

Data analysis: GS, BK Writing: GS, BK, NJC, JJK

Funding acquisition: GS, BK, NJC, JJK Supervision: NJC, JJK

## Materials and Methods

### Subjects

Five adult Long–Evans rats (Envigo Harlan), 3 M (994, 995, and 1041) and 2 F (1061 and 1062), served as subjects. At the time of surgery, males weighed 440–470 g and were 6–9 months old, whereas females weighed 315–325 g and were 15–18 months old. Animals were housed individually in a temperature- and humidity-controlled room on a 12 h:12 h light/dark cycle; all behavioral training and electrophysiological recordings were carried out during the dark phase. Rats were handled nearly daily throughout habituation, training, surgery, and data-collection periods. Data from all five animals were pooled for all subsequent analyses, but individual rat data are shown in supplementary tables. Potential sex differences were confounded by non-uniform sampling of cell layers across animals, and small sample sizes for each sex prevented meaningful assessment of se as a biological variable. All procedures conformed to the National Institutes of Health guidelines and were approved by the Johns Hopkins University Institutional Animal Care and Use Committee.

### The Dome apparatus

The virtual-reality Dome (*12*, *33*) is a planetarium-style setup built around a hemispherical fiberglass shell with a 2.3-m inner diameter. A 15-cm aperture at the apex allowed light emitted from a ceiling-mounted projector (Sony VPL-FH30) fitted with a long-throw lens (Navitar ZM 70–125 mm). A 25-cm hemispherical first-surface mirror (JR Cumberland), mounted on a central pillar at the geometric center, reflected the projected image onto the inner surface of the Dome to create full-field visual stimulation. Visual scenes were rendered in C++ using OpenGL and pre-warped to compensate for the projection optics before display. To provide uniform, rotationally symmetric background illumination that did not overlap the landmarks, a narrow overhead ring was projected continuously, including during landmark-absent epochs. Behavior was monitored with an overhead near-infrared camera providing real-time, top-down video (see “*Experiment control and behavior tracking*”).

Rats locomoted along the outer edge of an annular table (outer diameter 152.4 cm, inner diameter 45.7 cm) positioned at the center of the Dome. An enclosure subtending 45° around the rat was attached to a motorized central pillar via radial boom arms to ensure that the animal remained near the table’s outer rim and to carry the transmitter unit of the neural recording system (in wireless recording sessions) or the headstage and Litz tethers (for wired recording sessions). The motorized central pillar incrementally advanced the enclosure to keep the rat near a predefined setpoint within the enclosure; it engaged only when the rat was within a defined window around that setpoint, ensuring the enclosure moved along with the rat. Liquid rewards (Ensure diluted 50% with water) were delivered pseudo randomly onto the table via a feed tube mounted on a boom arm diametrically opposite the enclosure; delivery timing and volume were controlled by a micro-peristaltic pump (Takasago fluidics) on the central pillar using custom software (*33*). This boom arm also held a plastic spreader and paper towels to disperse urine and food odors and to remove feces, minimizing the salience and stability of local olfactory cues. After each session the track was thoroughly wiped and then rotated by a multiple of 45° (one of eight possible orientations), altering the spatial relationship of any tactile cues to the rest of the apparatus. This prevented local tactile features on the annular track from becoming stable global landmarks.

### Closed-loop gain manipulation experiments in the Dome

During closed-loop gain manipulation sessions, the locations of visual landmarks (3 or 4 landmarks, depending on the session; Supplementary table 1) were updated in closed loop with the rat’s movement according to an experimental gain, G, defined as the ratio between landmark-frame and laboratory-frame speed of the animal. During these manipulations, landmarks were moved in unison, maintaining their relative configuration, and rotating as a coherent ensemble. Landmarks were stationary when G = 1. They were rotated along the circular track in the same direction as the animal when 0 < G < 1 or in the opposite direction when G > 1.

Every gain-manipulation session comprised three epochs.

- **Epoch 1**: G = 1 (veridical condition). During this Epoch, the animal ran 8–15 laps with stationary landmarks.
- **Epoch 2**: G was ramped linearly at a rate of 1/52 per lap until it reached its final value, G_final_.
- **Epoch 3**: G = G_final_ remained constant.

After the final epoch (Epoch 3), all landmarks were extinguished, leaving only a dim, non-polarizing ring for illumination.

The final gain value, G_final_, took the form of 1 ± n/13 (n = 0, 1, 2, 3, 4…), so that positions in the laboratory and landmark frames coincided only once every 13 laps when n > 0. Each rat first completed one to three baseline sessions with G_final_ = 1; thereafter the gain was systematically altered in following sessions. Sessions with more extreme values (G_final_ ≫ 1 or G_final_ ≪ 1) often showed landmark-control failure. Thus, to maximize the number of sessions that retained strong landmark control, more extreme values of G_final_ were chosen progressively across days and were seldom repeated on consecutive sessions. Experimenters were aware of the gain during data collection and analysis, and sample sizes were determined empirically without formal power calculations.

### Luminance manipulation experiments in the Dome

Landmarks and the background were implemented as separate OpenGL objects, allowing independent control of their luminance by scaling object intensities. Intensities were normalized to the projector’s dynamic range (0–1). Landmark (LM) luminance took values 1.00 (High) and 0.55 (Low); background (BG) luminance took values 0.47 (High) and 0.01 (Low). By providing greater LM luminance than BG luminance, these values ensured no “negative contrast” conditions (i.e., landmarks being darker than the background).

During luminance manipulation blocks consisted of equal-length epochs where landmarks were stationary (experiment gain G = 1). Within a given epoch, LM and BG luminance were held constant. The four illumination conditions (2 LM luminance levels × 2 BG luminance levels) were presented across 6-lap epochs in pseudorandom order predetermined by the experimenter, with the constraint that no condition was repeated in consecutive epochs and all conditions were experienced an equal number of times. Across a session, the luminance conditions changed only at epoch boundaries, and each of the four conditions were presented at least three times.

At the end of the luminance epochs, the experiment gain was abruptly changed to an extreme value (i.e., much larger or smaller than 1), which induced landmark failure, allowing us to test the landmark anchoring strength for all cells, as described in the “Landmark-locking score” subsection of Methods.

### Open-loop gain manipulation experiments in the Dome

To dissociate landmark motion from the animal’s self-motion in open loop experiments, visual landmark movement was controlled by a pre-recorded animal motion trajectory obtained from a different session. The reference trajectory (recorded as a ROS .bag file) contained the animal’s angular position along the circular track as a function of time. During open-loop sessions, this trajectory was replayed in real time and used to control landmark motion via the same experiment gain (G) parameter used in closed-loop experiments. If the open-loop session duration outlasted the reference trajectory duration, the recorded animal trajectory was looped from its beginning without interruption to provide continuous control of landmark position.

Each open-loop session followed a standardized gain protocol.

- **Epoch 1:** G = 1, during which landmarks were stationary for at least 8 laps (same as closed-loop).
- **Epoch 2:** G was ramped from 1 to a target value, as in closed-loop sessions. However, unlike closed-loop sessions where G was applied to the animal’s own movement, here G was applied to a pre-recorded trajectory from a reference session, thereby dissociating landmark motion from the animal’s own movement in the current session.
- **Epoch 3:** G underwent a series of step changes (“jumps”) to non-unity gain values across 2–4 sub-epochs, each sub-epoch lasting at least 8 laps. The set of gains across sub-epochs always included at least one value G > 1 and at least one value G < 1, ensuring the rat experienced landmarks moving in both directions relative to its own movement. As in Epoch 2, G was applied to the pre-recorded trajectory to move landmarks independently of the animal’s actual motion in the current session. Landmarks were extinguished at the end of the session.

### Open arena apparatus

During open-field foraging sessions, each rat foraged for randomly distributed precision chocolate pellets (Bio-Serv Dustless Precision Pellets, Purified), on a square platform that measured 137 × 137 cm and was surrounded by opaque walls 36.5 cm high. Experiments were conducted in two different recording rooms. In the larger room (approximately 330 × 700 cm) neural activity was transmitted wirelessly via a FreeLynx device, whereas in the second room (292 × 292 cm) signals were carried through a tether connected to a DigitalLynx SX system and Saturn commutator (Neuralynx, Inc.). Both rooms were dimly illuminated by overhead white LEDs and contained fixed distal cues—such as shelving, computers, and other laboratory equipment—that were kept in the same positions across days to provide stable landmarks for the animals. After every session the arena floor was wiped with 70 % ethanol. Any urine or feces were removed immediately during the session once the rat had moved away from that spot. Throughout each 30-40 min session the experimenter walked outside the arena while dispensing the chocolate precision pellets.

### Training

Rats were handled daily for 3-5 days initially to habituate them to human contact, then placed on a controlled feeding schedule to reduce body weight to approximately 80% of their *ad libitum* weight.

*Dome training:* Rats were first trained on a circular table in a room separate from the Dome apparatus. The table matched the Dome table in diameter but lacked the enclosure and automation (see “*<u>Dome</u> <u>apparatus</u>*”). Reward droplets (1:1 mixture of water and chocolate Ensure) were placed manually at arbitrary locations on the track, and the interval between rewards gradually increased to sustain running and delay satiation. Training continued until the rats consistently completed at least 80 laps in the same direction without experimenter intervention, typically after 3–4 weeks. The rats were then introduced to the Dome, which contained stationary visual landmarks, and were trained there until their lap counts matched those achieved on the open table, usually within one additional week. After hyperdrive implantation, the rats were retrained in the Dome and transitioned to automated reward delivery, in which drop timing varied to encourage continuous forward running with minimal pauses. Experiments began once the tetrodes reached the target recording regions and the rats consistently ran many laps.

*Open-arena training*: We acclimated rats to the open arena by giving them 2–3 consecutive days of free foraging for Bio-Serv Dustless Precision Pellets, intended to promote spontaneous exploration. After 1– 2 weeks they explored the entire arena readily and uniformly. After hyperdrive implantation, training continued until the rat, with the transmitter in place, traversed the arena broadly and almost continuously, pausing only briefly—a criterion typically achieved within 7–14 days after surgery.

Experiments began once this behavioral criterion, the Dome-training criterion, and accurate tetrode placement had all been met.

### Surgery and electrode implantation

After behavioral training in the Dome and open arena achieved desired behavioral performance, rats were implanted with custom hyperdrives carrying 18 independently drivable tetrodes. Each tetrode was composed of four twisted, heat-bonded nichrome wires (0.0017″ diameter, VG/HML insulation; California Fine Wire Co., CA, USA). Hyperdrives were built in-house using a custom design around a 72-channel interface board (EIB-72-QC; Neuralynx, Bozeman, MT, USA). During the 5–6 days preceding surgery, tetrode tips were gold-plated daily with a nanoZ system (White Matter LLC, Seattle, WA, USA) to reduce and stabilize impedance from 2–3 MΩ to ∼130 kΩ. Each assembled hyperdrive was sterilized in ethylene-oxide gas (12-h cycle) and allowed to aerate for 36 h before implantation.

Anesthesia prior to surgery was induced with 4% isoflurane in oxygen (1 L/min) followed by ketamine (80 mg/kg) and xylazine (10 mg/kg); anesthesia was maintained with isoflurane as needed. Local analgesia was provided by applying 0.15 mL of bupivacaine (Marcaine) or ropivacaine directly to the skull. Postoperatively, meloxicam (1–2 mg/kg; Ostilox, Meloxidyl, or Metacam) was administered orally for two days. Postoperative antibiotics (30 mg tetracycline and 0.15 mL of a 22.7% enrofloxacin solution) were given orally once daily for 10 days post-surgery. Under isoflurane anesthesia, a craniotomy was made over the right hemisphere, the rostral edge of the transverse sinus was exposed, and the dura was removed. The hyperdrive was positioned so that the posterior-most central tetrode entered the brain 350–530 µm anterior to the rostral edge of the transverse sinus and 4.30–4.55 mm lateral to bregma, yielding a mediolateral recording span of 3.4–5.45 mm lateral to bregma across animals. To increase effective tetrode travel within MEC, the hyperdrive was tilted 10° anteriorly in the sagittal plane. The craniotomy was then sealed with Kwik-Sil (World Precision Instruments).

Hyperdrives were anchored to the skull with jeweler’s screws; a screw over the cerebellum served as the recording ground. The implant was secured with a three-layer build of OptiBond (Kerr Dental), Charisma (Kulzer), and dental cement.

### Neural recordings

Wideband signals were digitized at 30 kHz and acquired on a computer running Neuralynx Cheetah software using either a wireless FreeLynx (Neuralynx) transmitter or, for wired sessions, a Litz tether connected to a Digital Lynx SX system by a unity gain 72 channel headstage (HS72QC, Neuralynx) attached to the implanted hyperdrive. Both Dome and open arena experiments were run with either wired or wireless configurations of this recording system. TTL pulses from the behavioral control computer (see Experimental control) were recorded as timestamped events on the acquisition system to enable post hoc synchronization of behavioral and neural data. Raw wideband data were saved as Neuralynx .nrd files and used for spike sorting.

### Experiment control and behavior tracking

Dome sessions were run using three computers: (1) an experiment-control PC, (2) a neural–acquisition PC running Cheetah (Neuralynx, Inc.), and (3) a video–tracking PC. Task logic and device control were implemented as Robot Operating System (*106*) nodes communicating continuously with a master node on the experiment-control PC. Synchronization was achieved by sending pseudorandom TTL pulses from the control PC to the acquisition system; these were time-stamped in Cheetah and used to align data streams across machines. Hardware/software integration and control details are described in a previous publication (*33*). Head position and 3D head orientation were estimated in real time at 30-45 fps using a single–camera, ROS–based tracker that used custom software and fiducial markers mounted on the hyperdrive in a known 3D configuration (*107*).

Open–arena sessions typically used two computers: one for neural acquisition with FreeLynx/Cheetah and one for behavior tracking running the single–camera 3D tracker. Synchronization between the neural acquisition computer and the behavior tracking computer was implemented using the Domain Time II software (Greyware Automation Products, Inc.). In a subset of sessions recorded with the wired Digital Lynx SX system, a single computer handled both neural acquisition and behavioral tracking through the Cheetah software (Neuralynx, Inc.); in those sessions, head position and 2D head orientation were extracted from two LEDs mounted on the headstage.

### Histological identification

After the final recording sessions, rats were transcardially perfused with 3.7% formalin. To improve the quality of tetrode tracks, the brain was partially exposed and soaked in PFA or formalin overnight with the hyperdrive attached before tetrodes were retracted. The brain was extracted and stored in 30% sucrose-formalin solution until fully submerged, and sectioned sagittally at 40 µm intervals. The sections were mounted and stained with 0.1% cresyl violet.

For tetrode track reconstruction, Nissl-stained serial sections were imaged on a light microscope, and whole-section images were acquired with IC Capture 2.1. Images were imported into Free-D (*108*) for manual tracing of tetrode tracks across consecutive sections and 3D reconstruction of each tetrode’s trajectory. Reconstructed trajectories were visualized from the top-down and registered to the known hyperdrive bundle geometry to assign tetrode identities. For each identified tetrode, the 3D reconstructed trajectory was exported as an STL file for further analysis in MATLAB to represent the track as a 3D curve and to compute arc length (distance along the curve) at each point.

Recording depths were logged for each tetrode in each session as cumulative turns of a 0-80 hyperdrive screw from its initial position at implantation. To map these depths onto the reconstructed tracks for each tetrode, we used the most ventral point of each reconstructed track as a reference corresponding to the deepest recording depth, defining arc length as the distance measured along the track from this point. Screw-turn values of each tetrode were converted to expected tetrode displacement using the 0-80 screw pitch (317.5 µm/turn) and adjusted for ∼20% tissue shrinkage, yielding an arc-length position along the reconstructed trajectory for each session’s recording depth. For each tetrode, these arc-length positions were then converted to a specific histological section and an (x, y) coordinate within that section using the microscope’s pixel-to-micron scale (µm/pixel) for that imaging magnification.

For each tetrode, the regional identity of the recording location in each session was determined by examining the corresponding (x, y) location within the identified histological slice under high magnification based on cytoarchitectonic features and anatomical (e.g., sulcal) landmarks. Borders involving area 29e, PR and PA were identified using Haug’s Silver-sulfide stained sagittal sections (*71*) as a reference, whereas borders involving MEC, PA, and PO were identified following previously published parahippocampal and entorhinal anatomical boundaries (*22*, *109*) and a Long-Evans rat atlas from Menno Witter and colleagues.

High-resolution images of histological sections (Fig. 1A; fig. S1) were acquired on a KEYENCE BZ-X1000 fluorescence microscope, capturing six brightfield tiles (2×3) with a 4x objective and stitching them into a single mosaic with the BZ-X Analyzer stitching & multi-point imaging module software.

### Spike sorting

Wideband extracellular recordings were band-pass filtered between 300 and 3000 Hz to isolate spiking activity, whitened, and artifact-masked before automatic spike detection and clustering with MountainSort4 (*110*). Outlier events within each cluster were identified in the peak–amplitude feature space of the spike waveforms using MATLAB’s *robustcov* function and removed, thereby reducing residual contamination from other sources (*111*). The cleaned clusters were transferred to *Phy* via the *SpikeInterface* framework (*112*) for further manual curation based on spike waveforms, amplitude stability over time, and the distribution of waveforms in principal-component feature space. Inter-spike-interval histograms and pairwise cross-correlograms were also examined to decide whether clusters should be merged or split: clusters were split when mixed waveform populations were evident and merged when two clusters displayed nearly identical waveforms together with a pronounced refractory trough in their spike cross-correlogram. Units showing clear amplitude drift or more than 1% of inter-spike intervals shorter than a 1.5 ms refractory period were discarded. The remaining well-isolated, temporally stable clusters were retained for all subsequent analyses. In a subset of sessions, MountainSort4 units were compared to manually sorted clusters using custom software (WinClust, J. J. Knierim) and were found to achieve comparable isolation quality, as indicated by similar waveform features and refractory-period violations.

### Selection criteria for independent neuron and LFP samples

To avoid including the same neuron more than once in an analysis, we employed a deliberately conservative, depth-based duplicate-control procedure. Two clusters recorded on different days were judged to be distinct only if the tetrode had been advanced by at least three ⅛-turn screw steps(∼119 µm). For each analysis, clusters from all sessions were first pooled by rat and tetrode; the largest subset of clusters that satisfied this constraint was then chosen through an optimization algorithm. The same depth-based constraint was applied to LFP signals: LFPs from the same region were considered as different samples only if they were recorded at least 119 µm apart. Mixed-effect models were used to account for the nonindependence of LFP signals from the same animal. See relevant sections for details.

### Machine learning-assisted physiological recording region identification for ambiguous histological cases

*Pipeline architecture*

In cases where the histological brain-region assignment of a tetrode was ambiguous or infeasible due to histological imperfections, we developed a machine-learning (ML) pipeline to classify the recording region, based on the firing properties of all neurons recorded on that tetrode (fig. S2B). The pipeline was designed to ensure high confidence assignments (i.e., to minimize incorrect assignments to a particular brain region) at the possible expense of classifying neurons as ambiguous. Because histologically ambiguous tetrode recording sites were most often located near the borders of area 29e or PR with neighboring regions, we trained two separate binary classification pipelines: area 29e vs. rest (i.e., MEC, PA, PO and PR) and PR vs. rest (i.e., 29e, MEC, PA, PO). For each pipeline, the “positive class” was defined as neurons that were unambiguously assigned to the target region (area 29e or PR), while the “negative class” included the remaining neurons unambiguously assigned to all other regions. Training data for all pipeline components consisted exclusively of neurons from recording sites with unambiguous histological region assignments (i.e., ground-truth data); neurons from histologically ambiguous recording sites were never used for model development or its performance verification.

The first layer of each pipeline (i.e., 29e vs. rest and the PR vs. rest) comprised an XGBoost model (a boosted decision-tree algorithm; *83*) that performed brain-region classification using the physiological features of single neurons. Features included: (i) spike-waveform characteristics: the peak-aligned normalized waveform, spike width, and spike asymmetry (trough-to-peak amplitude ratio); (ii) theta-modulation measures: the mean resultant length (MRL) of spike–theta phases (*50*) and three metrics quantifying theta rhythmicity (*50*, *51*, *36*); (iii) firing-pattern features: mean firing rate, burst ratio (proportion of inter-spike intervals < 8 ms), burst index and refractory period (*114*); and (iv) spike-autocorrelogram (ACG) properties: mean and mode lag of the ACG, and the power spectrum of the spike ACG (1–90 Hz). After eliminating highly correlated features in this set, a four-fold cross validation routine was employed for final feature selection and optimization of model hyperparameters within the training data. Classification probabilities from the resulting XGBoost model were then calibrated with Platt scaling (*115*, *116*) using an additional cross-validation procedure within the training data.

The second layer employed stacked generalization (*117*), a meta-model that classified the brain region of a tetrode by combining the single-neuron probability outputs from the first classification layer across all simultaneously recorded neurons on that tetrode. To ensure the meta-model was invariant to the presentation order of neurons, we first transformed single-neuron probabilities from the first layer into a permutation-invariant feature representation using symmetric aggregations. e.g., sum, sum-of-cosines (*118*). These permutation-invariant representations served as input to a logistic regression meta-model. We trained this meta-model with a synthetic dataset generated by random subsampling of the positive and negative classes in the training data for each pipeline (i.e., the pipeline for 29e vs. rest or PR vs. rest classification). Hyperparameter and Platt scaling-based probability calibration for the meta model were optimized using cross-validation on this synthetic training dataset.

The final layer applied independent confidence thresholds to the output probabilities of the meta-model in each pipeline for both positive (“29e” or “PR”) and negative (“not 29e” or “not PR”) class assignments—a technique known as classification with a reject option (*119*, *120*). Region assignments were accepted only when the output probability for either class exceeded its respective threshold; otherwise, no assignments were made to ensure that only high-confidence assignments were accepted. Thresholds for each pipeline were optimized via cross-validation on the training data: after meta-model fitting on each fold’s training data, a range of threshold values was scanned on validation data to quantify both coverage (the fraction of accepted assignments) and accuracy among accepted assignments. The resulting trade-off between coverage and accuracy was visualized as a Pareto front (*121*, *122*) from which threshold values were manually selected to provide an optimal balance between these metrics for both positive and negative class assignments.

*Pipeline verification*

To assess pipeline performance on unseen data, we performed tetrode-stratified, five-fold cross-validation (CV) on the histological ground-truth dataset. We refer to this as external CV to distinguish it from the internal CV procedures used by the pipeline for tuning its parameters within the training data, as described above. In each fold of the external, tetrode-stratified CV, the dataset was split into training and test sets in a “tetrode-aware” fashion, i.e., by leaving out all data from selected tetrodes. This split ensured that neurons recorded on the same tetrode did not appear in both sets, preventing potential data leakage due to the correlated nature of single units recorded on the same tetrode. All tuning and calibration procedures of the pipeline (e.g., feature selection, hyperparameter optimization, probability calibration, and reject-option threshold selection) described above were carried out using internal CV restricted to the training set within each fold of the external CV.

We then evaluated the resulting pipeline on the test set of the external CV, comprising neurons from held-out tetrodes. To quantify performance as a function of the number of neurons available for tetrode-level regional classification, we generated synthetic test samples by pooling neurons across held-out tetrodes within each class (positive or negative) and then randomly subsampling N neurons (N=1, 2, …,7) from the appropriate pool. This approach provided sufficient test cases at each N, allowing us to quantify each pipeline’s sensitivity and specificity for identifying the respective positive class (“area 29e” or “PR”) as a function of N (fig. S2D). The sensitivity and specificity of both 29e and PR models increased as N increased.

*Pipeline application*

After verifying pipeline performance via tetrode-stratified cross-validation (fig. S2D), we trained the final versions of each pipeline (29e vs. rest and PR vs. rest) on the complete histologically ground-truth dataset. For these final models, all tuning procedures—feature selection, hyperparameter optimization, probability calibration, and reject-option threshold selection—were performed via internal cross-validation on this full dataset, following the same methodology described under *pipeline architecture*. Note that performance metrics reported in fig. S2D were obtained exclusively from the tetrode-stratified verification CV, not from this final refit. The finalized pipelines were then applied to classify the recording regions of tetrodes with ambiguous or infeasible histological localization.

Since classification performance (sensitivity and specificity) improved as the number of simultaneously recorded neurons increased (fig. S2D), we set minimum thresholds for the number of simultaneously recorded cells per tetrode based to ensure that the pipeline was applied only under conditions of high reliability (both sensitivity and specificity ≥ 0.85). Specifically, we applied the 29e vs. rest pipeline to ambiguous tetrodes with at least 5 neurons, and the PR vs. rest pipeline to ambiguous tetrodes with at least 7 neurons. Assignments were accepted only when the model’s output probability exceeded the confidence threshold for either the positive or negative class; otherwise, no assignment was made. All ML-based region assignments for ambiguous tetrodes were finalized prior to downstream data analyses, ensuring that classification decisions remained blinded to experimental outcomes.

### Inclusion criteria

During preprocessing, we enforced a common set of inclusion criteria across analyses. Unless otherwise specified, spikes were analyzed only when the animal’s running speed exceeded 4 cm/s and landmarks were visible. Units were included if they produced ≥ 100 spikes within the relevant analysis epochs.

Only clusters that could be unequivocally assigned to MEC, PA, PO, PR, or 29e—by histological examination, or, when histology was ambiguous, by our physiological-feature classifier—were retained (see “*Brain region classification from single–cell physiological features*”).

For tetrode–level landmark–OFF response analyses (Fig. 3A, B), tetrodes were included only if the population mean firing rate was ≥ 2.5 Hz during the analyzed period. For cross-regional LFP interaction analyses, a tetrode was included only if at least one neuron with spatially correlated activity was recorded during the period with stationary landmarks in the Dome VR. For monosynaptic–connection analyses, putative monosynaptic pairs were considered only if both presynaptic and postsynaptic units had average firing rates ≥ 0.2 Hz.

For open-arena analyses were restricted to sessions that were ≥ 20 min. To ensure sufficient coverage during running, we required that in every square bin of a coarse 4 × 4 grid tiling the arena, the animal had spent at least 75 seconds over the full session and at least 30 seconds within each half of the session.

For cross-regional LFP–LFP and spike–LFP coupling analyses (see below), we used tetrodes that had at least one neuron with spatially correlated activity, detected based on the initial epoch (G = 1) of the Dome VR experiments with stationary landmarks.

### Neuron matching between Dome and open arena recording sessions

Putative neurons (clusters) were matched across Dome and open arena recording sessions performed on the same day based on the similarity of their spike waveform features, as in (*123*). Peak waveform voltage features from the tetrode channels were extracted for each cluster and modeled as multivariate Gaussian distributions. The ‘cluster distance’ between each pair of clusters across sessions was quantified as the minimum of the two asymmetric Kullback-Leibler divergence values between their Gaussian distributions. For each cluster in a session (i.e., Dome or open arena), we selected the cluster in the other session with the minimum cluster distance as its putative match. To ensure confidence in this match, we checked whether any other candidate cluster had a distance falling within 0.5 of this minimum distance: if so, we treated the match as ambiguous and discarded it, leaving the cluster without a match in the other recording session. For each cluster that retained a putative match after this confidence check, we additionally compared the putative match’s cluster distance against a threshold to reject poor matches. To optimize this threshold, we followed a procedure which divided sessions in half to create ground-truth same-unit pairs and then searched for a threshold that balanced matching accuracy with rejection rate. This procedure identified a threshold of 8, which we applied to reject all Dome– open-arena cluster pairs with distances exceeding this value. Finally, we only kept matches that were bidirectional, i.e. each cluster had to choose the other as its unambiguous match.

### Statistics

Statistical analyses were performed in MATLAB (MathWorks, Inc.). All tests were two-sided and evaluated at a familywise α = 0.05 unless otherwise indicated. Whenever multiple comparisons were performed, *p*-values were adjusted using the Holm–Bonferroni step-down method to control the familywise error rate.

Nonparametric tests (Kruskal-Wallis test if number of groups > 2, Wilcoxon rank-sum test otherwise) were used for group comparisons unless the data satisfied the following parametric assumptions.

Normality was assessed with the Shapiro–Wilk test (*124*), except for groups with sample kurtosis > 3, where we used the Shapiro–Francia test, which has greater power to detect non-normal distributions with more extreme values or heavy tails (*125–127*). Homogeneity of variances across groups was examined with both Levene’s test (absolute-deviation variant) and Bartlett’s χ² test (MATLAB’s *vartestn* with options ‘LeveneAbsolute’ and ‘Bartlett’). If every group passed the normality test and either one of the tests for equal variances were non-significant (*p* > 0.05), we treated the data as suitable for parametric analysis. We ran a one-way ANOVA to test for overall group differences, checked that the ANOVA residuals were still normal, and then compared groups pairwise with standard two-sample t-tests. For two group unpaired comparisons, we used the nonparametric Wilcoxon rank-sum test unless the data met parametric assumptions. If both groups were normally distributed, we tested equality of variances with the two-sample F-test; when variances were equal we used the standard two-sample t-test, and when variances differed we used the Welch’s t-test. For paired data, we analyzed the within-pair differences: if these difference scores were normally distributed, we applied a paired t-test; otherwise, we used the Wilcoxon signed-rank test.

To account for non-independence among LFP measurements from the same animal (and, where applicable, repeated measurements from the same tetrode pair, neuron, or brain region), we employed linear mixed-effects models. Following established practices for inference in linear mixed-effects models (*128*, *129*), models were fit by restricted maximum likelihood (MATLAB’s *fitlme*) and statistical significance of fixed effects was evaluated using Satterthwaite-approximated denominator degrees of freedom (MATLAB’s *anova* and *coefTest* functions on fitted mixed-effects models). This approach yields accurate *p*-values, when the number of animal is relatively small and/or when the effective sample size is closer to the number of animals than to the total number of observations, because repeated measurements from the same animal are correlated. In contrast, alternative approaches that assume large-sample asymptotics (e.g., Wald/z-tests) or use residual degrees of freedom (e.g., classical regression/t-tests) can yield inaccurate, optimistically low p-values by overstating the amount of independent evidence under these conditions.

Using Satterthwaite-approximated degrees of freedom, we evaluated statistical significance of fixed effects as follows: Fixed effects spanning more than two conditions (e.g., differences among more than two brain regions) or interaction terms were assessed via F-tests on the fitted model (MATLAB’s *anova*). All other tests on fixed effects, including single coefficients vs. zero (e.g., a condition mean or the slope relating two variables) and post-hoc pairwise comparisons, were evaluated via t-tests on linear contrasts of the fixed-effects coefficients from the fitted model (MATLAB’s *coefTest*). Model-specific details are provided in the relevant sections below.

### Power spectral analysis of LFP

*1/f background correction*

The wavelet transform was applied to each tetrode’s LFP signal using Morlet wavelets spanning 1–140 Hz (Buzsáki Lab, GitHub: https://github.com/buzsakilab/buzcode). Instantaneous power was computed from the wavelet coefficients and averaged across the session. To isolate oscillatory power from the aperiodic background in each LFP power spectrum, we used the FOOOF algorithm (*130*) with the following parameters: knee mode, peak threshold = 2.5, peak width = 2–45 Hz, and max_n_peaks = 4. Starting from the original spectrum, FOOOF was run iteratively as suggested in (*131*). In each iteration, FOOOF took the spectrum from the previous iteration as an input and fitted a parametric model for the oscillatory peaks and the aperiodic (1/f) background. Subtracting the identified peaks from this spectrum resulted in a refined spectrum, which was then used as input for the next iteration. This process continued until no oscillatory peaks greater than 1% above the aperiodic background remained, or until a maximum of ten iterations. The final peak-removed spectrum was then fit with a simple log–linear model to robustly estimate the aperiodic background. This fitted background was subtracted from the log-transformed original spectrum to obtain the 1/f-corrected (flattened) log power spectrum for each tetrode. Positive values in the 1/f-corrected log power spectrum indicate frequencies where the power exceeded the modeled aperiodic background, suggesting presence of oscillatory components, while values near or below zero indicate frequencies without a detectable oscillatory excess.

*Cluster-based permutation testing of significant oscillatory power across frequencies*

We assessed the statistical significance of oscillatory components in the LFP signal across tetrodes within each brain region while controlling for familywise error (due to multiple comparisons). A naïve Bonferroni correction would not account for the continuous (correlated) nature of the frequency spectrum, so we opted for a cluster-based permutation test (*53*, *132–134*). For each permutation, the sign of the entire 1/f-corrected log power spectrum from each tetrode was randomly flipped for a given session. At each frequency, a one-sample, one-sided t-test was performed across tetrodes for both the observed and permuted values. Clusters were defined as contiguous frequencies ranges over which there were significant positive deviations from zero according to *p* < 0.05 (one-sided t-test). The size of each cluster was calculated as the sum of t-statistics within the cluster. A null distribution was created from the maximal cluster size from each of the 1000 permuted data sets. Each cluster in the observed data was then compared to this null distribution, yielding permutation-based cluster-level *p*-values. Clusters that satisfied *p* < 0.05 for this permutation test were considered significant.

*Mixed effects-model of theta- and gamma-band LFP power*

Average powers in theta (5–10 Hz) and gamma (50–120 Hz) bands were calculated from the 1/f-corrected log-power spectrum of each tetrode to test for regional differences. We used 1/f-corrected power spectra rather than raw power due to potential differences in aperiodic background activity across days and/or animals, which were not perfectly balanced across recorded regions. To also account for non-independence of LFP signals within animals when comparing band-limited power across regions, we used linear mixed-effects modeling.

For each frequency band (theta and gamma), data were arranged as one observation per tetrode location with the following variables: band-limited 1/f-corrected power, region identity (categorical variable with five values: 29e, MEC, PA, PO, and PR), and rat identity (categorical variable). Region identity was included as a fixed effect to test for systematic differences in power across regions. The model included two random-intercept terms: a rat-specific random intercept (1|rat) to capture correlation among LFP observations from the same animal, and a rat-by-region specific random intercept (1|rat:region) to capture additional correlation among LFP observations from the same region within the same animal (i.e., allowing each rat to have a region-specific baseline). For each frequency band, we then fit the following mixed-effects model for each band (theta or gamma):

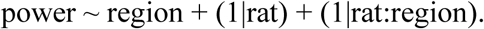

Each power band model was fit by restricted maximum likelihood (using MATLAB’s *fitlme*) and the statistical significance of fixed effects (i.e., both omnibus test of region and post-hoc pairwise contrasts between regions) was evaluated using Satterthwaite-approximated denominator degrees of freedom.

Omnibus effect of region was assessed with F-test using MATLAB’s *anova* function on the fitted mixed-effect models. When this omnibus effect was significant, we performed post-hoc pairwise comparisons between each region and 29e using t-tests on linear contrasts of fixed effects’ coefficients from the fitted model (implemented via MATLAB’s *coefTest*). *P*-values across the four region-versus-29e contrasts for each model were adjusted for multiple comparisons using the Holm–Bonferroni correction.

### Theta and gamma rhythmicity of single neurons

Rhythmicity was assessed from power spectrum of each neuron’s spike autocorrelogram as follows: the theta index was calculated as the mean power in 5–10 Hz divided by the mean power in 1–50 Hz (*50*), and the gamma index as the mean power in 50–120 Hz divided by the mean power in 5–250 Hz (*135*).

### Coupling of single neurons to LFP theta rhythm

Each tetrode’s LFP signal was band-pass filtered between 5–10 Hz (4^th^ order Butterworth filter). The Hilbert transform was estimated using MATLAB’s *hilbert* command, which returns a complex signal called the analytic signal. Instantaneous theta-band power and phase were extracted from the magnitude and angle of the analytic signal, respectively. For each neuron, the spike-phase distribution was obtained by circularly interpolating the instantaneous theta phase from each tetrode’s LFP signal at the neuron’s spike times. Pairwise phase consistency (PPC) was then computed for each neuron’s spike-phase distribution relative to each tetrode using FieldTrip (*136*). The highest PPC across tetrodes was used to quantify each neuron’s coupling to the LFP theta rhythm.

### Spatially correlated activity in the Dome

A circular firing-rate map was created for each neuron to quantify its spatial tuning in the Dome when landmarks were stationary (i.e., experimental gain, G = 1). We assigned both the angular position of each spike and the animal’s position at each behavioral time point to 36 angular bins (10° per bin) spanning all locations across the circular track. We then computed the firing rate in each bin by dividing the number of spikes by the amount of time spent in that bin. The resulting rate maps were smoothed with a circular Gaussian kernel (σ = 15°). The smoothed circular rate map was then used to calculate each neuron’s Skaggs’ spatial information (SI) score (in bits per spike; *38*):

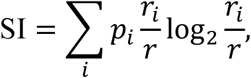

where *p*_*i*_ is the fraction of total session time the animal spent in *i*^th^ angular bin, *r*_*i*_ is the neuron’s firing rate in the same bin, and *r* is the neuron’s overall mean firing rate; the summation is over the bins, *i* = 1, …,36. Because this score (originally developed for 2D firing-rate maps) can produce both false-positive and false-negative errors when applied to data from 1D circular tracks (*137*), we also quantified spatial tuning using two alternative SI metrics, the Olypher SI score and the mutual SI score, defined below.

The Olypher SI score is defined as the maximum Kullback-Leibler divergence between the neuron’s spike count distribution at any given position and its overall spike count distribution (*138*):

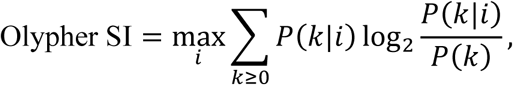

where *P*(*k*|*i*) is the probability of observing *k* spikes within a 100 ms time-segment when the animal occupies positional bin *i*, and *P*(*k*) is the marginal probability of observing k spikes, regardless of the animal’s position.

The mutual SI score is defined as the mutual information between the neuron’s spike count and the animal’s position (*139*, *140*):

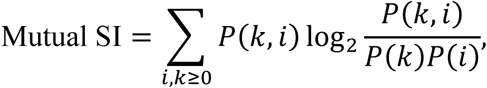

where *P*(*i*) is the marginal probability of the animal occupying the positional bin *i* within a 100 ms time segment (independent of the number of spikes), *P*(*k*) is the marginal probability of observing *k* spikes within a 100 ms time segment (independent of occupancy), and *P*(*k*, *i*) is the joint probability of observing *k* spikes and the animal being in positional bin *i* within the same 100 ms time segment.

To assess significance of each SI score for each neuron, we generated a null distribution by circularly shifting the neuron’s spike train relative to behavioral data by a randomly chosen time offset (uniform distribution between 25 and T – 25 seconds, where T is the total duration of the first experimental epoch) and recalculating the SI scores 100 times. A neuron was considered to have significant spatially correlated activity if at least one of its observed SI scores exceeded the 99^th^ percentile of the respective shuffled SI distribution. We refer to such neurons as “spatially correlated”.

### Landmark-selective firing modulation in the Dome

*Z-index*

The circular firing-rate map was linearly interpolated to 1° resolution (0–359°) and normalized to z-scores across angular bins. For each landmark location, we identified the angular bin corresponding to the circular track position closest to the landmark. We then computed the mean z-scored firing rate within a symmetric, 24° angular window centered on that bin (i.e., ±12° around the landmark angle, wrapping circularly as needed). For comparison, we identified midpoint angles halfway between each pair of adjacent landmarks and computed the mean z-scored firing rate within an identically defined 24° window centered on each midpoint. The landmark selectivity z-index was then defined as the absolute difference between the mean z-scored firing rate averaged across all landmark-centered windows and that across all midpoint-centered windows.

*Local similarity index*

Each neuron’s circular firing rate map was linearly interpolated with 1° resolution and partitioned into sectors around each landmark. Each landmark sector was defined as the arc spanning between the midpoints of that landmark and its two neighboring landmarks. For example, if there were three equally spaced landmarks, there would be three 120° sectors, one centered on each landmark; however, if the landmarks were unevenly spaced (as was typical), the sectors would have different widths. For the given neuron and for each pair of landmarks, the Pearson correlation coefficient was computed between the peri-landmark firing-rate profiles, using only the angular range relative to the center of the landmarks shared by both sectors (i.e., their intersection). This shared range varied from as little as ±20° up to ±60° for different pairs, depending on the number and spatial arrangement of landmarks within the Dome (Fig. 1B). For two landmarks, there was one such pairwise correlation, for three landmarks there were three pairwise correlations, and four landmarks there were six pairwise correlations, based on the formula *N(N-1)/2* where *N* is the number of landmarks. The local similarity index for each neuron was then defined as the average of all such pairwise correlations across landmark pairs.

### Landmark-locking score

For each neuron, we computed a lapwise firing-rate map in the landmark frame by binning spikes and the animal’s behavioral data within each lap into 10° angular bins, dividing the spike count in each bin by the time spent in that bin, and smoothing with a Gaussian kernel with σ = 15°. Using the last 12 laps in each session of this firing-rate map, we calculated the spatial autocorrelation function for spatial lags between 0 and 9 laps. (Thus, the minimum overlap in the autocorrelation calculation was 3 laps, corresponding to a lag of 9 laps within the last 12 laps.) Neurons with stable lapwise firing exhibited autocorrelation functions with prominent oscillations at integer-lap periods, indicating consistent periodicity across laps (Fig. 1E; fig. S2A), whereas neurons that broke free from landmarks displayed autocorrelation functions with weak or irregular oscillatory structure (Fig. 1F; fig. S2B).

To quantify the lap-to-lap periodicity of each neuron’s autocorrelation function, we first computed all possible pairwise Pearson correlations among circularly seven shifted versions of the 9-lap autocorrelation function at full-lap lags (circular shifts of 0, 3, 4, 5, 6, 7, and 8 laps, leading to 21 pairwise Pearson correlation values amongst these autocorrelograms) as well as all possible correlations between these full-lap shifted versions and the half-lap rotated versions (five circular shifts from 3.5 to 7.5 laps, in 1-lap steps, leading to 35 pairwise Pearson correlation values between full-lap and half-lap shifted autocorrelograms). Neurons with stable lapwise periodicity are expected to show higher correlations among the full-lap rotations than between the full-lap and half-lap groups.

Accordingly, the landmark-locking score was defined as a normalized difference measure: the sum of all Pearson correlations within the full-lap rotation group minus the sum of all correlations between the full-lap and half-lap groups, divided by the total number of correlations (Fig. 2; fig. S2). A score close to 1 indicates strong, consistent periodicity at the spatial scale of one lap, while a score near zero indicates little or no periodic firing structure.

### Spatial tuning in stationary and mobile segments of open-loop sessions

The decoupling of landmark rotation from animal movement in open-loop experiments created periods where landmarks rotated while the animal remained stationary. Supplementary analysis (fig. S4) examined whether neural firing patterns relative to landmarks were preserved during these *stationary* periods. Accordingly, we defined two conditions from data where the animal’s speed relative to landmarks was > 4 cm/s: *mobile* and *stationary. The mobile* condition corresponded to periods when the animal’s lab-frame speed was also ≥ 4 cm/s (regardless of landmark motion), whereas the *stationary* condition corresponded to periods when the animal’s lab-frame speed was < 4 cm/s (meaning landmarks moved relative to the stationary animal).

The discontinuous nature of *stationary* segments precluded use of the landmark-locking score—which requires continuous spatial sampling across laps—to quantify the extent of landmark-anchoring during these segments. Instead, we assessed spatial tuning during *stationary* segments by comparing landmark-referenced firing patterns between *stationary* and *mobile* segments. We restricted this analysis to “landmark-anchored” neurons—defined as those with landmark-locking scores > 0.52 (the threshold separating *LM-control* and *LM-failure* sessions in Fig. 1G)—to ensure that anchoring to landmarks during stationary segments were evaluated only for neurons with stably anchored firing fields to the rotating landmarks during *mobile* periods.

For each landmark-anchored neuron, we computed the 1D circular firing rate map as a function of angular position relative to landmarks (10° bins) for both *mobile* and *stationary* periods. Pearson’s correlation between the two rate maps quantified the *mobile–stationary* similarity of the rate maps. To determine the statistical significance of the observed similarity for each neuron, we circularly shifted the *stationary* rate map by 2 to 34 bins and computed its Pearson correlation relative to the *mobile* rate map, yielding a shuffled distribution of correlation coefficients. Landmark-anchored neurons with observed correlations exceeding the 95^th^ percentile of its shuffle distribution were considered statistically significant in terms of similarity of its rate maps across *mobile* and *stationary* conditions. Due to limited number of landmark-anchored neurons from MEC, PA, PO, and PR in these experiments, neurons from these regions were pooled into a single “non-29e” group for comparison with area 29e.

### Unsupervised identification of LM-control and LM-failure session types

To identify session-wise differences in landmark control over neuronal populations, we first computed the average landmark-locking score across all recorded neurons from all regions in each session.

Sessions were then clustered based on their respective average landmark-locking score using an enhanced density-based clustering algorithm (MATLAB’s *dbscan*) procedure that set MinPts = 4 and automatically tuned the neighborhood radius ε through k-distance heuristic with k = MinPts (*141*).

Briefly, the absolute difference between the average landmark-locking score of each session and that of all remaining sessions was calculated. We then measured, for each session, the distance to its k^th^-nearest neighbor session. Measurement of this distance for each session led to the so-called k-distance graph.

The neighborhood radius ε was set as the distance corresponding to the knee point of this graph. DBSCAN was run with this ε and the corresponding MinPts. Any session labeled as noise (i.e., left out of clustering) was assigned to the cluster whose centroid was nearest. This unsupervised approach identified two clusters which we defined as LM-control and LM-failure. See Fig. 1G.

### Landmark removal score

Multi-unit, unclustered spike trains from each tetrode were binned at 10 ms resolution. To quantify the neural response to landmark removal, we aligned spike counts to the time of landmark extinction and extracted a window from –1 to +1 seconds relative to this event for each session.

The pre-removal baseline was defined as the period from –0.6 to 0 seconds before landmark removal, and the post-removal response period as 0 to +0.4 seconds after removal (the larger pre-removal time period was used to get a better estimate of the intrinsic variability before removal; the result does not change when symmetric 400 ms pre-and post-removal time periods were used). For each time bin, we calculated the absolute deviation from baseline as *d*(*i*) = | spike count in bin *i* – mean baseline spike count |. The landmark removal response score was then computed as:

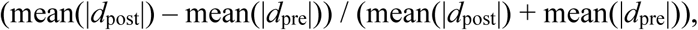

where the means are taken over all 10ms time bins in the pre- or post-removal period; thus, mean(|*d*_pre_|) and mean(|*d*_post_|) represent the mean absolute deviations from baseline during pre- and post-removal periods, respectively.

This absolute deviation approach treats increases, decreases, and even biphasic responses (i.e., a combination of increase and decrease) equivalently. The normalized score ranges from –1 to +1, with positive values indicating that post-removal firing deviates more from baseline than expected from pre-removal fluctuations (suggesting a response to landmark removal), while values near zero indicate deviations comparable to baseline fluctuations.

### Single neuron responses to landmark and background luminance manipulations

Lapwise firing rate maps for each neuron were computed as before (see Landmark-locking score section). Landmark and background luminance were each set to one of two levels (high/low), yielding four landmark–background luminance combinations (2 × 2 factorial design).

*Luminance sensitivity analysis*

We fitted a two-way ANOVA to each neuron’s lapwise peak firing rates, using lapwise landmark luminance (LM) and background luminance (BG) as factors. The effect size for each factor X ∈ {LM, BG} was quantified by

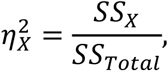

where *SS*_*X*_ denotes the sum of the squares in peak firing rate explained by the variation in the factor *X* and *SS*_Total_ denotes the total sum of squares in the peak firing rates. Because these luminance factors were independently manipulated and uncorrelated, we quantified their combined overall effect size as the sum 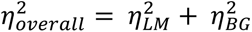.

Although landmark and background luminance were independently manipulated by a two-factorial design, the resulting contrast (C=LM – BG) and total luminance (TL=LM + BG) values were linearly dependent and did not obey a fully factorial design, precluding the use of ANOVA to analyze firing-rate dependence on (C, TL). Accordingly, a linear model was fit with C and TL as predictors and each neuron’s peak firing rates as the dependent variable. The unique contribution of each predictor X ∈ {C, TL} was quantified using partial R²:

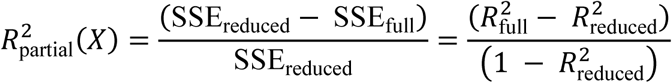

Here, SSE is the sum of squared errors, “full” refers to the model with both predictors, and “reduced” refers to the model with X removed. Partial R² thus quantifies the additional variance in firing rate explained by a predictor, after accounting for the presence of the other, even when the predictors are correlated.

### Allocentric place and head direction tuning in the open arena

For robust characterization of allocentric place and HD tuning scores despite differences in behavioral sampling of place and HD, we used a Poisson generalized linear model (GLM), as in previous work (*44*, *142*). Briefly, each neuron’s spike train was binned into *dt =* 125 ms segments. For each segment *t*_*i*_, the expected spike count was modeled as

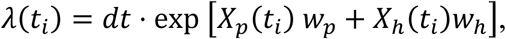

where *X*_*p*_ and *X*_ℎ_ are one-hot encoded row vectors representing, respectively, the animal’s (x,y) position (in 7 × 7 cm^2^ bins) and HD (in 8° angular bins) at the center of that segment. The column vectors *w*_*p*_ and *w*_ℎ_ denote the predictive weights for position and HD (analogous to tuning curves). Fitting of these weights was performed by maximizing an objective function that combined the Poisson log-likelihood of the observed spike counts with a smoothing penalty to discourage large weight differences between adjacent bins, as previously described (*44*, *142*). Optimization was performed using the MATLAB’s *fminunc* search routine, aided by analytically computed gradients. Using the optimized weights *w*^_*p*_and *w*^_ℎ_ for each neuron, we constructed a 2D spatial firing rate map and a 1D HD tuning curve, each marginalized over the other behavioral variable:

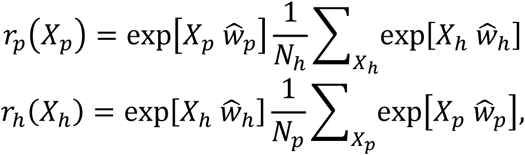

Here, *N_h_* = 72 (the number of 5° HD bins) and *N_p_* = 35 × 35 (the number of 4 × 4 cm^2^ positional bins). These marginal rate maps were then used to calculate the Skaggs’ SI score in bits per spike as a measure of each neuron’s place-tuning strength and Rayleigh score (i.e., mean resultant length) as a measure of its allocentric-HD tuning strength.

*Testing significance and stability of allocentric place tuning*

We circularly shifted each neuron’s spike train by a randomly chosen time offset (between 25 and T – 25 seconds, where T is the total session duration) 100 times, refitting a GLM to each shuffled spike train to generate a null distribution for both SI and Rayleigh score.

For tuning stability of each neuron, we first computed the conventional (i.e., without using a GLM) 2D place rate maps and HD polar tuning curves from the first and second halves of the session, separately. The split-session stability of place tuning was then quantified as the Pearson correlation between the split-session rate maps, while the split-session stability of HD tuning was quantified as the difference in mean angular direction between the first and second halves of the session.

A neuron was considered significantly tuned to place if its GLM-derived SI exceeded the 99^th^ percentile relative to the shuffled distribution of GLM-derived SI scores and if its split-session stability exceeded 0.5. A neuron was considered significantly tuned to allocentric HD if its GLM-derived allocentric HD tuning curve passed the Rayleigh test of uniformity at α = 0.01, if its GLM-derived Rayleigh score exceeded the 99^th^ percentile of the respective GLM-based shuffled distribution, and if its split-session mean directional difference was less than 45° as previously used (*45*).

### Egocentric head-direction tuning in the open arena

Because egocentric coding can in principle be anchored to any arbitrary point in an environment that is unknown to the investigator, quantifying a neuron’s egocentric tuning curve typically involves an exhaustive search to determine its optimal anchor point. In contrast, allocentric coding is characterized by a single possible tuning curve per neuron, requiring no anchor-point search. If one compares the single allocentric tuning curve to the most specifically tuned egocentric tuning curve derived from the anchor-point search, this asymmetry creates a bias in favor of the egocentric model over the allocentric model when evaluating a neuron’s preferred frame of reference (*43*). Model selection metrics criteria (e.g., AIC/BIC) partly compensate for this bias but rely on assumptions that poorly capture the complexity of the anchor search and can still reward overfitting. More rigorous controls (e.g., surrogate tests) would help but are computationally prohibitive because each surrogate demands a new exhaustive anchor search.

To circumvent these issues, we developed an *egocentric vector-field* (EVF) score that quantifies egocentric tuning without requiring specification of an anchor point. The EVF score is computed as follows: We constructed a 2D HD vector field for each neuron by dividing the arena into a uniform 9 × 9 grid of (x, y) locations with each spatial bin being 15 × 15 cm². We identified all time points when the animal’s position fell within the bin and assigned the corresponding HDs to 12 angular bins (30° each). If any angular bin at a grid location had less than 0.2 seconds of occupancy, the spatial bin size at that location was increased iteratively until every HD bin reached this sampling threshold.

For the local HD tuning curve constructed at each (x, y) grid location, we computed its complex-valued first harmonic vector,

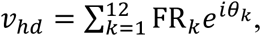

where FR_*k*_ is the firing rate in HD bin *k* associated with the angle *θ*_*k*_. Note that the angle of this vector is the same as the mean angular direction of the local HD tuning curve, while its magnitude is equal to the product of the summed firing rate within the local HD tuning curve and the MRL of the local HD-tuning curve. This vector is equal to the first (i.e., fundamental) Fourier component of the local HD-tuning curve, which defines the single sinusoid that best fits the tuning curve in the least-squares sense. The collection of these vectors across all grid locations comprised the 2D HD vector field for the neuron.

*Egocentric vector field (EVF) score*

To quantify egocentric tuning without specifying an anchor point, we analyzed the spatial derivatives of each neuron’s local HD vectors (i.e., how these vectors vary along the x and y axes of the arena). Specifically, we computed two classical vector calculus measures, *curl* and *divergence,* for each spatial bin (x, y) of the open arena (fig. S8).

The *Curl* of a vector field captures local rotational (swirling) structure of the HD vectors:

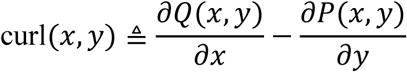

The *Divergence* captures the local inward/outward flow structure of the HD vectors:

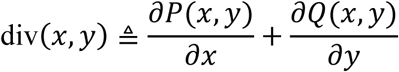

where, *P*(*x*, *y*) and *Q*(*x*, *y*), are the *x*- and *y*-components of the two-dimensional HD vector-field (corresponding to real and imaginary components of *v*_*hd*_ at each x,y location). These partial derivatives were estimated using centered-difference approximations between bin centers. Neurons lacking any systematic HD tuning and neurons with allocentric HD tuning (characterized by parallel vectors throughout the arena) exhibited near-zero curl and divergence everywhere (these neurons can be distinguished from each other by measuring the sharpness and significance of their allocentric directional tuning curves). By contrast, neurons with strong egocentric tuning generated spatially consistent, high curl and/or divergence values (fig. S8).

We combined the curl and divergence values at each spatial bin into a 2D vector. The curl was assigned as the horizontal component of this vector, whereas the divergence was assigned as its vertical component. This approach merged the separate curl and divergence fields into a new vector field that captures both rotational and flow characteristics of the preferred HD throughout the arena.

We then calculated the Egocentric Vector Field (EVF) score as the MRL of all divergence-curl vectors across the arena. This score ranges from 0 to 1, where values close to 0 indicate random or inconsistent curl/divergence patterns, whereas values close to 1 indicate highly consistent patterns characteristic of strong egocentric tuning.

*Generalized linear models for egocentric versus allocentric preference*

To dissociate each HD-tuned neuron’s preferred frame of reference (egocentric vs. allocentric) and to assess split-session stability of egocentric tuning, we used a Poisson GLM framework (*43–45*).

Specifically, for each neuron, we fitted two separate GLMs to its spike train, binned into 125 ms time segments: one GLM used allocentric head direction (HD) as a predictor, while the other used egocentric HD, computed relative to the (x, y) coordinates of a candidate anchor point. For the egocentric model, the optimal anchor point was identified by grid search: the GLM was re-fit at each candidate anchor location spaced every 5 cm across the arena in both x and y directions, and the location yielding the largest log-likelihood was selected.

This GLM-based anchor optimization served two purposes: (1) comparing predictive performance of allocentric versus egocentric models, and (2) identifying an anchor point to assess split-session stability of egocentric tuning (see below) and to compare arena-to-anchor point distance across regions (see Fig. 3G). Note that statistical significance of egocentric tuning was assessed separately using the anchor-free EVF score (below), which avoids the statistical biases inherent in anchor optimization as described above.

To compare the predictive performance of the two models while accounting for the increased complexity of the egocentric GLM (*K*_*ego*_ = 2 additional parameters corresponding to the anchor–point optimization), we calculated the difference in Bayesian information criterion

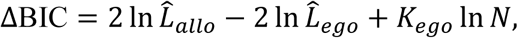

where *L̂*_*ego*_, *L̂*_*allo*_ are the optimal log-likelihoods of GLMs with egocentric HD and allocentric HD variables, respectively, and *N* is the number of bins (time segments). A positive ΔBIC suggests that the allocentric model provides a better fit, whereas a negative ΔBIC suggests a better fit for the egocentric model.

*Testing significance and stability*

To assess statistical significance of egocentric tuning, we generated two null distributions for the anchor-free EVF score of each neuron. Similar to other spatial tuning measures, one null distribution was obtained by circularly shifting the neuron’s spike train relative to the animal’s behavioral data by a single, randomly chosen time offset (between 25 seconds and T−25 seconds, where T is the total session duration), and recomputing the EVF score 100 times. The second null distribution was generated by disrupting the relationship between position and head-direction vectors in the HD vector field 1,000 times: for each iteration, we circularly translated the entire HD vector field along both the x and y axes by random amounts and then recomputed the EVF score.

To assess split-session stability of egocentric tuning, we constructed conventional (i.e., without using a GLM) egocentric-HD polar tuning curves from the first and second halves of the session. Both tuning curves were referenced to the optimal anchor point identified by the GLM grid search on the full session data (described above). Stability was quantified as the absolute angular difference between the preferred directions of the tuning curves from the two halves.

A neuron was considered significantly tuned to egocentric HD if its observed EVF score exceeded the 99^th^ percentile of both null distributions and the split-session mean directional difference was less than 45°.

### Classification of spatial and directional tuning of cells in open field

All neurons were evaluated for allocentric HD, egocentric HD, and allocentric place tuning as described above. Neurons meeting criteria for only egocentric-HD or only allocentric-HD tuning were classified accordingly. For neurons meeting criteria for both, we compared their egocentric and allocentric GLMs (i.e., the egocentric one corresponding to the optimal anchor point and the allocentric one) and classified each neuron according to the GLM that yielded the lower Bayesian Information Criterion score (BIC).

Results were unchanged when Akaike Information Criterion was used instead. Neurons failing to meet either HD tuning criterion were classified as nondirectional and screened for allocentric place tuning.

### LFP–LFP coherence

LFP signals were segmented into 1-second candidate windows with 95% overlap (0.05 s step size). These windows were evaluated sequentially from the start of the recording. Any window in which the animal’s velocity was below 4 cm/s for more than 25% of the time, or where the maximum velocity did not reach 10 cm/s, was discarded. If a window satisfied both speed criteria, it was retained for analysis. Other candidate windows that overlapped in time with this accepted window were excluded from further consideration. This process was repeated iteratively until no windows remained, resulting in a set of non-overlapping, speed filtered 1-second windows for the subsequent spectral analysis.

Within each retained window, LFP signals were determined. Spectral analysis was performed using the multi-taper Fourier transform with Slepian (DPSS) tapers via FieldTrip (*136*) for frequencies between 0 and 140 Hz with a frequency resolution of 1 Hz (as determined by the 1-second window size). For frequencies below 25 Hz, three tapers were used (yielding a frequency smoothing half-width of 2 Hz); for frequencies above 25 Hz, six tapers were used (smoothing half-width 3.5 Hz). For each window and frequency, the complex-valued cross-spectral density (CSD) matrix was computed from all simultaneously recorded tetrode pairs.

*Baseline coherence levels*

For each tetrode pair, we estimated the complex coherency (*C*) based on averaged bivariate CSD matrices across all retained, non-overlapping windows within the initial stationary-landmark epoch (G = 1). The magnitude of the resulting complex coherency estimate was biased upward due to a limited number of windows and has a variable variance that again depends on the number of windows. Under the null hypothesis of zero coherence, both bias and variance have analytically known forms (*143*, *144*) which can be used for bias correction and variance normalization as follows:

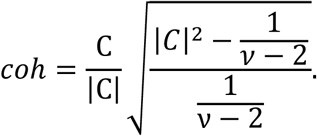

Here, ν denotes the degrees of freedom (equal to twice the product of the number of tapers and the number of windows), *C* is the raw coherency estimate, and *coh* is its bias-corrected and variance normalized version. To capture true (i.e., time-lagged, non-spurious) coupling between LFP signals while remaining insensitive to instantaneous shared signals arising from volume conduction or common reference effects, we focused on the imaginary part of this corrected coherency (i.e., *coh*), following previous work (*145–149*). Specifically, we took the absolute value of the imaginary component, normalized it by √1 − Re(coh)^2^—the theoretically maximum possible value of the imaginary component given the observed real component—and applied a Fisher z-transform, yielding what we refer to as the “z-imaginary coherence spectrum” (zCoh):

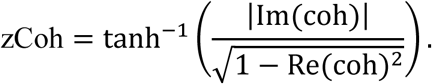

The resulting z-imaginary coherence spectrum at the single tetrode-pair level was then averaged across all 29e-MEC tetrode pairs, yielding a region pair-level average z-imaginary coherence spectrum.

To assess significance of the average z-imaginary coherence spectra across all 29e-MEC tetrode pairs, we used a hierarchical, cluster-based permutation test (*53*, *150*). For each tetrode pair, a null distribution of CSD values was generated by randomly flipping the sign of only the imaginary component of the CSDs from retained windows (which leaves the real part and magnitude unchanged), then recalculating the z-imaginary coherence spectrum after averaging the (imaginary) sign-flipped CSDs. This permutation was repeated 1,000 times per tetrode pair. To generate a group-level (region-pair) null, we randomly sampled one null spectrum from each of the above-mentioned individual tetrode-pair null distributions 5,000 times and, for each random sample, calculated the average z-imaginary coherence spectrum. The observed region pair-level z-imaginary coherence at a frequency was considered significant if it exceeded the 95^th^ percentile of the permutation group-level null distribution. Clusters were defined as contiguous frequencies that are significant with no gap. To control the familywise error rate of the comparisons across the whole frequency spectrum while respecting spectral contiguity, the cluster size (sum of coherence values within the cluster) was compared to the distribution of maximal cluster sizes from the group-level permutations. Clusters with corrected *p* < 0.05 were considered significant.

*Coherence change from initial to final epochs*

For each tetrode pair, we computed the z-imaginary coherence spectrum in the initial (G = 1) and final (G = G_final_) experimental epochs after averaging their bivariate CSD matrices across accepted 1-second, non-overlapping windows from that epoch. For each frequency band—theta (5–10 Hz) and gamma (50– 120 Hz)— and for each epoch (initial and final), we took the maximum z-imaginary coherence across frequency bins within the band and used this value as the band-specific coherence for that epoch. The normalized change in coherence for each band was then calculated as ΔCoherence = (final – initial)/(final + initial).

These within-session changes from the initial to final epoch were analyzed using linear mixed-effects models to account for repeated nature of theta- and gamma-band measurements from the same 29e– MEC tetrode pair and for the non-independence of measurements among multiple tetrode pairs recorded from the same animal.

Fixed effects included Session Type (*LM-control* vs. *LM-failure*), Band (theta vs. gamma), and their interaction (Session Type × Band). Because each 29e-MEC tetrode pair contributed a pair of observations (one for theta, one for gamma), and multiple tetrode pairs were recorded from the same rat, observations were not independent. We modeled this nested dependence structure using three random effects:

i. a rat-specific random intercept (1 | RatID), allowing variability of ΔCoherence across rats to capture correlation among observations from the same animal (across all tetrode pairs and both bands within each rat).
ii. a rat-specific random band effect (Band | RatID), allowing variability of theta-gamma difference across rats to capture covariance between theta and gamma-band observations from the same animal (across different tetrode pairs within each rat).
iii. a rat-specific random intercept for tetrode pairs (1 ∣ RatID:TetrodePairID), allowing repeated measurements from each tetrode pair (within a rat) to have its own baseline level of ΔCoherence, thereby capturing the correlation between theta and gamma measurements from the same tetrode pair.

The model was formulated as:

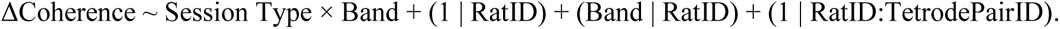

We fit the model by restricted maximum likelihood (using Matlab’s *fitlme* function) and evaluated statistical significance of fixed effects—including both omnibus tests on main and interaction effects as well as post-hoc comparisons—using Satterthwaite-approximated denominator degrees of freedom.

Omnibus effects were assessed with F-tests using MATLAB’s *anova* function based on the fitted model. When the Session Type × Band interaction was significant, post-hoc pairwise comparisons of simple effects (Control vs. Failure within each band; Theta vs. Gamma within each session type) were performed using t-tests on linear contrasts of the fixed effects from the fitted mixed-effects model (implemented via MATLAB’s *coefTest* function). Each condition mean (e.g., Control–theta vs. 0) was tested against zero using analogous linear contrasts on the same fitted model with the *coefTest* function. For each family of tests (four post-hoc pairwise comparisons and four vs-zero comparisons), *p*-values were adjusted using Holm–Bonferroni correction.

*Temporal relationship between ongoing changes in landmark-control strength and changes in LFP coherence*

The speed-filtered windows included in the spectral computation of LFP–LFP coherence were distributed unevenly relative to the animal’s unwrapped position (track angle) in the landmark frame of reference, because windows were selected over the time domain, while both running speed and landmark rotation varied over time. Consequently, consecutive accepted 1-second windows were typically separated by 30–90° (median 45°) yet collectively covered the full range of unwrapped landmark-frame positions within each session. To investigate the temporal relationship between ongoing changes in landmark control strength and changes in LFP–LFP coherence for every tetrode pair during a session, we resampled the CSD measurements from these unevenly distributed windows onto a uniform spatial grid (90° spacing) in the landmark frame of reference, using local linear kernel regression with a Gaussian kernel (135° standard deviation). The resulting spatially uniform, resampled CSDs were used to quantify the position-resolved z-imaginary coherence spectrum between each tetrode pair.

Landmark control-strength score was computed at the same spatial resolution as follows. For each 90° spatial bin, we calculated the mean firing rates of all spatially correlated neurons (excluding those from 29e) and constructed a population vector (PV). The full-lap PV correlation at each bin was calculated as the average Pearson correlation between the PV at that position and those from the same location one and two laps earlier (offsets of 360° and 720°). The half-lap PV correlation was calculated as the average correlation between the PV at that position and PVs from bins 180° and 540° earlier. At each spatial bin, the landmark-control strength was defined as (full-lap PV correlation – half-lap PV correlation)/2, where the division by 2 rescales the difference to a correlation-like, normalized range of – 1 to +1. The resulting, position-resolved, normalized landmark-control strength serves as a measure of landmark-anchored neuronal firing in non-29e populations throughout each session.

Prior to cross-correlation analysis, both the position-resolved landmark control-strength scores as well as z-imaginary coherence values were further smoothed with a Gaussian filter (240° standard deviation). At each frequency within theta and gamma frequency bands, we computed the Pearson correlation coefficient between the smoothed trajectories of landmark-control strength and z-imaginary coherence across multiple spatial lags ranging from –18 to 18 bins (± 4.5 laps). For each frequency band (theta and gamma), we identified the lag that maximized this correlation and recorded its corresponding correlation coefficient. We assessed the statistical significance of these band-specific correlation coefficients relative to null distributions obtained by circularly shifting the landmark control-strength trajectory by offsets ranging from 32 [8 laps] to N–32 bins, where N is the total number of spatial bins (typically

>200). For each shifted trajectory, we repeated the cross-correlation analysis and recorded the maximum correlation values within both theta- and gamma-bands for each LFP–LFP coupling measure. This procedure yielded a null distribution for each band and each coupling measure. The observed correlation for each band and coupling measure was considered statistically significant if it exceeded the 95^th^ percentile of its respective null distribution.

For tetrode pairs showing significant correlations in each frequency band, we collected the lags at which their maximum correlations occurred and tested whether these lags differed significantly from zero within each frequency band (Wilcoxon signed-rank test) and whether lags differed between frequency bands or between LFP–LFP coupling measures (i.e., coherence vs. Granger causality [see below]; Wilcoxon rank-sum test).

*Statistical association between ongoing changes in landmark-control strength and changes in LFP coherence*

To further test the statistical significance of positive associations between ongoing changes in landmark-control strength and changes in LFP–LFP coherence while accounting for non-independence among observations from each animal, we performed linear mixed-effects modeling. We fit separate models for each frequency band (theta and gamma). Each model used the position-resolved landmark-control strength as the dependent variable (*y*) and the z-imaginary coherence at the maximally correlated frequency within that band as the predictor (*x*).

For each tetrode pair and frequency band, position-resolved coherence values were mean-centered to remove baseline differences across tetrode pairs, ensuring that the analysis captured the average within-tetrode-pair association rather than being dominated by tetrode pairs with higher or lower overall coherence. After mean-centering, data from all tetrode pairs were concatenated to form the predictor (*x*) and dependent variable (*y*) vectors for the model.

Because multiple 29e–MEC tetrode pairs were recorded from the same rat, and coherence (*x*) and landmark-control strength (*y*) values across spatial bins represented repeated measurements from each tetrode pair, observations were not independent. We modeled this nested dependence structure using four random effects: random intercepts and slopes for rats, and random intercepts and slopes for tetrode pairs nested within rats:

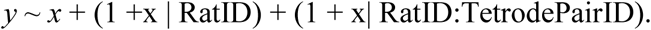

This structure accounts for: (i) rat-level variation in baseline landmark-control strength and in the strength of the *x*–*y* relationship, and (ii) tetrode-pair-level variation in baseline landmark-control strength and in the strength of the *x*–<u>y</u> relationship within each rat. By allowing these parameters to vary across rats and tetrode pairs, the model captures correlations among observations from the same rat and among repeated measurements from the same tetrode pair, providing a conservative test of the population-level fixed effect (i.e., the population-average slope of the association between coherence [*x*] and landmark-control strength [*y*]).

Models were fit using restricted maximum likelihood (using MATLAB’s *fitlme* function). The fixed effect for each model was evaluated using t-test with Satterthwaite-approximated denominator degrees of freedom, as described above. To map the fixed effect strength to a normalized range, the fixed-effect slope was converted to a standardized coefficient (β) by multiplying the raw slope by the ratio of the global standard deviations of *x* and *y*.

### LFP–LFP Granger causality and directed asymmetry index (DAI)

For Granger causality analysis, LFP signals were segmented into candidate windows of 3 seconds with 98% overlap (0.05 s step size). Due to the larger window size used in this analysis compared to LFP– LFP coherence analysis, we adjusted the speed-filtering criterion: velocity was required to exceed 4 cm/s for more than 50% of the window, rather than the 75% threshold used previously. If a candidate window did not satisfy this adjusted criterion, it was discarded.

Within each accepted 3-second window, LFP signals were detrended and transformed to the frequency domain using a multi-taper Fourier transform with Slepian tapers (0.4 Hz resolution) via FieldTrip (*136*). For low frequencies (<20 Hz), the entire 3-second window was used. For higher frequencies (≥20 Hz), each 3-second window was divided into two equal, non-overlapping 1.5-second segments; spectral analysis was performed separately for each segment to improve robustness against nonstationarity of LFP signals in the higher frequencies, and the results were subsequently averaged. The combined spectra from both frequency ranges yielded a full-frequency spectral representation for each window.

Using these full-frequency spectral representations, we computed conditional (partialized) Granger causality (*56*, *151*) for each 29e-MEC tetrode pair and each accepted 3-second window in both directions (i.e., *GC*_29*e*→*MEC*_ and *GC*_*MEC*→29*e*_), conditioning on all other simultaneously recorded tetrodes in 29e, MEC, and other parahippocampal regions via FieldTrip’s *ft_connectivityanalysis* function (configuration: method = ’granger’, conditional = ’yes’). As with our use of imaginary coherency to guard against spurious synchronization from common input, this multi-channel conditioning guards against spurious directionality by removing indirect and common-source effects, to the extent such influences are represented in the conditioning signals. However, whereas imaginary coherency is robust only against instantaneous (zero-lag) common input, conditional Granger causality can mitigate common-source confounds even when shared input reaches 29e and MEC with different delays (*57*, *152–154*). To smooth the resulting Granger causality spectra, a moving average filter spanning 2 frequency bins (i.e., half-width 1 Hz) was applied in the frequency domain.

*Baseline DAI levels*

For each 29e–MEC tetrode pair, the smoothed Granger causality spectra from all accepted windows in the initial stationary landmark epoch (G = 1) were averaged, yielding two mean spectra: one for the 29e→MEC direction (*GC*_29*e*→*MEC*_), and one for the reciprocal MEC→29e direction (*GC*_*MEC*→29*e*_). The directed asymmetry index (DAI; *58*) was then calculated for each pair as the normalized difference between the two:

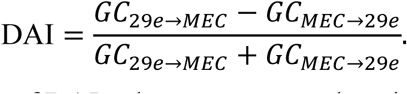

To assess the statistical significance of DAI values across tetrode pairs, we used a cluster-based permutation approach. At each frequency, a one-sample t-test determined whether DAI values across tetrode pairs for a region pair significantly deviated from zero. To control the familywise error rate, null distributions of maximal cluster size were generated by randomly flipping the region labels of tetrode pairs and repeating the t-test procedure for 1000 permutations. Clusters were defined as contiguous frequencies with a significant deviation from zero (*p* < 0.05, two-sided). For each observed cluster, the sum of t-statistics within the cluster was compared to the permutation distribution to generate cluster-level, permutation-corrected *p*-values. Clusters with corrected *p* < 0.05 were considered significant.

*Granger Causality and DAI changes from initial to final epoch*

For each tetrode pair, we calculated the average Granger Causality spectra for both directions (i.e., 29e→MEC and MEC→29e) during the initial (G = 1) and final (G = G_final_) experimental epochs by averaging the Granger causality spectra from all accepted windows in each of these epochs. To extract theta- and gamma-band-specific Granger causality values for each direction and each epoch, we identified the frequency bin with the maximum Granger causality within each band and averaged over ±1 Hz around that peak. For each band and direction, the initial epoch-to-final epoch change in Granger causality was calculated as (GC_final_ – GC_initial_)/(GC_final_ + GC_initial_). To calculate the change in DAI, we first computed theta- and gamma-band DAI in each epoch from corresponding Granger causalities in both directions as above, and the change was calculated as ΔDAI = DAI_final_ – DAI_initial_.

Analysis of ΔDAI values across session types (*LM-control vs. LM-failure)* and across frequency bands (theta vs. gamma) was performed using a linear-mixed effects model with the same random effect structure as in ΔCoherence analysis (see above).

*Temporal relationship between ongoing changes in landmark-control strength and changes in DAI*

Position-resolved, uniformly sampled DAI values were computed using the same resampling procedure described for z-imaginary coherence (see “*Temporal relationship between ongoing changes in landmark-control strength and changes in LFP coherence*”). Briefly, Granger causality spectra from unevenly distributed windows were resampled onto a uniform spatial grid (90° spacing) using local linear kernel regression with a Gaussian kernel (135° standard deviation). DAI was then computed at each spatial bin from the resampled Granger causality spectra in both directions. Both position-resolved DAI and landmark-control strength trajectories were smoothed with a Gaussian filter (240° standard deviation) prior to cross-correlation analysis. Cross-correlation and statistical significance testing followed the same procedures as for z-imaginary coherence.

*Statistical association between ongoing changes in landmark-control strength and changes in LFP coherence*

To test the significance of associations between ongoing changes in landmark-control strength and changes in DAI while accounting for non-independence among observations, we applied linear mixed-effects modeling using the same approach as for z-imaginary coherence (see “*Statistical association between ongoing changes in landmark-control strength and changes in LFP coherence*”). Briefly, for each frequency band (i.e., theta or gamma), position-resolved DAI values were mean-centered within each tetrode pair, and data from all tetrode pairs were concatenated to form the predictor (*x*) and dependent variable (*y*) vectors. The model used the same random-effects structure described above:

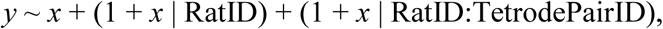

where *x* denotes the position-resolved DAI and *y* denotes the position-resolved landmark-control strength. Models were fit using restricted maximum likelihood (using MATLAB’s *fitlme* function), and fixed effects were evaluated using t-tests with Satterthwaite-approximated denominator degrees of freedom. Fixed-effect slopes were converted to standardized coefficients (β) as described above.

We also assessed the overall relationship between position-resolved trajectories of landmark control-strength score and band-specific LFP–LFP coupling measures across all sessions, animals, and tetrode pairs by fitting linear mixed-effects (LME) models. Separate models were fit for each coupling measure (z-imaginary coherence and DAI) and each frequency band (theta and gamma). In each model, the position-resolved landmark control-strength score served as the predictor variable (fixed effect), and the coupling measure at the frequency yielding the maximal correlation within the corresponding frequency band (theta or gamma) served as the dependent variable. The models included random intercepts and slopes for each animal and random intercepts for each tetrode pair nested within animal to account for the repeated-measures structure of the data. Model parameters were estimated using restricted maximum likelihood (using MATLAB’s *fitlme* function), which also provided statistical significance of the fixed-effect slope based on the t-statistic and associated *p*-value.

### Cross-regional spike–LFP coupling

To quantify cross-regional spike–LFP coupling between 29e and MEC, we computed pairwise phase consistency (PPC) spectra for each neuron relative to LFP channels in the other region. The LFP phase at each frequency was estimated by convolving the continuous LFP trace with complex wavelet-like filters constructed from DPSS tapers using the multitaper convolution method of the FieldTrip toolbox (*132*–*134*, *136*; frequency range: 1–140 Hz; spectral smoothing: 0.25 × center frequency, corresponding to ∼3 tapers; time window: 7 cycles per frequency). For each frequency, the phases at all spike times within the analysis window (e.g., epoch 1) were extracted to form a spike-phase distribution, from which we calculated PPC to obtain an unbiased estimate of spike–LFP coupling strength. We interpolated the obtained PPC spectra to a uniform grid between 1 to 140 Hz with 1 Hz spacing.

*Baseline coupling levels*

We restricted the analysis windows for PPC computation to spikes from the initial epoch with stationary landmarks (i.e., epoch 1, G = 1) and included only neurons with at least 40 spikes in that epoch. Because neurons may show large variability in their coupling to LFP across laminar positions in the other region, we constructed a representative spectrum for each neuron by taking, at each frequency, the maximum PPC across all available LFP channels, capturing the strongest cross-regional coupling for between-area comparisons.

*Spike–LFP coupling change from initial to final epoch*

To examine how cross-regional spike–LFP coupling changed over the course of a session, we compared PPC values between the initial epoch (G = 1) and the final epoch (G = G_final_), calculated for each neuron in each region relative to the other region’s LFP (29e relative to MEC and MEC relative to 29e). Only neurons that fired at least 40 spikes in each of these epochs were included.

When multiple LFP channels were available, we used, for each neuron, its median PPC across channels (instead of the maximum used for between-area comparisons in the baseline) to prevent within-neuron initial-to-final epoch comparisons from being confounded by shifts in the optimal channel. From the median PPC spectra for each neuron in each epoch, we extracted the maximum PPC value within the theta and gamma bands, and quantified the initial-to-final change for each band as (PPC_epoch3_ − PPC_epoch1_) / (PPC_epoch3_ + PPC_epoch1_).

*Temporal relationship between ongoing changes in spike–LFP and landmark-control strength*

To examine the temporal relationship between ongoing changes in cross-regional spike–LFP coupling and landmark control strength, we performed a cross-correlation analysis. Only neurons with at least 8 spikes per lap in at least 75% of laps were included.

For each neuron, we computed the PPC spectrum relative to the LFP from the other region separately for each lap (360° segment) in the reference frame of the rotating landmarks. When multiple LFP channels from the other region were available, we computed the median PPC spectrum across channels for each lap. Both the lap-wise PPC spectra and landmark control-strength scores were smoothed with a Gaussian filter (320° standard deviation) and resampled onto a common grid of 90° spatial bins in the reference frame of rotating landmarks.

We then performed a cross-correlation analysis between the position-resolved trajectories of landmark control strength and PPC at each frequency, computing the Pearson correlation coefficient across spatial lags ranging from −18 to +18 bins (±4.5 laps) and identifying the lag that maximized this correlation.

The maximal correlation coefficient, its corresponding lag and frequency within the theta and gamma bands, and the corresponding PPC values at those frequencies were recorded for each neuron. (These theta and gamma band PPC trajectories were also used in the subsequent section.)

Statistical significance of each neuron’s maximal correlation coefficient within each frequency band (i.e., theta or gamma) was assessed by comparison with a null distribution. The null was generated by circularly shifting the landmark control-strength trajectory by offsets ranging from 32 bins (8 laps) to N−32 bins, and repeating the identical cross-correlation procedure for each shift—i.e., computing correlations across all frequencies and lags, and recording the maximum within each frequency band. A neuron’s observed correlation in a frequency band was considered statistically significant if it exceeded the 95th percentile of its respective null distribution.

For neurons showing significant correlations in each region, we collected the lags at which their maximum correlations occurred. We then tested whether these lags differed significantly from zero within each region (Wilcoxon sign-rank test) and whether lags differed between 29e and MEC (Wilcoxon rank-sum test).

*Statistical association between ongoing changes in landmark-control strength and changes in spike– LFP coupling*

To test the significance of associations between ongoing changes in landmark-control strength and changes in spike–LFP coupling while accounting for non-independence among observations, we applied linear mixed-effects modeling using a similar approach as for LFP–LFP coherence (see “*Statistical association between ongoing changes in landmark-control strength and changes in LFP coherence*”).

Separate models were fitted for neurons in each region (29e neurons to MEC gamma and MEC neurons to 29e theta), yielding two models in total.

For each neuron, we used the position-resolved, cross-regional PPC values identified in the previous section, corresponding to the theta- and gamma-band frequencies with the maximum correlation coefficient. These PPC values were mean-centered within each neuron to remove baseline differences, ensuring that the analysis captured within-neuron variation rather than being dominated by neurons with higher or lower overall coupling strength. After mean-centering, data from all neurons within a region were concatenated to form the predictor (*x*: landmark-control strength) and dependent variable (*y*: PPC) vectors for the model of each regions’ neurons.

Because multiple neurons were recorded from the same rat, and values of position-resolved PPC and landmark-control strength represented repeated measurements across spatial bins from all neurons recorded in a session, we used random effects (both intercepts and slopes) at both the rat and neuron levels. In mixed-effects notation, each model took the form:

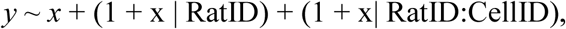

This random-effects structure allowed the relationship between landmark-control strength (*x*) and PPC (*y*) to vary across rats and neurons, capturing correlations among observations from the same rat and among repeated measurements from the same neuron, and thereby providing a conservative test of the population-level fixed effect. As in LFP–LFP analyses, model parameters were estimated using restricted maximum likelihood (using MATLAB’s *fitlme* function), and the fixed effect (i.e., the population-average slope) was evaluated using t-tests with Satterthwaite-approximated denominator degrees of freedom.

### Identification of monosynaptic excitatory connections

We ran the GLMCC algorithm (*155*) in likelihood-ratio mode (*156*), which returned a directed connectivity matrix among all simultaneously recorded neurons in each session. A neuron was classified as excitatory if ≥ 80% of its outgoing weights were positive. Monosynaptic connections of excitatory neurons were then defined as outgoing edges with weights exceeding a threshold of 1, as suggested in (*155*).

## Supplementary Figures

**Fig. S1.**
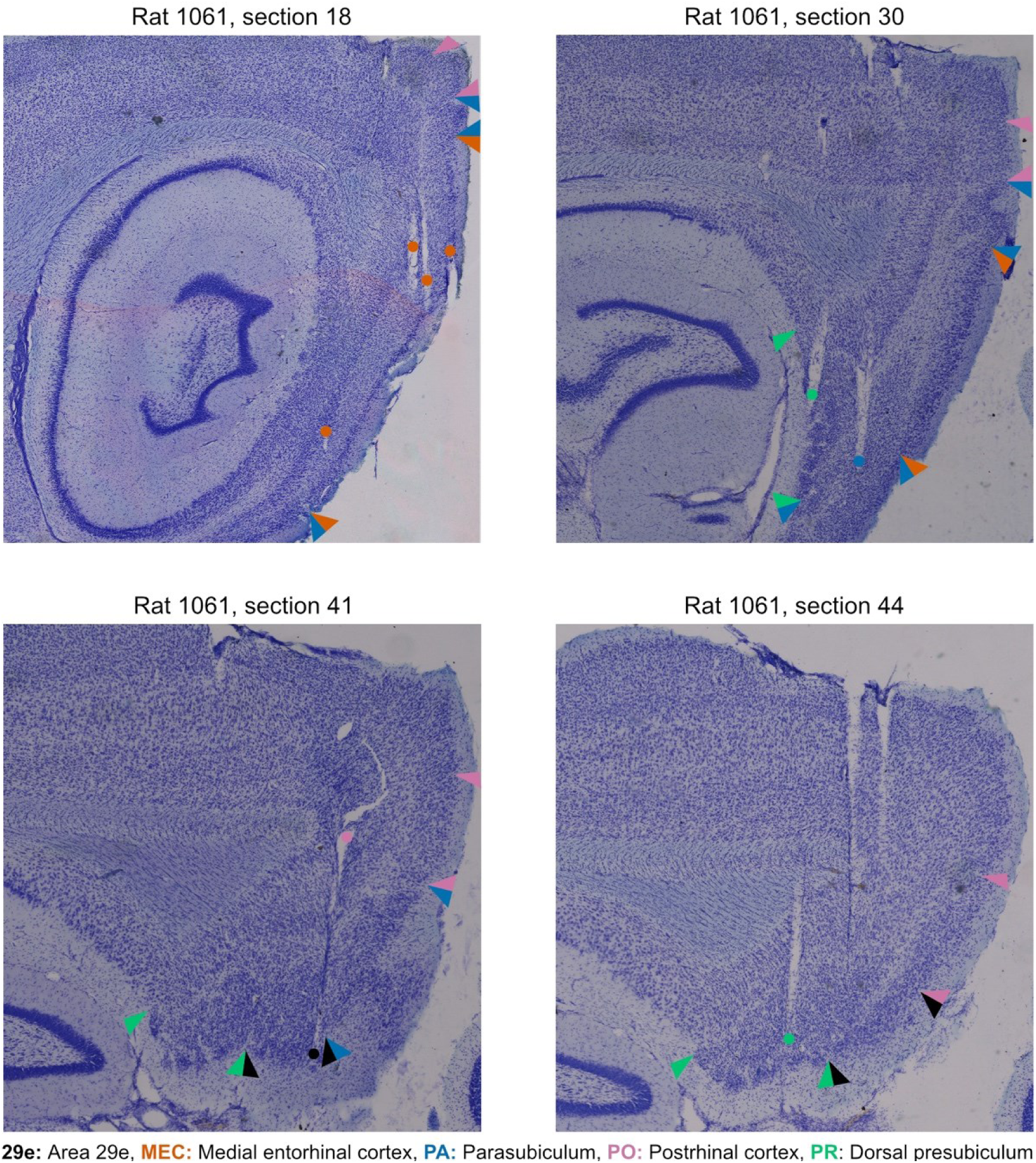
Representative Nissl-stained sagittal sections showing tetrode recording sites across parahippocampal regions in Rat 1061. Four sections are arranged from lateral (top left) to medial (bottom right). As in Fig. 1, colored arrowheads indicate the superficial borders of recorded parahippocampal regions namely, area 29e (black), MEC (orange), PA (blue), PO (pink), and PR (green). Dots mark tetrode recording locations, with dot color corresponding to the recorded region.

**Fig. S2.**
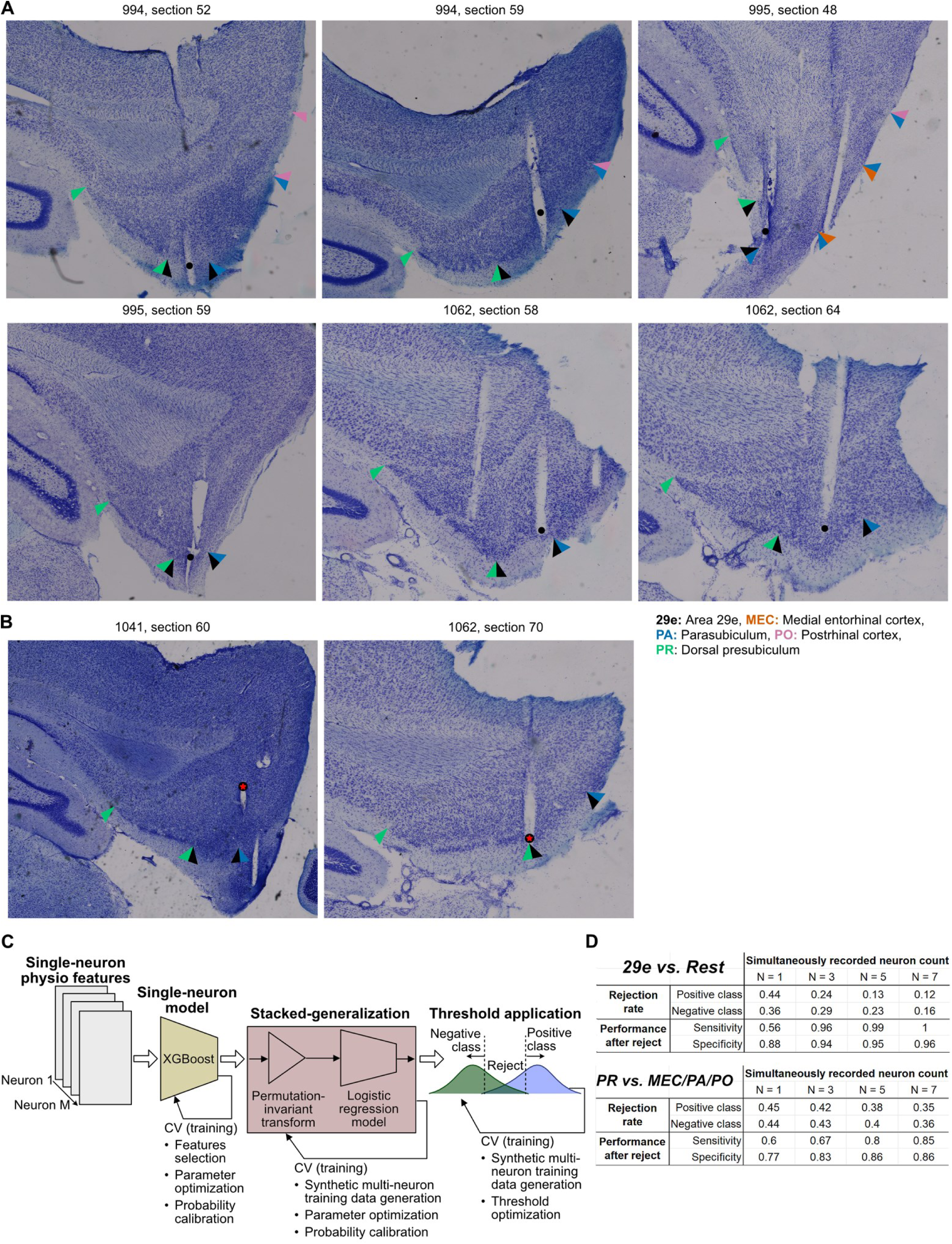
Histologically identified recording sites in area 29e and machine-learning pipeline using single-neuron physiological properties applied to resolve ambiguous histological cases (see Methods). (**A**) Example histological sections illustrating recording sites in area 29e from three rats (994, 995, and 1062). See fig. S1 for an additional histologically identified 29e site (rat 1061). Colored arrowheads indicate boundaries between areas following the same color scheme as Fig. 1A and fig. S1: area 29e (black), MEC (orange), PA (blue), PO (pink) and PR (green). (**B**) Example histological sections illustrating recording sites with ambiguous localization between area 29e and neighboring regions. Left: Rat 1041, section 60 shows a recording location at the 29e/PA border (indicated by the red star). Application of the machine learning model (see panels C, D and the methods) classified this location as area 29e. Right: Rat 1062, section 70 shows a recording location at the 29e/PR border (red star) that was likewise classified as area 29e by the machine learning model. (**C**) Machine-learning model architecture. When histological identification of a tetrode’s recording region was ambiguous (as in panel B), we employed a machine-learning classification approach using physiological features of single neurons recorded on the tetrode to identify the region the tetrode was located in. Because ambiguous tetrode tracks were most often along the borders of area 29e or PR with neighboring regions, this approach relied on two separate one-vs.-rest classification pipelines (29e vs. rest and PR vs. rest) to distinguish recording sites in 29e from other regions and those in PR from other regions, respectively. Each pipeline was trained exclusively on neurons from tetrodes with unambiguous histological region assignments (ground-truth data) and proceeded through three steps: *Step 1 (Single-neuron classifier):* An XGBoost classifier used each neuron’s physiological features to estimate the probability that it belonged to the target region. Cross-validation (CV) on the ground-truth training data was used for feature selection, hyperparameter optimization, and probability calibration. *Step 2 (Stacked-generalization as multi-neuron evidence accumulator)*: Probability outputs from the single-neuron classifiers were aggregated across all simultaneously recorded neurons on the same tetrode into a single tetrode-level probability estimate by applying a permutation-invariant transformation followed by a logistic-regression “meta-model” used for stacked-generalization. For hyperparameter optimization and probability calibration of this meta model, CV was performed on synthetic multi-neuron samples generated by combining and subsampling ground-truth neurons. *Step 3 (Reject-option thresholding):* Class-specific confidence thresholds were applied to the tetrode-level probabilities to assign either the positive (e.g., 29e) or negative (e.g., not 29e) label only when the corresponding probability exceeded its threshold; otherwise, we made no assignment (reject). Threshold values were optimized via CV on the synthetic multi-neuron samples (as in step 2) to balance coverage and accuracy. See Methods for further details. (**D**) Machine-learning model validation and application criteria. To validate model performance, we performed tetrode-stratified fivefold cross-validation (CV), ensuring neurons from the same tetrode never appeared in both training and test sets. The table reports performance metrics evaluated on held-out test data as a function of the number of simultaneously recorded neurons per tetrode (N = 1, 3, 5, or 7). Because individual held-out tetrodes had variable neuron counts, synthetic test cases were generated by pooling neurons across held-out tetrodes within each class and randomly sampling N neurons to simulate tetrodes with a specific neuron count. For each pipeline (29e vs. rest, top; PR vs. rest, bottom), metrics include: (i) the rejection rate: the fraction of test cases where the pipeline declined to assign a label because neither the positive-class nor negative-class probability exceeded its respective confidence threshold. Rows show separate rejection rates for tetrodes truly belonging to the positive class (e.g., 29e/PR) versus the negative class (e.g., not 29e/not PR). Lower rejection rates indicate the pipeline more frequently made confident assignments. (ii) performance after reject: Among the test cases that were not rejected (i.e., received a confident assignment), sensitivity is the proportion of true positive-class tetrodes correctly identified as positive, and specificity is the proportion of true negative-class tetrodes correctly identified as negative. As the number of simultaneously recorded cells on a tetrode increased, rejection rate decreased, and sensitivity/specificity increased for both models. To ensure reliable assignments (both sensitivity and specificity ≥ 0.85), we applied the 29e vs. rest pipeline only to tetrodes with at least 5 simultaneously recorded neurons and the PR vs. rest pipeline only to tetrodes with at least 7 simultaneously recorded neurons. Final models trained on all ground-truth data (i.e., data from tetrode recording locations with histologically unambiguous region assignments) were then applied to classify histologically ambiguous tetrode recordings meeting these criteria (see Methods).

**Fig. S3.**
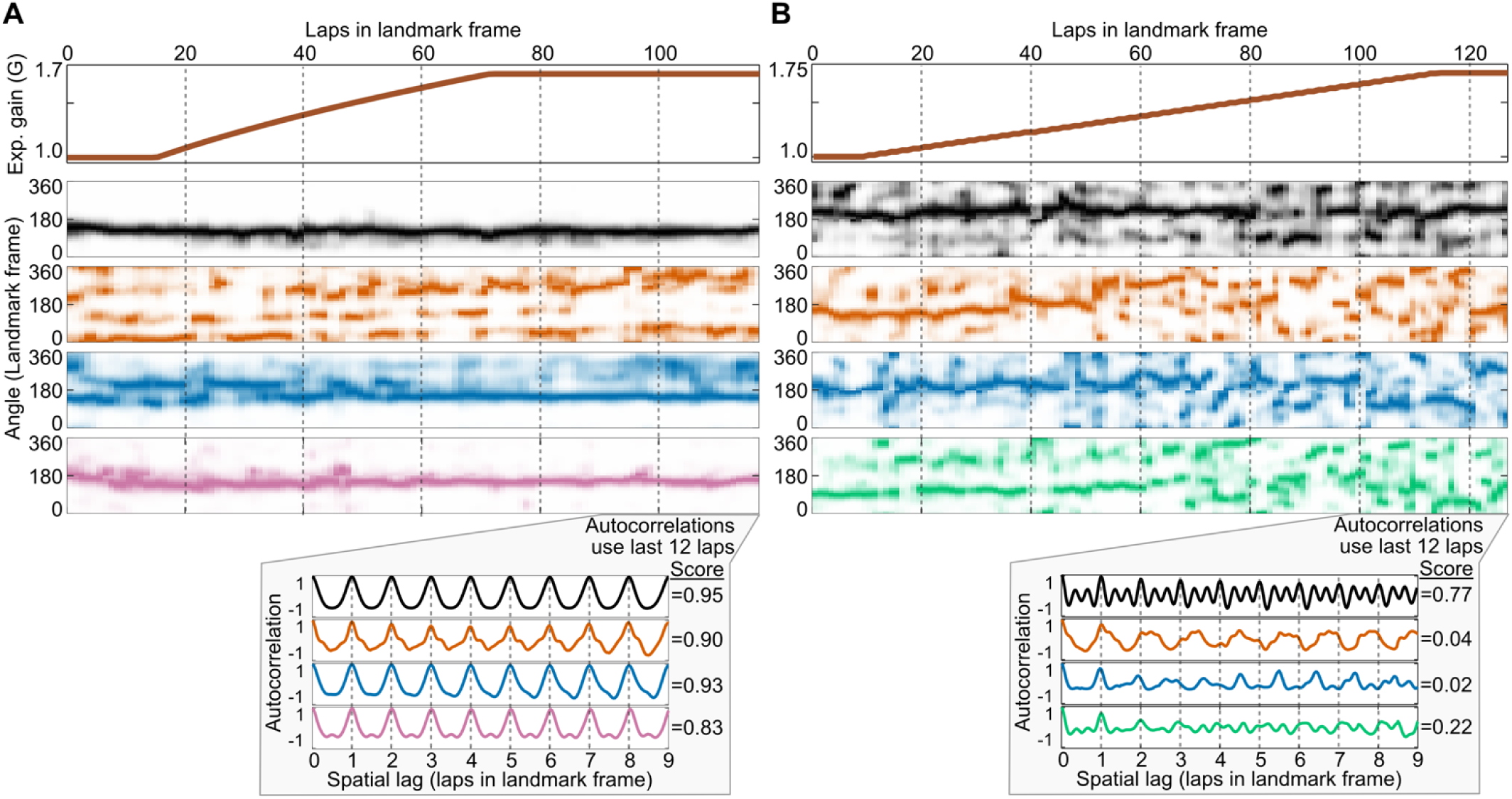
Examples of landmark anchoring in 29e relative to other regions. Figure organization and panels are similar to Fig. 1E-F. **(A)** Example *LM-control* session in which spatially correlated neurons remain anchored to the rotating landmarks. Top panel shows the experimental gain protocol (gain = 1 at baseline, then ramped to a higher target gain and held there for 10-30 laps). Middle panel shows the spatial firing rates of simultaneously recorded neurons across laps in the landmark reference frame. Neurons fired at fixed angles relative to landmarks throughout the session, producing stationary, horizontal bands of firing rates in the landmark frame. Bottom shows the spatial autocorrelograms of the last 12 laps for example neurons. Clear peaks indicate strong 1-lap periodicity; the accompanying score quantifies this periodicity (see “landmark-locking score” in Methods). **(B)** Example *LM-failur*e session in which most spatially correlated neurons from regions other than 29e lose landmark anchoring. Conventions as in (A). Around laps 65-75, firing fields of neurons recorded outside 29e started to drift in the landmark frame, yielding autocorrelograms that were not lap-to-lap periodic and that exhibited low landmark-locking scores. By contrast, a subset of 29e neurons (like the example in black) remained locked to landmarks, firing stably at fixed angular locations in the landmark frame. In this example, the 29e cell had 2 firing fields: one at ∼90^°^ and one at ∼180^°^. Early in the session, the field at ∼90^°^ fired variably while the field at ∼180^°^ was more consistent. Shortly after the landmark-control failure outside 29e, the field at ∼90^°^ strengthened, while the field at ∼180° weakened. This reversed pattern persisted for about 10 laps, until lap 90. After this point, fields returned to their original relative firing levels, with the field at ∼180^°^ firing more consistently than the field at ∼90^°^. Despite these fluctuations in relative field strength, the locations of fields remained anchored to landmarks throughout. This landmark-anchoring of 29e neurons was reflected in their spatial autocorrelograms as a clear 1-lap periodic pattern (black trace in the bottom inset).

**Fig. S4.**
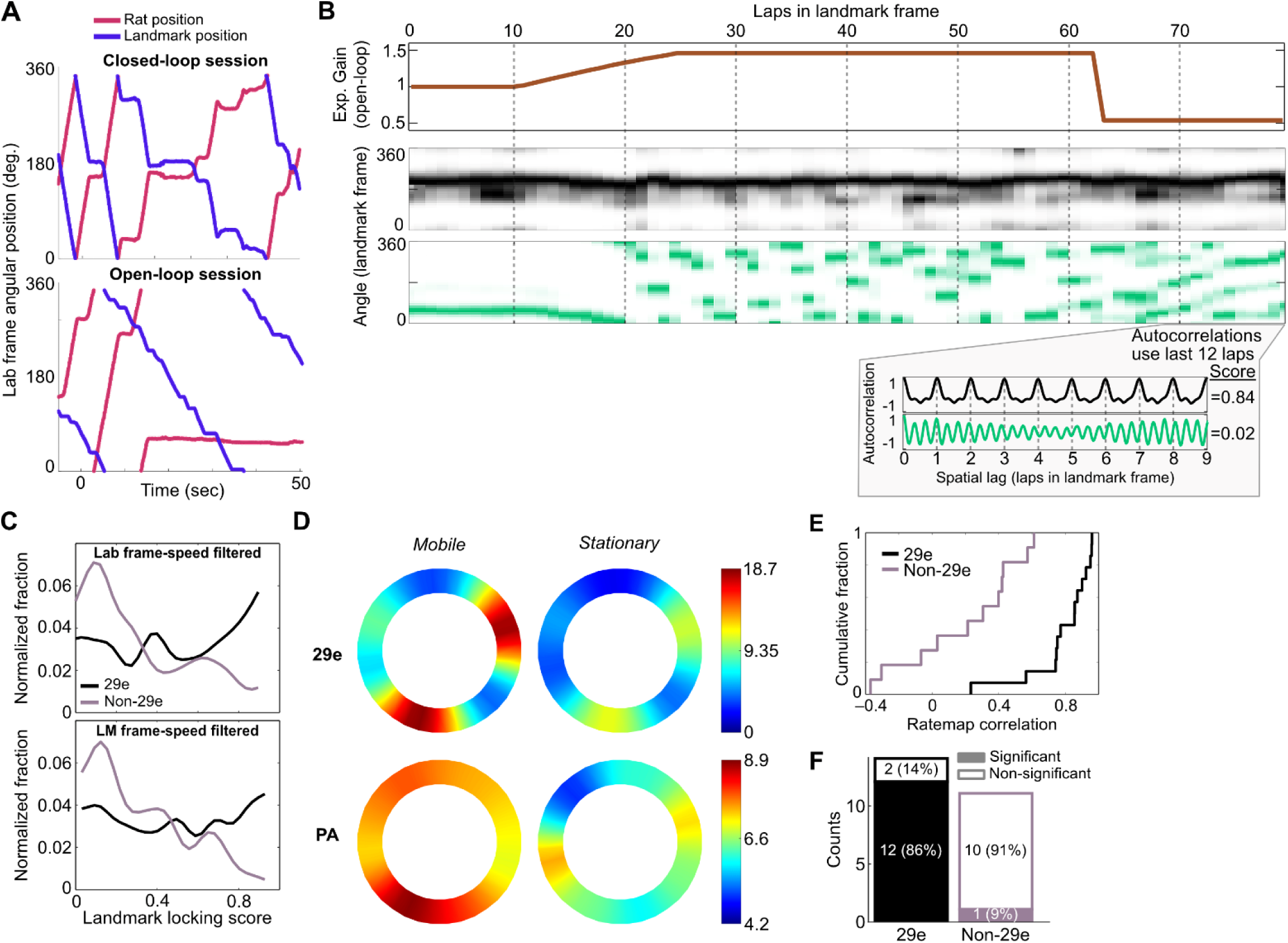
Greater landmark anchoring in neurons of 29e compared to other regions during open-loop manipulation of visual landmarks. **(A)** Example rat and landmark trajectories in closed- and open-loop experiments. Top: Closed-loop (G > 1) session in which landmark rotation was coupled to the rat’s movement. Bottom: Open-loop session in which landmark rotation was yoked to a rat’s trajectory from a different session; note multi-second bout of immobility (starting at ∼15 s) where the rat was stationary while landmarks rotated past it. **(B)** Landmark-frame spatial firing rate map plot across laps in the landmark frame of reference (format same as in Fig. 1E,F) for an example 29e neuron (middle; black) and a PR neuron (bottom; green). The open-loop experiment gain is shown on top. The example PR neuron loses landmark anchoring towards the end of the gain ramp epoch, whereas the 29e neuron remained anchored to landmarks throughout. Notably, the 29e neuron maintains a lap-to-lap periodicity of 1 even when landmarks are moved in open-loop fashion, as evident in the autocorrelation trace, in contrast to the PR neuron, which is no longer controlled by landmarks. **(C)** Distributions of landmark-locking scores in open-loop experiments for 29e (n = 29 neurons) versus non-29e (n = 39 neurons) regions. Neurons outside area 29e were pooled into a single non-29e group due to lower sample sizes with unambiguous histological assignments and similar closed-loop response profiles across regions. Scores were computed with velocity filtering in the laboratory frame (top) and in the landmark frame (bottom). 29e neurons showed higher landmark-locking scores in both cases (Lab frame: median [IQR] 29e = 0.431 [0.193, 0.741], Non-29e = 0.231 [0.107, 0.536], Wilcoxon rank-sum test: *Z =* 2.28, *p* = 0.023; Landmark frame: median [IQR] 29e = 0.484 [0.231, 0.725], Non-29e = 0.275 [0.139, 0.474], Wilcoxon rank-sum test: *Z =* 2.08, *p* = 0.037). **(D)** Comparison of circular spatial firing rate maps across track angles in the landmark reference frame. Representative examples are shown for a landmark-anchored 29e neuron (top) and a landmark-anchored PA neuron (bottom) under open-loop conditions when the rat was *mobile* (left) and *stationary* (right; see Methods for details). Note that the spatial firing rate map for the 29e neuron shows greater similarity between the *mobile* and *stationary* conditions compared to the PA neuron. **(E)** Cumulative distributions of landmark frame rate-map correlations between stationary and mobile periods (*stationary*: the animal’s speed in laboratory frame < 4 cm/s, *mobile*: otherwise; see Methods) for landmark-anchored 29e (n = 14) and non-29e (n = 11) neurons. 29e landmark-anchored neurons showed higher stationary-vs-moving rate map correlations than non-29e landmark-anchored neurons (median [IQR] 29e = 0.856 [0.746, 0.924], Non-29e = 0.306 [-0.042, 0.425], Wilcoxon rank-sum test: *Z =* 3.75, *p* < 0.001). **(F)** Proportion of landmark-anchored neurons with significant stationary-vs.-moving rate-map similarity based on a shuffle test (10,000 circular-shifted shuffles of the stationary map relative to the moving map; α=0.05). Most 29e neurons remain significantly landmark-anchored when the animal was stationary, unlike non-29e neurons (Fisher’s exact test: *p* < 0.001).

**Fig. S5.**
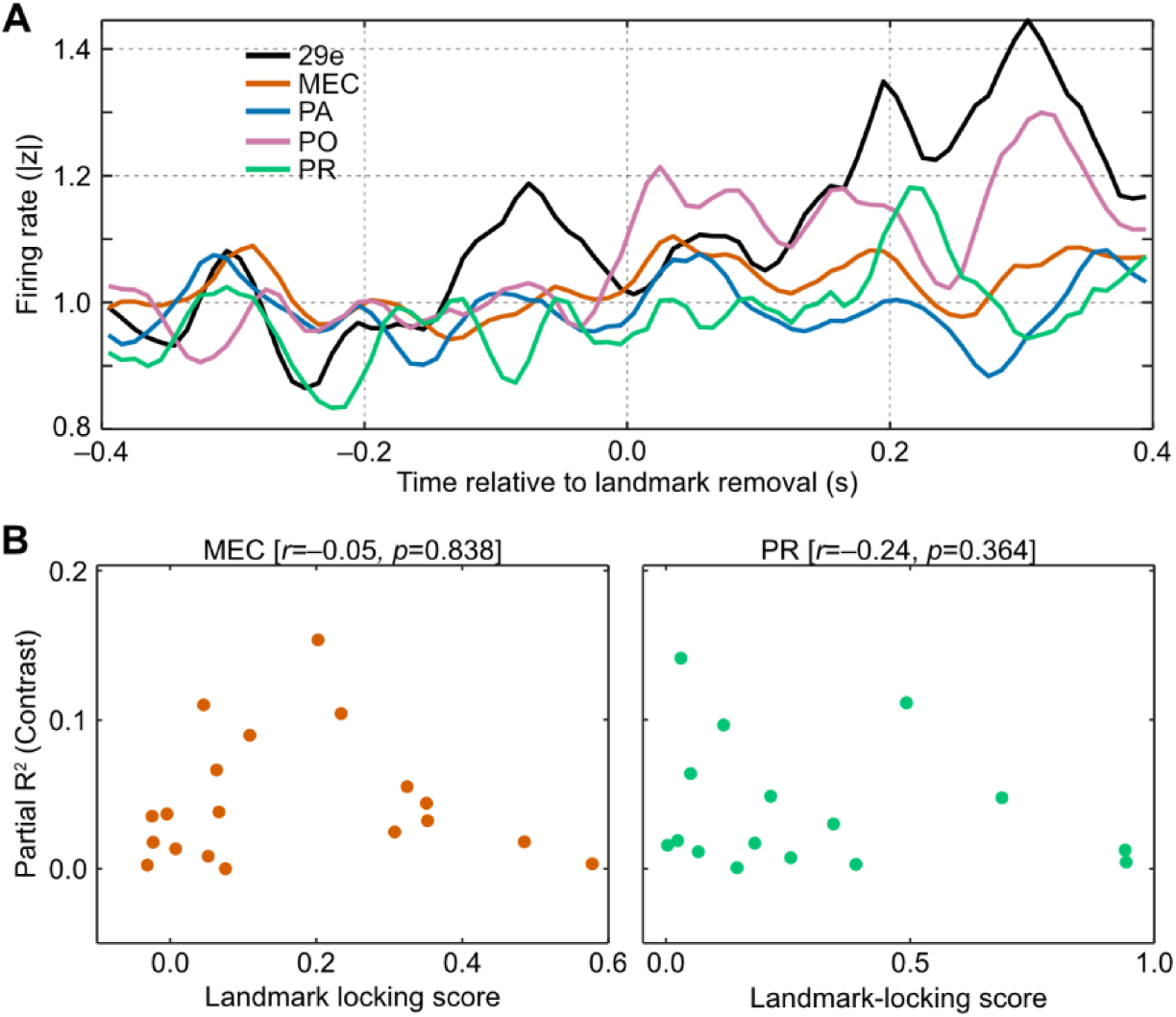
Regional firing-rate responses to landmark removal and relationship between contrast tuning and landmark locking. (**A**) Average baseline-normalized firing-rate deviation around landmark removal (time 0) across all tetrodes in each region. Because landmarks were removed only once at a single timepoint in each session, most neurons with spatially localized firing fields would not have been active at that moment (i.e., rat’s location at the time of removal would not necessarily be within the spatial receptive field boundaries), making landmark removal effects difficult to quantify reliably at the single-neuron level. We therefore used tetrode-level multi-unit activity. For each tetrode, we computed an unsmoothed peri-event firing-rate trace, standardized it using that tetrode’s own pre-removal baseline mean and variability (z-scoring relative to the pre-removal window), and took the absolute value to capture upward and downward deviations from the baseline mean symmetrically. The resulting rectified z-scored trace of each tetrode was then divided by its mean value during the pre-removal, baseline period. and traces were averaged across tetrodes within each region to compute the regional mean. Although the rescaled trace may show large fluctuations around 1 for a single tetrode, the regional means are expected to fluctuate closer to 1 during the pre-removal baseline period as a result of averaging across many tetrodes, as seen in the figure for all regions when t < 0. For the post-removal period, however, firing rates of tetrodes in 29e and PO consistently showed large deviations (|z| > 1) from their average baseline levels,, indicating a region-wide response to the removal of landmarks. (**B**) Absence of significant correlation between landmark-locking scores and contrast tuning strength in MEC and PR neurons. Each dot represents a single neuron’s landmark-locking score (*x*-axis) versus its partial R² for landmark contrast tuning (*y*-axis) in MEC (left) and PR (right). Pearson’s correlation coefficients (*r*) and corresponding *p*-values (computed using Fisher z-transformed landmark-locking scores) are shown above each plot. An ANCOVA with landmark-locking score as a continuous covariate and region as a categorical factor revealed a significant positive relationship between landmark-locking and contrast tuning across regions (F(1, 67) = 5.68, p = 0.020). There was also a significant main effect of region (F(2, 67) = 3.71, p = 0.030), likely reflecting higher contrast tuning in 29e (Fig. 2), but no significant interaction (F(2, 67) = 1.92, p = 0.155).

**Fig. S6.**
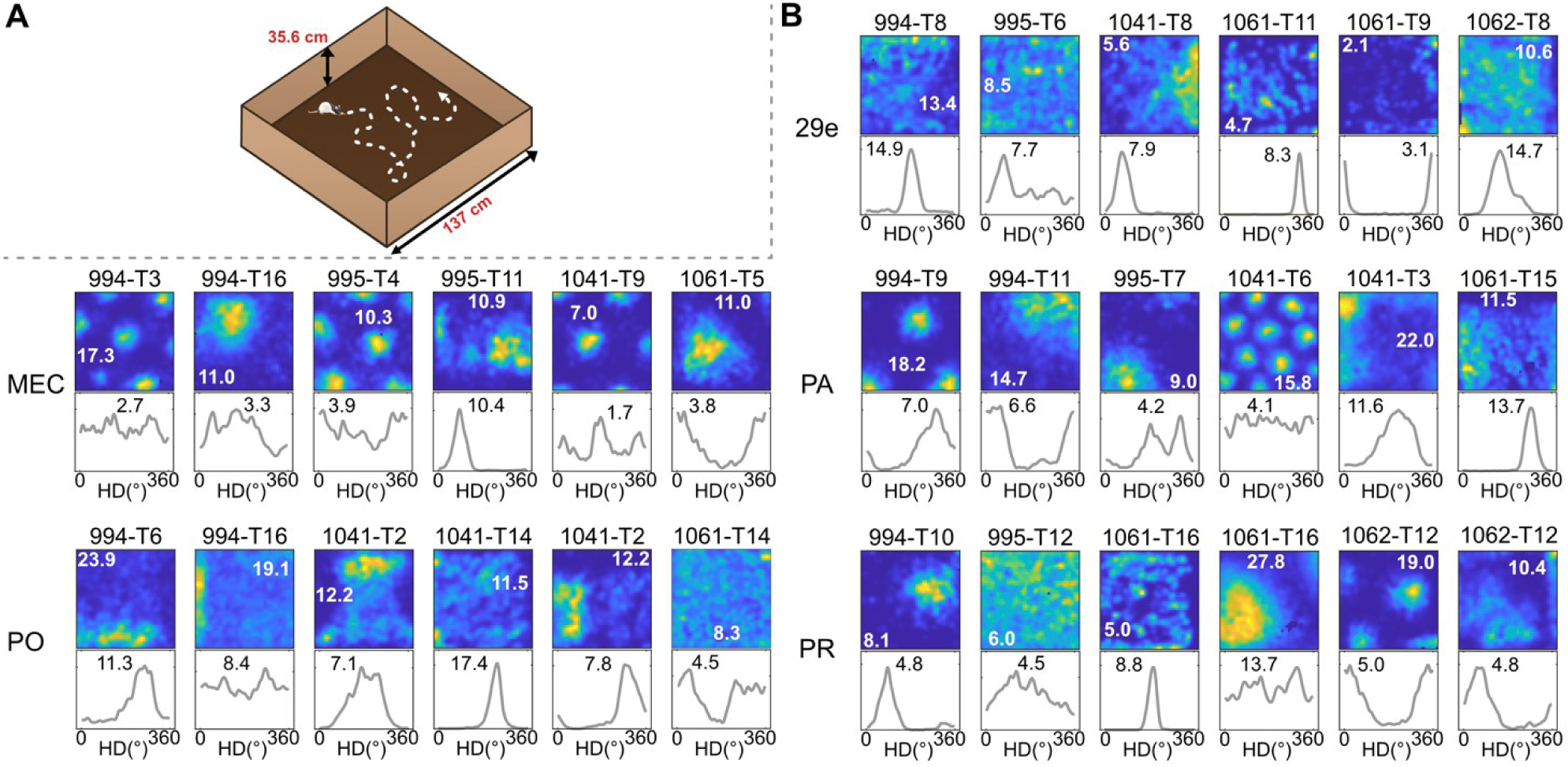
Two-dimensional allocentric spatial tuning in open arena across parahippocampal regions. **(A)** Schematic of the square open arena apparatus used to identify functional parahippocampal cell types. The open arena was placed on the floor in a room with multiple distal visual cues (not shown), providing stable landmarks. **(B)** Tuning to allocentric position and head direction across regions. Five representative neurons are displayed per region. Same format as Fig. 3A.

**Fig. S7.**
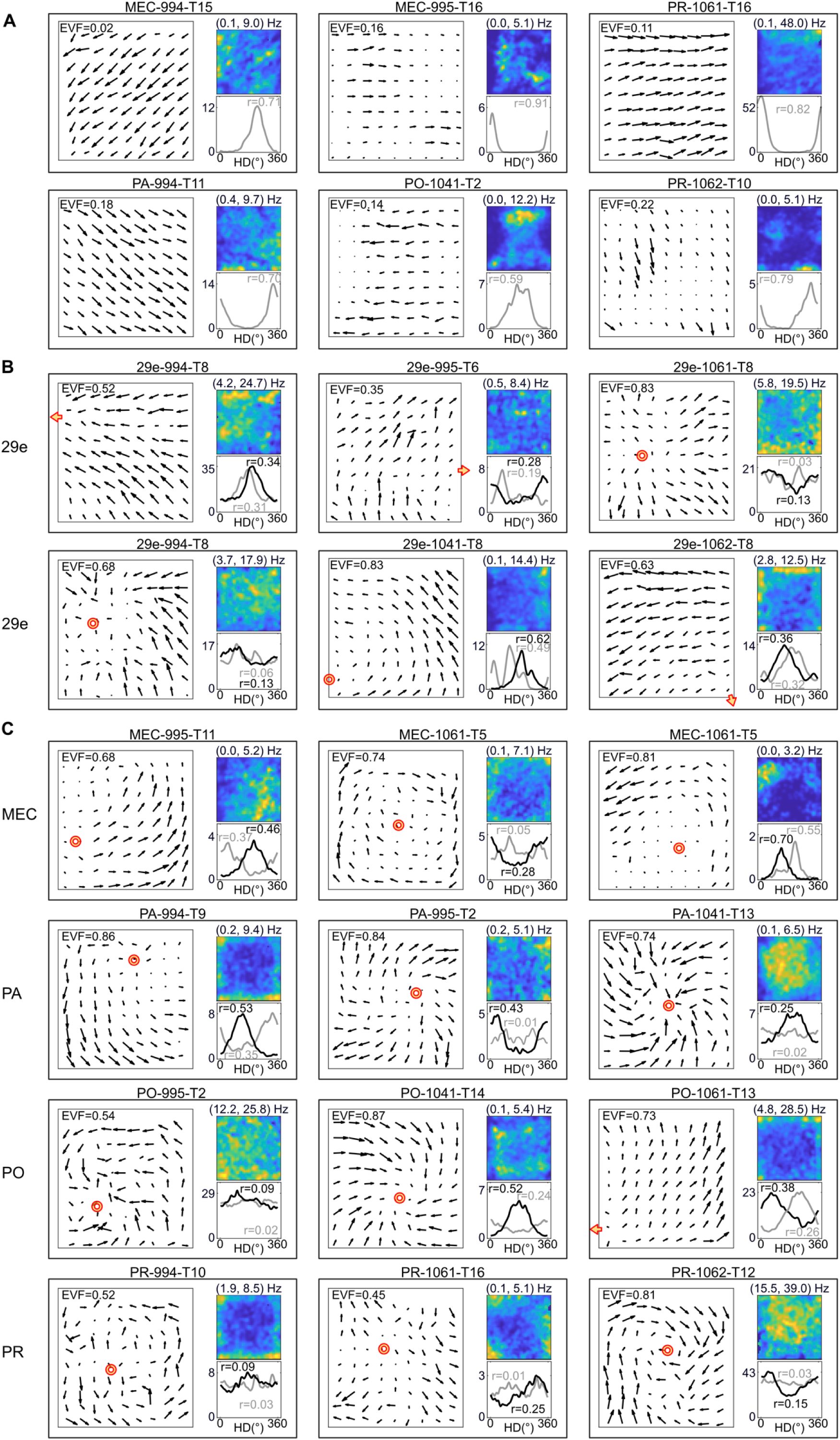
Egocentric tuning of example neurons across different regions. Same format as Fig. 3C. **(A)** Example HD neurons that show classic, allocentric tuning. The preferred HD vectors are roughly parallel at all locations across the open arena, as expected from their allocentric nature. Some cells showed a modulation by both place and HD. Note that it is impossible (and meaningless) to distinguish an allocentric HD cell from an egocentric HD cell with an anchor point infinitely far away. Rat ID and tetrode number for each neuron are indicated along with the region identity above the respective panels. **(B)** Example egocentric HD neurons from 29E. **(C)** Example egocentric HD neurons from MEC, PA, PO, and PR.

**Fig. S8.**
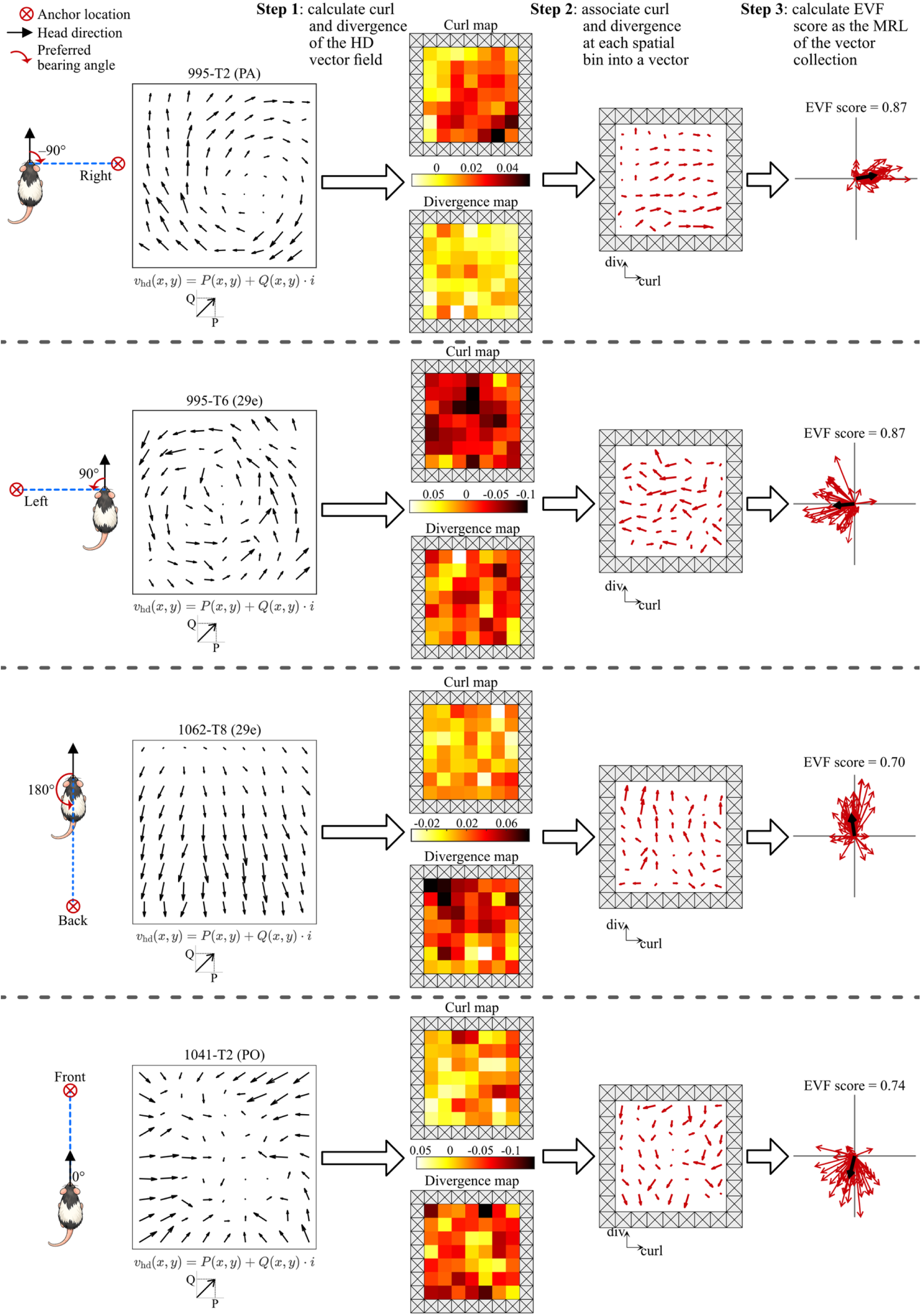
Workflow for computing the anchor-free egocentric-vector-field (EVF) score, illustrated with four example head-direction (HD) cells. A key question for HD-tuned neurons is whether their preferred direction is allocentric or egocentric. Standard approaches to quantifying egocentric tuning rely on an exhaustive search for an optimal anchor point in the environment that serves as a reference point to calculate the bearing angle between the rat’s current allocentric HD at a particular location in the environment and an anchor point (as illustrated by the rat cartoons on the left). The sharpest tuning curve among multiple candidates (each corresponding to a different anchor point) is considered the “true” egocentric tuning curve, against which a classic allocentric tuning curve is compared. This procedure creates a statistical bias in favor of egocentric tuning (the best curve selected from multiple candidates) over allocentric tuning (a single curve). To enable unbiased comparison, we developed an anchor-free Egocentric Vector Field (EVF) score that quantifies the spatial consistency of local head-direction preferences across the arena without specifying any anchor point. See Methods for further details. Figure organization: Each row corresponds to one example neuron. The leftmost column is for illustration purposes and approximately depicts the neuron’s preferred egocentric bearing, defined as the angle (red arc) between the allocentric HD vector (black arrow) located at the rat’s head and the head-to-anchor point vector (blue dashed line). The remaining columns illustrate the three-step workflow for computing the EVF score to quantify such egocentric tuning of single neurons. **Input**: HD vector field. The HD tuning curve within each spatial bin (*x, y*) is reduced to a single vector corresponding to the first Fourier component of the circular tuning curve (i.e., the single sinusoid that, in a least-squares sense, best fits the full circular tuning curve). The phase of this Fourier component is mathematically equivalent to the preferred HD (i.e., the mean angular direction) and its amplitude equals the summed firing rate of the HD tuning curve × the mean resultant length of the HD tuning curve. Collectively, these HD tuning vectors define a two-dimensional vector field *v*_*hd*_(*x*, *y*) = [*P*(*x*, *y*); *Q*(*x*, *y*)], where *P* and *Q* are the horizontal and vertical components of each vector, respectively. **Step 1**: Curl- and divergence-maps. For every spatial bin, we compute a curl score as 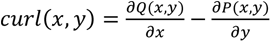and a divergence score as 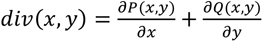. The resulting scalar values are displayed as heat maps (curl on top, divergence on bottom), where lighter colors correspond to values close to 0 and darker colors correspond to progressively larger absolute values. Therefore, the darkest color can represent either a large positive (e.g., CW, centrifugal) or a large negative (e.g., CCW, centripetal) value, depending on the neuron. The color scale is normalized separately for each neuron**. Step 2**: Curl/divergence vectors. For each spatial bin, the ordered pair (curl, div) is treated as a new 2D vector (horizontal = curl, vertical = div), forming the curl-divergence field shown next. Curl captures rotational patterns (preferred egocentric bearing to the left or right; top two rows). Divergence captures radial patterns (preferred egocentric bearing to the front or back; bottom two rows). **Step 3**: Calculation of the EVF score. All (curl, div) vectors are replotted in polar coordinates, with the black thick arrow showing their mean resultant vector. The circular statistics of this (curl, div) vector collection define the preferred bearing angle of an egocentric neuron and its EVF score as follows: • The circular mean indicates the neuron’s preferred egocentric bearing angle underlying the orientation of its preferred HD vectors across the arena as shown in the HD vector field in the input column. The examples from top to bottom illustrate this relationship for bearing angles associated with the four cardinal egocentric directions as depicted by the rat cartoons: rightward-pointing (curl,div) (i.e., pure positive curl) for right-anchoring, leftward-pointing (curl,div) (i.e., pure negative curl) for left-anchoring, upward-pointing (curl,div) (i.e., pure positive divergence) for front-anchoring, and downward-pointing (curl,div) (i.e., pure negative divergence) for back-anchoring. • Measuring the circular dispersion of the vector collection (i.e., its disorganization across the arena) with the mean resultant length gives the anchor-free EVF score (0 = randomly distributed vectors, 1 = perfectly aligned vectors).

**Fig. S9.**
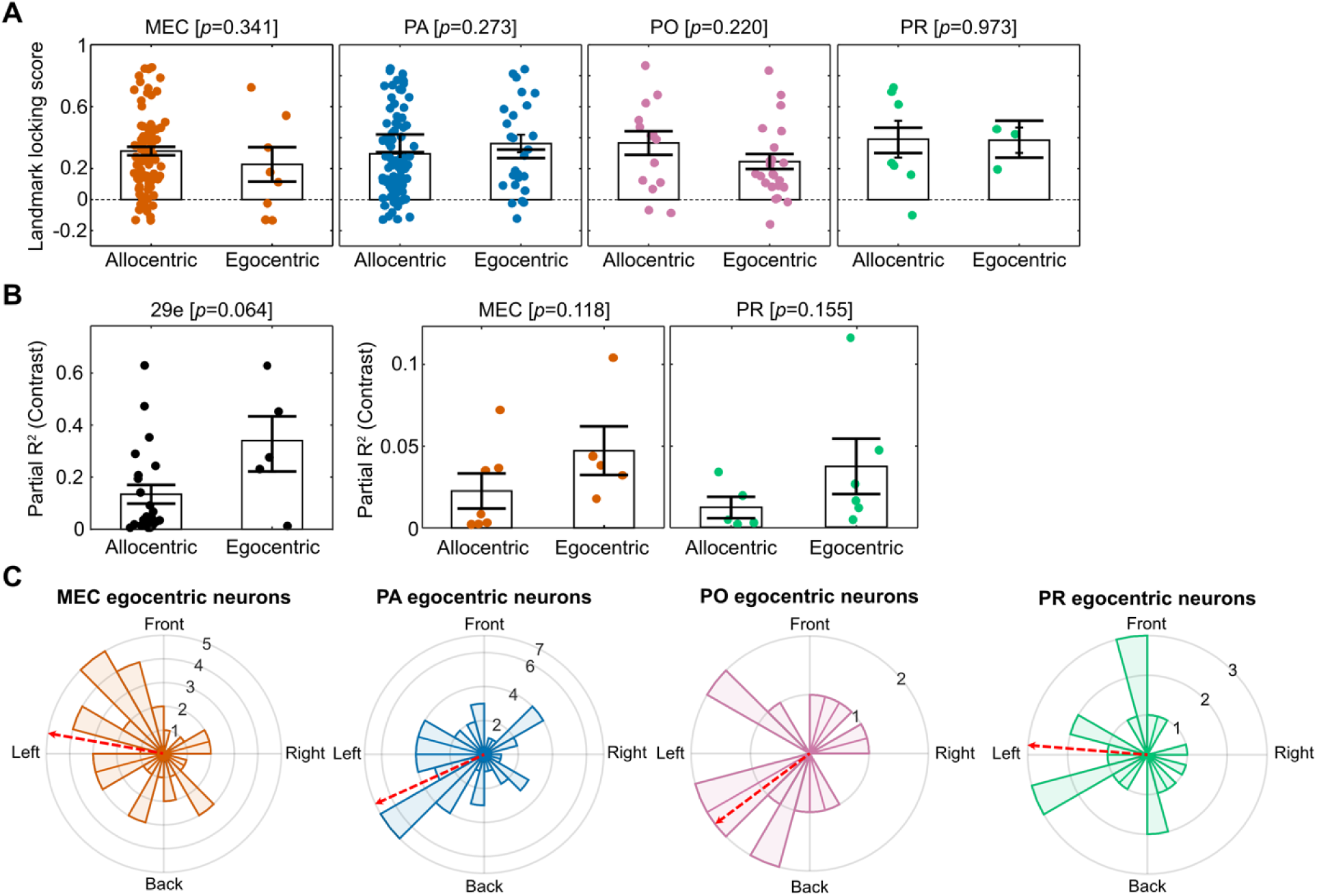
Functional properties of egocentric-versus allocentric-HD tuned neurons across parahippocampal regions. **(A)** Relationship between neurons’ landmark-locking scores (from *LM-failure* sessions) and their preferred frame of reference (egocentric vs. allocentric) in MEC, PA, PO, and PR. Unlike area 29e, where neurons are approximately equally split between allocentric and egocentric reference frames (Fig. 3D) and landmark-locking scores show considerable heterogeneity during LM-failure sessions (Fig. 2H, right), other regions showed imbalanced egocentric-allocentric splits (predominantly allocentric) with relatively homogeneous (and lower) landmark-locking scores. Given the small number of egocentric neurons and limited variability in landmark-locking scores, any observed differences within non-29e regions could reflect session-to-session or animal-to-animal variability rather than true differences in landmark anchoring between egocentric and allocentric neurons. We therefore did not perform a direct statistical comparison between 29e and other regions. Nevertheless, data from other regions are shown here for completeness, with *p*-values from Wilcoxon’s rank-sum tests (egocentric vs. allocentric) displayed above each plot for reference. **(B)** Relationship between neurons’ partial R² values for landmark-contrast tuning (from luminance-manipulation sessions) and their preferred reference frame (egocentric vs. allocentric) in area 29e (left), MEC (middle), and PR (right). As with landmark-locking scores (panel A), contrast tuning was stronger and more heterogeneous in 29e than in other regions (Fig. 2), making this comparison primarily relevant for 29e. For similar reasons as described in panel A, combined with lower neuron counts in luminance-manipulation sessions overall, we did not perform a direct statistical comparison across areas. Instead, we provide within-region comparisons (p-values from two-sample *t*-tests shown above each plot) for reference. Conventions as in panel A. The panels shows that even though no region shows statistical significance, all regions show a trend in the same direction. **(C)** Polar histograms showing the distribution of preferred bearing angles for egocentric HD cells in MEC, PA, PO, and PR. Same format as Fig. 3G. All regions showed a qualitative leftward bias in preferred bearing angles, contralateral to the recording hemisphere. This lateralization was statistically significant in MEC (left: n = 30, right: n = 14; binomial test, *p* = 0.023) and PA (left: n = 37, right: n = 17; *p* = 0.009), but not in PO (left: n = 11, right: n = 7; *p* = 0.481) or PR (left: n = 13, right: n = 8; *p* = 0.383).

**Fig. S10.**
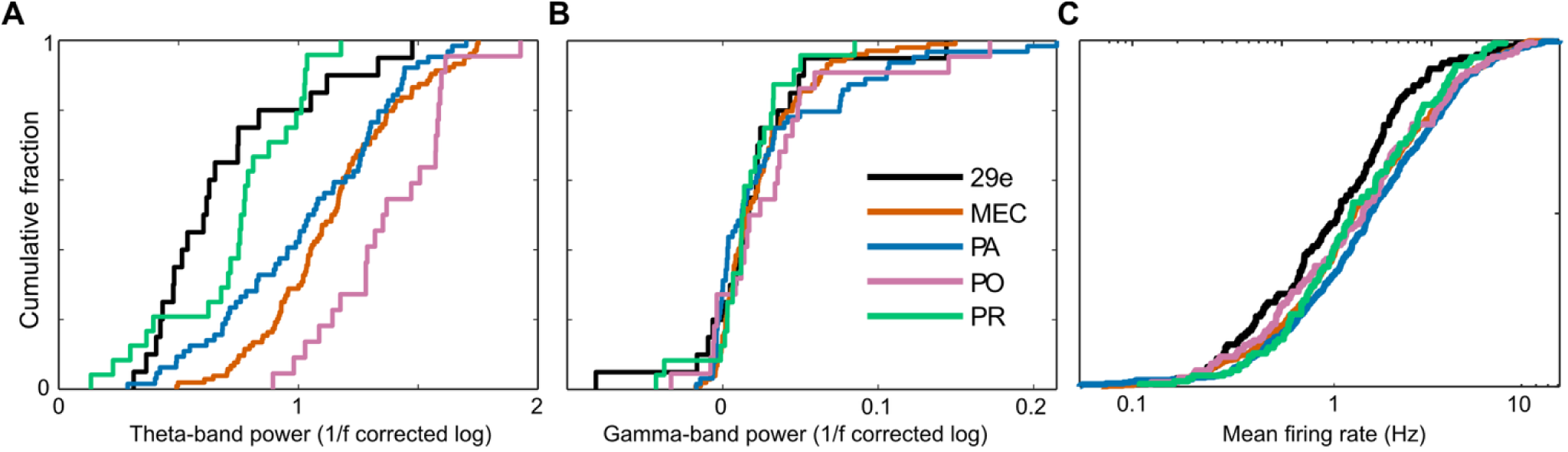
Reduced theta power and firing rates but comparable gamma power in area 29e compared to other areas. **(A)** Cumulative distribution of 1/f-corrected theta-band power across regions, computed at unique tetrode locations (29e: n = 20, MEC: n = 104, PA: n = 64, PO: n =22, PR: n = 24). A linear-mixed effects model including region identity as the fixed effect and two random intercepts (rat identity and region identity within each rat) showed a significant main effect of the region (*F*(4, 9.7) = 4.6, *p =*0.024). Post-hoc testing of pairwise contrasts comparing each region to 29e (Holm-Bonferroni corrected) based on the fitted model determined that theta-band power in 29e was significantly lower than MEC (*t*(10.1) = – 3.07, p = 0.0345 and PO (*t*(14.4) = –3.37, *p* = 0.018) but not PA (*t*(10.9) = –1.86, *p* = 0.179) or PR (*t*(12.2) = –0.154, *p* = 0.880). **(B)** Region-wise cumulative distribution of 1/f-corrected gamma-band power computed at unique tetrode locations. (sample size same as A). A linear mixed-effects model with the same fixed- and random-effects structure as the one in panel A identified main effect of the regional identity as statistically significant (*F*(4, 220.4) = 0.038). When post-hoc pairwise contrasts from the fitted model compared each region against 29e, however, we found that gamma-band power in 29e was comparable to MEC (*t*(204.2) = 0.32, *p* = 0.992), PA (*t*(203.6) = –1.25, *p* = 0.793), PO (*t*(217.3) = –1.29, *p* = 0.793), and PR (*t*(225.3) = –0.68, *p* = 0.992). Thus, although there was a statistically significant difference among recorded parahippocampal regions, it was not driven by a difference between 29e and another region. **(C)** Cumulative distribution of single neuron firing rates across regions. 29e neurons exhibited significantly lower mean firing rates (median [IQR] = 2.34 [0.90, 4.81]) compared to other regions except PO (MEC = 3.03 [1.45, 8.11], PA = 3.86 [1.61, 9.75], PR = 2.90 [1.42, 7.79], PO = 3.07 [0.97, 7.31]; Kruskal-Wallis test χ^2^(4) = 19.4, *p* < 0.001; post-hoc Wilcoxon rank-sum tests with Holm-Bonferroni correction 29e vs. MEC: *Z* = –2.97, *p =* 0.008; 29e vs. PA: *Z =* –4.24, *p <* 0.001; 29e vs. PO: *Z =* –1.61, *p =* 0.105; 29e vs. PR: *Z =* –2.28, *p =* 0.044).

**Fig. S11.**
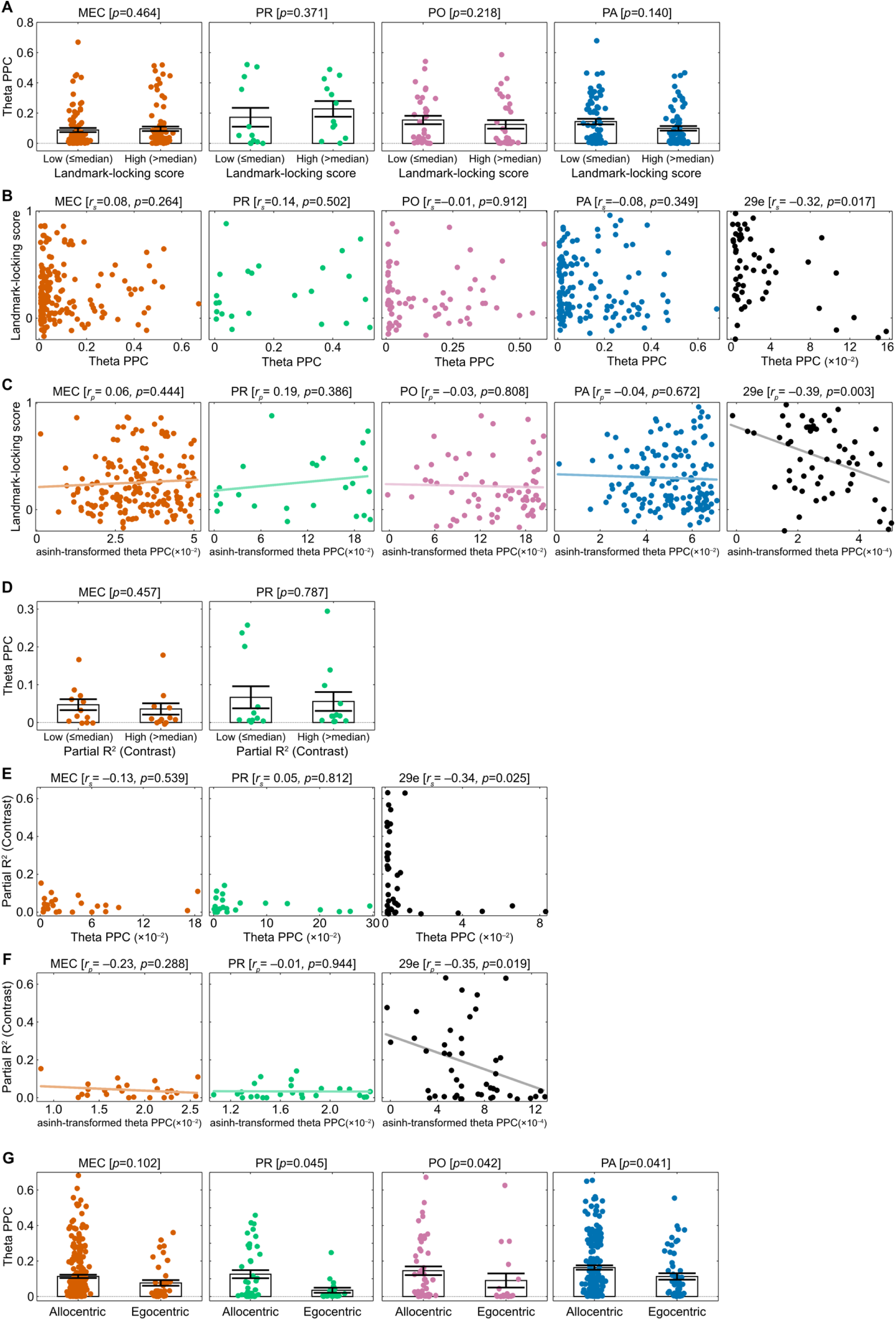
Relationship between single neurons’ spike–LFP theta phase coupling and their functional properties across non-29e regions. **(A)** Theta PPC in neurons with low (≤ median) versus high (> median) landmark-locking scores in non-29e regions during *LM-failure* sessions in the Dome VR. Dots indicate single neurons; bars show mean ± s.e.m.; p-values from Wilcoxon’s rank-sum tests on arcsinh-transformed PPC values are shown above each plot. Unlike area 29e, where landmark-locking scores show considerable heterogeneity (Fig. 2H, right), other regions showed relatively homogeneous (and lower) landmark-locking scores, making their median-split groups sensitive to session-to-session and animal-to-animal variability rather than truly distinguishing neurons based on their landmark anchoring. Additionally, baseline theta PPC differed substantially across regions (Fig. 4F), complicating any cross-regional comparison of the effect. We therefore did not perform a direct statistical comparison between 29e and other regions. Nevertheless, data from other regions are shown here for completeness, with *p*-values displayed above each plot for reference. **(B)** Same data as (A), shown as scatter plots to illustrate the continuous relationship between theta PPC (*x*-axis) and landmark-locking score (*y*-axis) across single neurons from each region recorded in *LM-failure* sessions. Note that, unlike panel A, this panel includes, because the main text and the accompanying main figure (Fig. 4) showed 29e data only in the median-split format as in panel A of this figure. Above each region’s plot, the Spearman’s correlation coefficient (*r_s_*) and the corresponding p-value are given. Only 29e showed a significant correlation, which was negative. **(C)** Same as (B) but with arcsinh-transformed theta PPC on the x-axis to account for the skewed distribution of PPC values, which can take zero or small negative values. The transformation spreads the data more evenly along the *x*-axis by reducing compression at low PPC values, allowing use of Pearson correlation (*r_p_*) instead of Spearman’s (*r_s_*). The negative relationship in 29e remains significant, appearing more uniformly linear across the full range of transformed values, whereas it remains non-significant in other regions. **(D)** Theta PPC in neurons with low (≤ median) versus high (> median) partial R² for contrast tuning in MEC and PR during luminance manipulation sessions in the Dome VR. Conventions as in panel A. As with landmark-locking scores (panel A), contrast tuning was stronger and more heterogeneous in 29e than in other regions, making median-split groups in non-29e regions more sensitive to session-to-session and animal-to-animal variability rather than truly distinguishing neurons based on their contrast tuning. Additionally, baseline theta PPC differed substantially across regions (Fig. 4F), and neuron counts were lower in luminance-manipulation sessions overall. We therefore did not perform a direct statistical comparison between 29e and other regions. Nevertheless, data from other regions are shown here for completeness. **(E)** Same data as (A), shown as scatter plots to illustrate the continuous relationship between theta PPC (x-axis) and partial R² for contrast tuning across single neurons from each region recorded in luminance manipulation sessions. As with panel B, we included Area 29e in this plot because Fig. 4 of the main text showed 29e data only in the median-split format as in panel D. **(F)** Same as (E) but with arcsinh-transformed theta PPC to reduce skewness, which is more suitable for Pearson’s correlation (*r_p_*). The significant negative relationship in 29e persisted after transformation, whereas other regions remained non-significant. **(G)** Theta PPC in egocentric versus allocentric neurons across non-29e regions. Conventions as in panel A. As explained in fig. S9A, non-29e regions showed imbalanced egocentric–allocentric splits (predominantly allocentric), unlike area 29e, where neurons are approximately equally split between egocentric and allocentric reference frames (Fig. 3D). For similar reasons as described there, we did not perform a direct statistical comparison between 29e and other regions. Nevertheless, data from other regions are shown here for completeness, with within-region *p*-values displayed above each plot for reference. Although some regions did not reach statistical significance, all showed a trend in the same direction, with allocentric neurons exhibiting higher theta PPC. than egocentric neurons, suggesting this as a general organizing principle of spatial processing across the parahippocampal cortex.

**Fig. S12.**
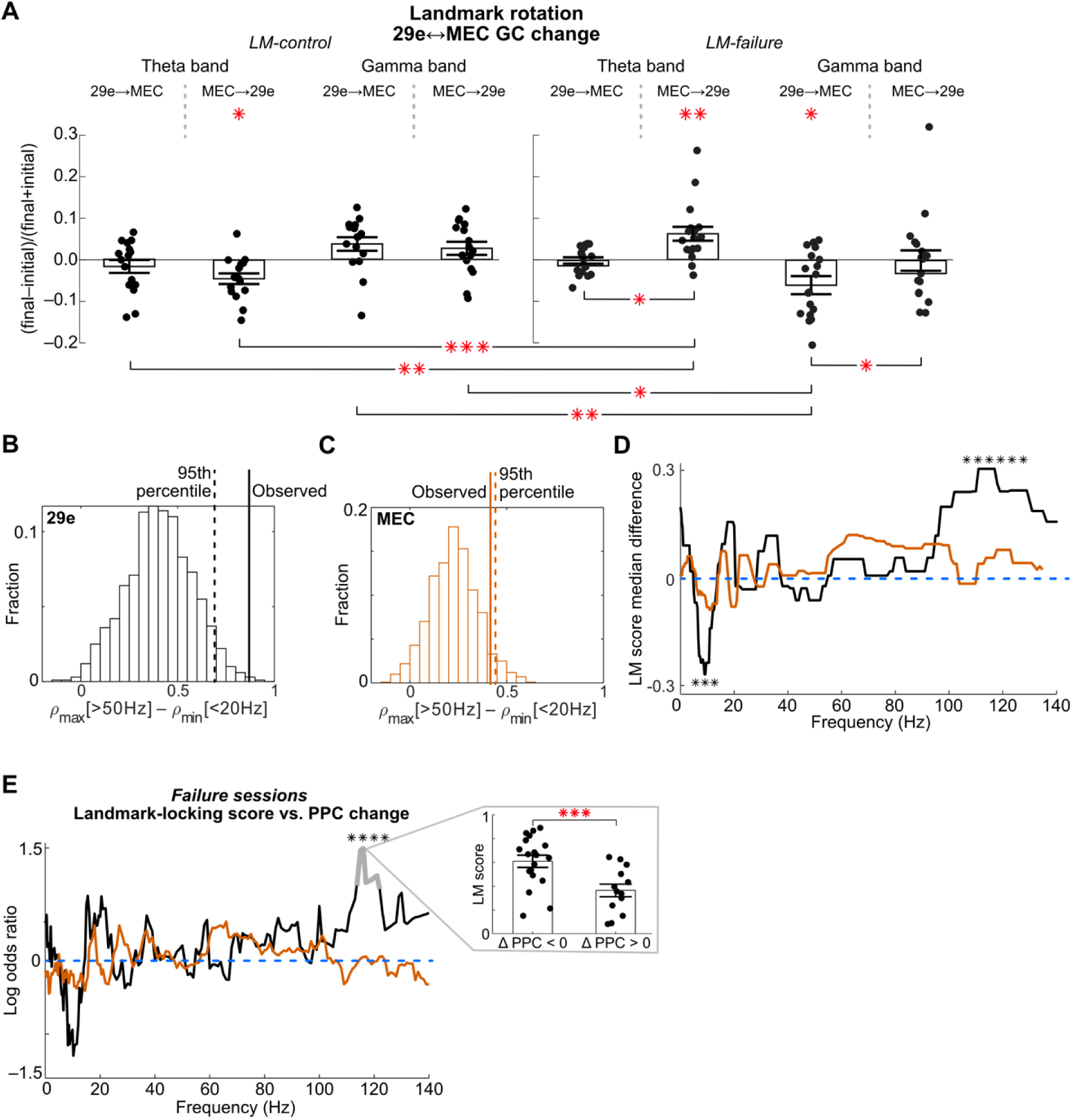
Changes in directed LFP interactions of 29e and MEC tetrode pairs during landmark rotations and the relationship between their cross-regional spike–LFP coupling and landmark anchoring. **(A)** Normalized changes in theta- and gamma-band Granger causality between 29e and MEC LFPs from the initial epoch to the final epoch across *LM-control* and *LM-failure* sessions. Theta band GC showed significant changes in the MEC→29e direction during both *LM-control* (mean ± s.e.m. = –0.05 ± 0.01, Holm-Bonferroni corrected one-sample t-test *t*(15) = –3.2, *p* = 0.022) and *LM-failure* sessions (0.06 ± 0.01, *t(*17) = –3.8, *p* = 0.005); theta-band GC changes in the reciprocal direction (29e→MEC) were not significant during both *LM-control* (–0.018 ± 0.008, *t(15)* = *p* = 0.057) and *LM-failure* sessions (0.003 ± 0.009, *t(*17) = 0.35 *p* = 0.726). For gamma-band GCs, changes were significant in the 29e→MEC direction during *LM*-failure sessions (–0.047 ± 0.014, *t(*17) = –3.3 *p* = 0.013) and marginally significant during *LM-control* sessions (0.041 ± 0.017, *t*(15) = 2.4 *p* = 0.074). Three-way repeated measures ANOVA (factors: session type, frequency band, and direction) identified significant session type × frequency band (*F*(1,32) = 16.2, *p <* 0.001) and significant session type × direction (*F*(1,32) = 25.9, *p* < 0.001) interactions, in addition to a significant main effect of direction (*F*(1,32) = 5.0, *p* = 0.031). Holm-Bonferroni corrected post-hoc pairwise t-tests revealed that, within the theta band, changes in MEC→29e GC during *LM-failure* sessions were significantly higher compared to the changes in [i] MEC→29e GC during *LM-control* sessions (*t(*32) = 4.9, *p* < 0.001), [ii] 29e→MEC GC during *LM-control* sessions (*t(*32) = 4.5, *p* < 0.001), and [iii] 29e→MEC GC during *LM-failure* sessions (*t(*17) = 2.9, *p* = 0.032). Within the gamma band, directionally opposite but statistically similar differences were observed: changes in 29e→MEC gamma-band GC during *LM-failure* sessions were significantly lower compared to the changes in [i] 29e→MEC gamma-band GC during *LM-control* sessions (*t(*32) = –3.9, *p* = 0.002), [ii] MEC→29e gamma-band GC during *LM-control* sessions (*t(*32) = –3.4 *p* = 0.008), and [iii] MEC→29e gamma-band GC during *LM-failure* sessions (*t(*17) = –3.1, *p* = 0.027). **(B-C)** Permutation-based significance testing of the frequency-selective landmark locking–LFP phase coupling pattern for 29e **(B)** and MEC **(C)** neurons. At each permutation, landmark-locking scores from LM-failure sessions were randomly shuffled across 29e or MEC neurons. For each frequency bin (as in Fig. 6C), the correlation across neurons between their baseline PPC values (from the initial stationary-landmark epoch) and the shuffled landmark-locking scores was recalculated, generating a new correlation spectrum for each permutation. From this spectrum, the difference between the maximum correlation above 50 Hz and the minimum correlation below 20 Hz was computed. Repeating this procedure for 1000 permutations yielded null distributions of frequency-based correlation-coefficient differences. The observed difference (solid line) significantly exceeded the 95^th^ percentile of the permutation distribution (dashed line) in 29e (B; *p* = 0.006) but not in MEC (C; *p* = 0.078). **(D-E)** Relationship between landmark locking of 29e and MEC neurons and loss of their cross-regional spike–LFP coupling during LM-failure sessions. (D) Median difference in landmark-locking scores between neurons that lost coupling (”PPC loss”) versus those that gained coupling (”PPC gain”) for 29e (black) and MEC (orange) populations across frequencies. At each frequency, neurons were divided into PPC loss or PPC gain groups, and the median difference in landmark-locking scores between groups was calculated across the population. The statistical significance of the difference between medians at each frequency was assessed with a Wilcoxon signed-rank test. To control for multiple comparisons, a cluster-based permutation test was used in conjunction with this rank test: contiguous frequency bins with uncorrected *p* < 0.05 were defined as clusters, and cluster-level statistics were compared to a null distribution generated by 1000 random permutations of group labels. Asterisks indicate frequency ranges with significant differences after cluster-level, permutation-based correction (*p* < 0.05). Among 29e neurons, those that lost coupling to MEC gamma had significantly higher landmark-locking scores than those that gained coupling, whereas those that gained coupling to MEC theta had significantly lower landmark-locking scores. No such pattern was observed for MEC neurons. (**E**) Logistic regression analysis testing whether landmark-locking score of a neuron predicts its “PPC loss” or “PPC gain” status, after controlling for baseline coupling levels. Because the frequency-dependent pattern among 29e neurons in panel D closely resembled the relationship between their landmark-locking score and their baseline PPC values (Fig. 6C), baseline coupling level may have confounded the observed differences in landmark-locking score between “PPC loss” and “PPC gain” groups by affecting both whether a neuron loses or gains coupling toward a mean level and its degree of landmark anchoring. To control for this potential confound, we performed a logistic regression analysis, using baseline PPC values and landmark-locking scores of each neuron during *LM-failure* sessions as predictors, and the direction of PPC change (loss vs. gain) as the outcome variable. The log odds ratio (β coefficient from logistic regression; *y*-axis) for landmark-locking score as a predictor of PPC loss (ΔPPC < 0) versus gain (ΔPPC > 0) is plotted as a function of frequency (*x*-axis) for 29e neurons’ coupling to MEC LFP (black) and MEC neurons’ coupling to 29e LFP (orange). A log odds ratio of 0 indicates no change in the odds of PPC loss per one standard-deviation increase in landmark-locking score, while a log odds ratio of 1.5 indicates an increase in odds by a factor of exp(1.5) ≈ 4.5 per the same amount of change in landmark-locking score. Asterisks and grey, thickened segments indicate the significant frequency ranges, determined by a cluster-based permutation test as before. Inset shows landmark-locking scores for 29e neurons grouped by whether they lost (ΔPPC < 0) or gained (ΔPPC > 0) coupling at the frequency with largest odds ratio. Overall, the logistic regression revealed that, after accounting for baseline PPC levels, the landmark-locking score remained a significant positive predictor of 29e neuron’s PPC loss to MEC gamma (particularly 100–120 Hz; Fig. 6D). No significant association remained relative to lower-frequency oscillations of MEC LFP (e.g., theta), nor was it observed for MEC neurons’ coupling to 29e LFP across the entire frequency spectrum. Collectively, the results in panels D and E indicate that landmark-anchored 29e neurons lost their coupling to MEC gamma, independent of their initial coupling levels, as landmark control was lost over parahippocampal populations during *LM-failure* sessions.

**Fig. S13.**
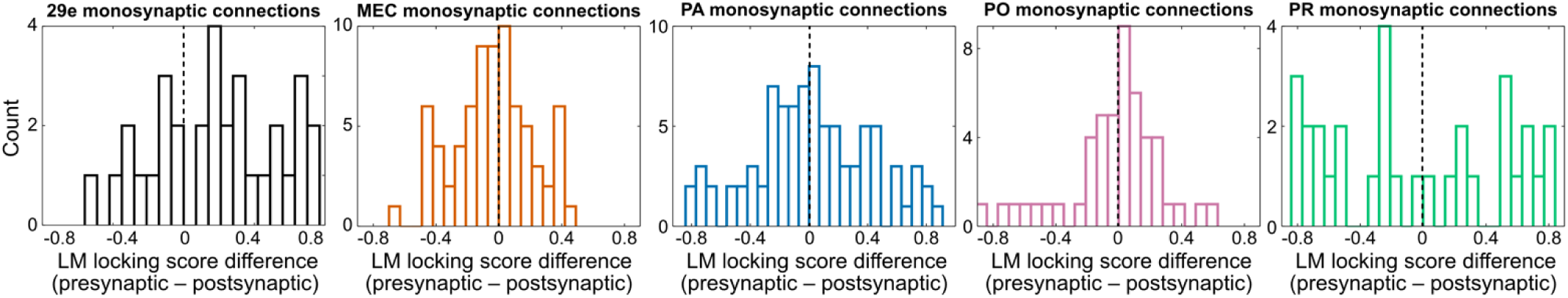
Distributions of landmark-locking score differences between putative pre- and post-synaptic neuron pairs across regions. Putative monosynaptic connections were identified from spike cross-correlograms between simultaneously recorded neurons. Interregional connections were too scarce for a meaningful analysis, so we examined intraregional connections within each region. Each histogram shows the distribution of landmark-locking score differences (presynaptic – postsynaptic) for putative monosynaptic pairs recorded in LM-failure sessions. For each region, we tested whether landmark-locking scores differed between pre- and postsynaptic neurons of putative monosynaptic pairs recorded in LM-failure sessions. Presynaptic neurons exhibited significantly larger landmark-locking scores than their postsynaptic targets only in area 29e (mean ± s.e.m. = 0.19 ± 0.06; Holm-Bonferroni corrected one-sample t-test: *t*(31) = 3.04, *p* = 0.02). In other regions (MEC, PA, PO, and PR), however, landmark-locking scores did not differ significantly between pre- and post-synaptic neurons of putative monosynaptic connections (MEC: mean ± s.e.m. = –0.04 ± 0.03, *t*(73) = –1.34, *p* = 0.724; PA: 0.02 ± 0.04, *t*(85) = 0.57, *p* = 1.000; PO: –0.04 ± 0.04, *t*(48) = 1.00, *p* = 0.957; PR: –0.06 ± 0.12, *t*(30) = 0.48, *p* = 1.000). One-way ANOVA comparing these landmark-locking score differences across regions showed a significant effect of region (*F*(4,267) = 2.59, *p* = 0.038). Post-hoc t-tests with Holm-Bonferroni correction comparing 29e against each other region showed that 29e differed significantly from MEC (*t*(104) = 3.97, *p* < 0.001) and PO (*t*(79) = 3.18, *p* = 0.006), but not PA (*t*(116) = 1.93, *p* = 0.110) or PR (*t*(61) = 1.96, *p* = 0.110).

## Supplementary Tables

**Table S1.** Number of sessions and neurons recorded with 3 versus 4 visual landmarks during closed-loop gain manipulation experiments in the Dome. M = male; F = female. Results were qualitatively similar regardless of whether 3 or 4 landmarks were used.

|  |  | Number of sessions | Number of neurons |
| --- | --- | --- | --- |
| **Rat 994 (M)** | *3 landmarks* | 4 | 58 |
|  | *4 landmarks* | 3 | 62 |
| **Rat 995 (M)** | *3 landmarks* | 3 | 30 |
|  | *4 landmarks* | 7 | 93 |
| **Rat 1041 (M)** | *3 landmarks* | 6 | 238 |
|  | *4 landmarks* | 0 | 0 |
| **Rat 1061 (F)** | *3 landmarks* | 7 | 114 |
|  | *4 landmarks* | 0 | 0 |
| **Rat 1062 (F)** | *3 landmarks* | 1 | 21 |
|  | *4 landmarks* | 1 | 10 |

**Table S2.** **Median values of Dome landmark selectivity metrics and landmark-locking scores in *LM-failure***

|  |  | **Z**<br>**index** | **Local**<br>**similarity**<br>**index** |  | **Landmark**<br>**locking**<br>**score** |
| --- | --- | --- | --- | --- | --- |
| **Rat 994**<br>**(M)** | *29e* (n=2) | 0.101 | −0.169 | *29e* (n=2) | 0.425 |
|  | *MEC* (n=37) | 0.356 | −0.213 | *MEC* (n=32) | 0.062 |
|  | *PA* (n=44) | 0.372 | −0.187 | *PA* (n=25) | 0.063 |
|  | *PO* (n=35) | 0.382 | −0.221 | *PO* (n=22) | 0.055 |
|  | *PR* (n=3) | 0.068 | −0.201 | *PR* (n=2) | 0.355 |
| **Rat 995**<br>**(M)** | *29e* (n=10) | 1.319 | 0.412 | *29e* (n=9) | 0.767 |
|  | *MEC* (n=93) | 0.345 | −0.186 | *MEC* (n=54) | 0.280 |
|  | *PA* (n=12) | 0.297 | −0.238 | *PA* (n=9) | 0.204 |
|  | *PO* (n=3) | 0.188 | −0.240 | *PO* (n=2) | 0.297 |
|  | *PR* (n=6) | 0.283 | −0.149 | *PR* (n=6) | 0.148 |
| **Rat 1041**<br>**(M)** | *29e* (n=19) | 0.625 | −0.025 | *29e* (n=19) | 0.610 |
|  | *MEC* (n=60) | 0.441 | −0.032 | *MEC* (n=57) | 0.296 |
|  | *PA* (n=110) | 0.512 | −0.235 | *PA* (n=97) | 0.253 |
|  | *PO* (n=49) | 0.466 | −0.232 | *PO* (n=41) | 0.197 |
|  | *PR* (n=0) | – | – | *PR* (n=0) | – |
| **Rat 1061**<br>**(F)** | *29e* (n=17) | 0.443 | −0.133 | *29e* (n=3) | 0.542 |
|  | *MEC* (n=67) | 0.295 | −0.284 | *MEC* (n=33) | 0.156 |
|  | *PA* (n=12) | 0.280 | −0.324 | *PA* (n=7) | 0.395 |
|  | *PO* (n=2) | 0.831 | −0.279 | *PO* (n=1) | 0.819 |
|  | *PR* (n=17) | 0.329 | −0.291 | *PR* (n=10) | 0.157 |
| **Rat 1062**<br>**(F)** | *29e* (n=24) | 0.613 | 0.047 | *29e* (n=24) | 0.292 |
|  | *MEC* (n=0) | – | – | *MEC* (n=0) | – |
|  | *PA* (n=0) | – | – | *PA* (n=0) | – |
|  | *PO* (n=0) | – | – | *PO* (n=0) | – |
|  | *PR* (n=7) | 0.496 | −0.323 | *PR* (n=7) | 0.103 |

**Table S3.** Median values of open-arena spatial tuning scores per animal.

|  |  | Skagg's SI<br>score | Rayleigh<br>score | EVF<br>score |
| --- | --- | --- | --- | --- |
| **Rat 994**<br>**(M)** | *29e* (n=7) | 0.016 | 0.179 | 0.343 |
|  | *MEC* (n=148) | 0.031 | 0.084 | 0.231 |
|  | *PA* (n=122) | 0.047 | 0.121 | 0.198 |
|  | *PO* (n=45) | 0.029 | 0.087 | 0.208 |
|  | *PR* (n=15) | 0.043 | 0.363 | 0.160 |
| **Rat 995**<br>**(M)** | *29e* (n=19) | 0.029 | 0.058 | 0.435 |
|  | *MEC* (n=137) | 0.036 | 0.051 | 0.185 |
|  | *PA* (n=16) | 0.029 | 0.182 | 0.298 |
|  | *PO* (n=4) | 0.045 | 0.041 | 0.378 |
|  | *PR* (n=14) | 0.035 | 0.050 | 0.218 |
| **Rat 1041**<br>**(M)** | *29e* (n=20) | 0.035 | 0.491 | 0.516 |
|  | *MEC* (n=79) | 0.045 | 0.041 | 0.136 |
|  | *PA* (n=133) | 0.035 | 0.063 | 0.176 |
|  | *PO* (n=40) | 0.024 | 0.112 | 0.298 |
|  | *PR* (n=0) | – | – | – |
| **Rat 1061**<br>**(F)** | *29e* (n=28) | 0.012 | 0.117 | 0.238 |
|  | *MEC* (n=66) | 0.023 | 0.107 | 0.265 |
|  | *PA* (n=23) | 0.014 | 0.351 | 0.276 |
|  | *PO* (n=3) | 0.028 | 0.303 | 0.381 |
|  | *PR* (n=52) | 0.012 | 0.069 | 0.253 |
| **Rat 1062**<br>**(F)** | *29e* (n=49) | 0.017 | 0.225 | 0.324 |
|  | *MEC* (n=0) | – | – | – |
|  | *PA* (n=0) | – | – | – |
|  | *PO* (n=0) | – | – | – |
|  | *PR* (n=10) | 0.021 | 0.078 | 0.315 |

**Table S4.** Median values of single-cell physiological features.

|  |  | **Theta**<br>**rhythmicity**<br>**index** | **Gamma**<br>**rhythmicity**<br>**index** | **Theta**<br>**PPC** |
| --- | --- | --- | --- | --- |
| **Rat 994**<br>**(M)** | *29e* (n=8) | 1.958 | 1.561 | 0.002 |
|  | *MEC* (n=190) | 8.730 | 0.674 | 0.050 |
|  | *PA* (n=161) | 5.983 | 0.529 | 0.104 |
|  | *PO* (n=58) | 3.892 | 0.926 | 0.038 |
|  | *PR* (n=27) | 8.783 | 0.322 | 0.022 |
| **Rat 995**<br>**(M)** | *29e* (n=12) | 1.168 | 0.840 | 0.0004 |
|  | *MEC* (n=166) | 5.722 | 0.755 | 0.057 |
|  | *PA* (n=27) | 5.259 | 0.529 | 0.071 |
|  | *PO* (n=5) | 2.670 | 1.760 | 0.0006 |
|  | *PR* (n=18) | 4.481 | 0.627 | 0.064 |
| **Rat 1041**<br>**(M)** | *29e* (n=21) | 2.649 | 0.741 | 0.0007 |
|  | *MEC* (n=89) | 2.922 | 0.671 | 0.046 |
|  | *PA* (n=195) | 9.755 | 0.418 | 0.146 |
|  | *PO* (n=67) | 6.281 | 0.730 | 0.097 |
|  | *PR* (n=0) | – | – | – |
| **Rat 1061**<br>**(F)** | *29e* (n=22) | 1.509 | 1.241 | 0.005 |
|  | *MEC* (n=121) | 2.279 | 1.099 | 0.014 |
|  | *PA* (n=31) | 2.309 | 0.555 | 0.023 |
|  | *PO* (n=3) | 3.028 | 0.892 | 0.002 |
|  | *PR* (n=65) | 1.897 | 1.048 | 0.006 |
| **Rat 1062**<br>**(F)** | *29e* (n=60) | 2.133 | 0.943 | 0.0015 |
|  | *MEC* (n=0) | – | – | – |
|  | *PA* (n=0) | – | – | – |
|  | *PO* (n=0) | – | – | – |
|  | *PR* (n=14) | 7.411 | 0.280 | 0.359 |

